# Behavioral phenotyping of an improved mouse model of Phelan-McDermid Syndrome with a complete deletion of the *Shank3* gene

**DOI:** 10.1101/278622

**Authors:** Elodie Drapeau, Mohammed Riad, Yuji Kajiwara, Joseph D. Buxbaum

## Abstract

**Abstract:** Phelan-McDermid Syndrome (PMS) is a rare genetic disorder in which one copy of the *SHANK3* gene is missing or mutated, leading to a global developmental delay, intellectual disability, and autism. Multiple intragenic promoters and alternatively spliced exons are responsible for the formation of numerous isoforms. Many genetically-modified mouse models of PMS have been generated but most disrupt only some of the isoforms. In contrast, the vast majority of known *SHANK3* mutations found in patients involve deletions that disrupt all isoforms. Here, we report the production and thorough behavioral characterization of a new mouse model in which all *Shank3* isoforms are disrupted. Our mice are more severely affected than previously published models. While the deficits were typically more pronounced in homozygotes, an intermediate phenotype was observed for heterozygotes in many paradigms. As in other *Shank3* mouse models, stereotypies, including increased grooming, were observed. Additionally, both sensory and motor deficits were detected in neonatal and adult mice. While social behaviors were not strongly impacted, *Shank3*-deficient mice displayed a strong escape behavior and avoidance of inanimate objects indicating increased novelty-induced anxiety. Similarly, increased freezing was observed during fear conditioning training and amygdala-dependent cued retrieval. Finally, deficits were observed in both initial training and reversal in the Barnes maze and in contextual fear memory that are memory tasks involving hippocampal-prefrontal circuits. This new mouse model of PMS, engineered to most closely represent human mutations, recapitulates core symptoms of PMS providing improvements for both construct and face validity, compared to previous models.

**Significant statement:** Phelan-McDermid syndrome, caused by happloinsufficiency of *Shank3*, is a severe and complex neurodevelopmental disorder. This study investigates the behavioral consequences of a disruption of all *Shank3* isoforms in neonatal and adult mice using a detailed battery of tests tailored to investigate core symptoms and usual comorbidities of PMS. We found that our new model is more severely affected than previously published mouse models with only partial deletions of *Shank3* and more closely recapitulates symptoms of PMS thus providing improvements for both construct and face validity. Our results highlight the significance of using a mouse model with a complete deletion of *Shank3* for studying mechanisms underlying autism spectrum disorder and PMS, carrying preclinical studies and testing test novel therapeutic approaches.

## Introduction

Phelan McDermid syndrome (PMS) is a rare and complex neurodevelopmental disorder that manifests with global developmental delay, mild dysmorphic features, motor deficits, variable degrees of intellectual disability (ID), and absent or delayed speech. Additionally, autism spectrum disorder (ASD), epilepsy, attention deficits and recurrent medical comorbidities are common in patients with PMS (Betancur and Buxbaum, 2013; Phelan and McDermid, 2012; Sarasua et al., 2014a; Soorya et al., 2013). Recent studies show that PMS is emerging as one of the most frequent and penetrant monogenic causes of autism and ID (Betancur and Buxbaum, 2013; Leblond et al., 2014; Soorya et al., 2013; Sykes et al., 2009).

In spite of overlapping etiologies between patients, there is a tremendous heterogeneity in the expression and severity of the phenotype (Cusmano-Ozog et al., 2007; Dhar et al., 2010; Phelan and Betancur, 2011; Soorya et al., 2013). This is no doubt in part due to the complexity in the genetic etiology of PMS. While a large body of data indicates that haploinsufficiency of *SHANK3* is the key contributor for the neurobehavioral manifestations of PMS, it can be caused by a variety of genetic rearrangements including unbalanced translocations, ring chromosome 22, terminal deletions (ranging from deletions of just *SHANK3* to large deletions of up to 9 Mb) and interstitial deletions or point mutation within the *SHANK3* gene (Bonaglia et al., 2011; Durand et al., 2007; Leblond et al., 2014; Moessner et al., 2007; Phelan and McDermid, 2012; Soorya et al., 2013; Sykes et al., 2009).

Genotype-phenotype analyses have shown positive correlations between the size of the deletion and the number and/or severity of some phenotypes (Bonaglia et al., 2011; Dhar et al., 2010; Luciani et al., 2003; Sarasua et al., 2014b; Soorya et al., 2013). However, findings on specific clinical variables have not been consistent across studies. Importantly, it has become clear that indels or point mutations that impact *SHANK3* alone can lead to all of the neurobehavioral phenotypes of PMS. The *SHANK3* gene has multiple promoters and is alternatively spliced and the number of *Shank3* isoforms can be extensive (Benthani et al., 2015; Maunakea et al., 2010). Some de novo microdeletions or mutations of *SHANK3* can therefore affect some but not other *SHANK3* isoforms. The genetic heterogeneity of PMS underscores the importance of studying a wide range of mutations and deletions. SHANK3 (ProSAP2) is a major scaffolding protein that forms a key structural part of the post-synaptic density of excitatory glutamatergic synapses. SHANK3 contains multiple protein-protein interaction domains that each mediates specific protein–protein interactions at synapses. Moreover, the expression and alternative splicing of Shank3 isoforms or even their subcellular distribution has been shown to be cell-type specific, activity-dependent as well as regionally and developmentally regulated (Wang et al., 2014) raising the possibility that differing SHANK3 isoforms may play distinct roles in synaptic developmental and function and hence may make distinct contributions to the pathobiology of PMS.

More than a dozen *isoform-specific Shank3* mouse models have been independently generated (Table 1). As expected, these models shared some similarities but also showed significant differences in molecular, synaptic, and behavioral phenotypes. Depending on the targeted exons, alterations have been reported in motor functions, social interactions, ultrasonic vocalizations, repetitive grooming, cognitive functions and anxiety. However, very high variability has been observed regarding the presence or the intensity of such impairments across several types of *Shank3*-deficient models or even across different cohorts of the same model. These models are based on exonic deletions that have not been reported in human and do not reflect the vast majority of known PMS cases, which are caused by deletions affecting all *SHANK3* isoforms. There was therefore an urgent need to develop an animal model with broader construct validity for PMS to fully understand the consequences of a complete deletion of *SHANK3* across the range of behavioral phenotypes which we achieved through a deletion of exons 4 to 22.

**Table 1:** Summary of existing mouse models of Phelan-McDermid Syndrome. 1-5: Targeted deletions in the ankyrin repeat domain. Δ4-7: deletion of exons 4 to7; Δ4-9: deletion of exons 4 to 9; Δ9: deletion of exon 9. 6: Targeted deletion in the SH3 (Src Homology 3) domain. Δ11: deletion of exon 11. 7-9: Targeted deletions in the PDZ (PSD95/Discs large/zona-occludens-1) domain. Δ13: deletion of exon 13; Δ13-16: deletion of exon 13 to 16). 10-14: Targeted deletions of point mutations in the proline-rich domain. Δ21: deletion of exon 21. 15: Deletion of all functional domains. Δ4-22: deletions of exons 4 to 22. 16: Overexpression of the full Shank3 gene.

|  |  | Strategy | Targeted exons | Domains | Expressed isoforms |  |  |  |  |  |  |  |  |  | Original publication | Other publications | Model name |
| --- | --- | --- | --- | --- | --- | --- | --- | --- | --- | --- | --- | --- | --- | --- | --- | --- | --- |
|  |  |  |  |  | a | a(e10-12s) | b | b(e10-12s) | a/b(e11-12s) | c | d | e | e-1 | f |  |  |  |
| 1 | deletion | Ubiquitous Cre-loxP-mediated excision | exons 4-9 | ankyrin | - | - | - | - | - | + | + | + | + | + | Bozdagi et al, 2010 | Yang et al, 2012; Bozdagi et al, 2013; Drapeau et al, 2014 |  |
| 2 | deletion | Homologous recombination (replacement of exon 4-9 by NEO cassette) | exons 4-9 | ankyrin | - | - | - | - | - | + | + | + | + | + | Wang et al, 2011 | Bariselli et al, 2017 |  |
| 3 | deletion | Ubiquitous Cre-loxP-mediated excision | exons 4-9 | ankyrin | - | - | - | - | - | + | + | + | + | + | Jaramillo et al, 2016 |  |  |
| 4 | deletion | Homologous recombination (replacement of exon 4-7 by NEO cassette) | exons 4-7 | last 3 ankyrin repeats | - | - | - | - | - | + | + | + | + | + | Peça et al, 2011 |  | Shank3A |
| 5 | deletion | Ubiquitous Cre-loxP-mediated excision | exon 9 | last ankyrin repeat | - | - | - | - | - | + | + | + | + | + | Lee et al, 2015 |  | Shank3 (Δ9) |
| 6 | deletion | Homologous recombination (introduction of stop codon in exon 11) | exon 11 | SH3 | - | + | - | + | + | - | + | + | + | + | Schmeisser et al, 2012 | Vicidomini et al, 2017; Reim et al, 2017 | Shank3αβ, Shank3Δ11 |
| 7 | stop codon | Insertion of Neo-Stop cassette in intron 12 | exon 13 | PDZ | - | - | - | - | + | + |  |  |  |  | Jaramillo et al, 2017 |  | Shank3E13 |
| 8 | deletion | Homologous recombination (replacement of exon 13-16 by NEO cassette) | exons 13-16 | PDZ | - | - | - | - | + | - | - | + | + | + | Peça et al, 2011 | Lu et al, 2016; Copping et al, 2017 | Shank3B |
| 9 | inducible deletion | Homologous recombination (inversion of exons 13-16 and flanking with FLEX cassette) + crossing with CAGGS-CreER mice for tamoxifen rescue | exons 13-16 | PDZ | - (+) | - (+) | - (+) | - (+) | + | - (+) | - (+) | + | + | + | Mei et al, 2016 |  | Shank3fx/fx and Shank3fx/fx:CAGGS-CreER |
| 10 | deletion | Ubiquitous Cre-loxP-mediated excision | exon 21 | PRO | - | - | + | + | + | - | - | - | + | - | Bangash et al, 2011 (retracted) | Cope et al, 2016 |  |
| 11 | deletion | Ubiquitous Cre-loxP-mediated excision | exon 21 | PRO | - | - | + | + | + | - | - | - | + | - | Kouser et al, 2013 | Kloth et al, 2015; Duffney et al, 2015; Bidinosti et al, 2016; Li et al, 2017 | Shank3ΔC/ΔC |
| 12 | inducible point insertion | Insertion of a floxed mutated exon 21 followed by a transcriptional stop (Neo-stop) cassette + crossing with B6.Cg-Tg(CAG-cre/Esr1*)5Amc/J for tamoxifen rescue | exon 21 | PRO | - | - | + | + | + | - | - | - | + | - | Speed et al, 2015 |  | Shank3G/G and Reversible-Shank3GCre+ |
| 13 | point insertion | Homologous recombination (G insertion at position 3680 causing a frameshift and premature stop codon) | exon 21 | PRO | - (+) | - (+) | + | + | + | - (+) | - (+) | - (+) | + | - (+) | Zhou et al, 2016 |  | InsG3680 |
| 14 | point mutation | Homologous recombination (R1117X non sense mutation) | exon 21 | PRO | - | - | + | + | + | - | - | - | + | - | Zhou et al, 2016 |  | R1117X |
| 15 | deletion | Ubiquitous Cre-loxP-mediated excision | exons 4-22 | ANK, SH3, PDZ, PRO, SAM | - | - | - | - | - | - | - | - | - | - | Wang et al, 2016 | Han et al, 2016 | Shank3Δe4-22 |
| 16 | overexpression | EGFP-Shank3 BAC transgenic mice | full gene |  | ++ | ++ | ++ | ++ | ++ | ++ | ++ | ++ | ++ | ++ | Han et al, 2013 |  |  |

Interestingly, as our work was progressing, a completely independent mouse model, similarly targeting exons 4 to 22, was reported (Wang et al., 2016b). These mice highlight cortico-striatal circuit abnormalities and demonstrate a behavioral phenotype that resemble features of PMS. We therefore decided to conduct a comprehensive and behavioral evaluation of our mouse model evaluating many more phenotypes relevant to PMS and ASD. Critically, our findings complement and supplement the observations made by the Jiang group with many results clearly confirmed across two independent laboratories as well as unique analyses in each study.

## Materials and Methods

### Generation of inbred strains of *Shank*^*Δ4-22*^ -deficient animals

All animal procedures were approved by the Institutional Animal Care and Use Committee of the [author’s institution]. A *Shank3^Δ4-22^* mouse line with a complete disruption of the *Shank3* gene was generated at Ozgene (Perth, Australia) by retargeting Bruce4 C57BL/6 embryonic stem cells from a previously published mouse. A third loxP site was inserted immediately downstream of exon 22 in addition of the 2 pre-existing loxP sites flanking exons 4 and 9 (Figure 1A). To generate the mice used in the present study, the floxed allele was excised by breeding with a ubiquitous Cre transgenic line resulting in a deletion of exons 4 to 22 and therefore a constitutive disruption of all the Shank3 murine isoforms.

**Figure 1:**
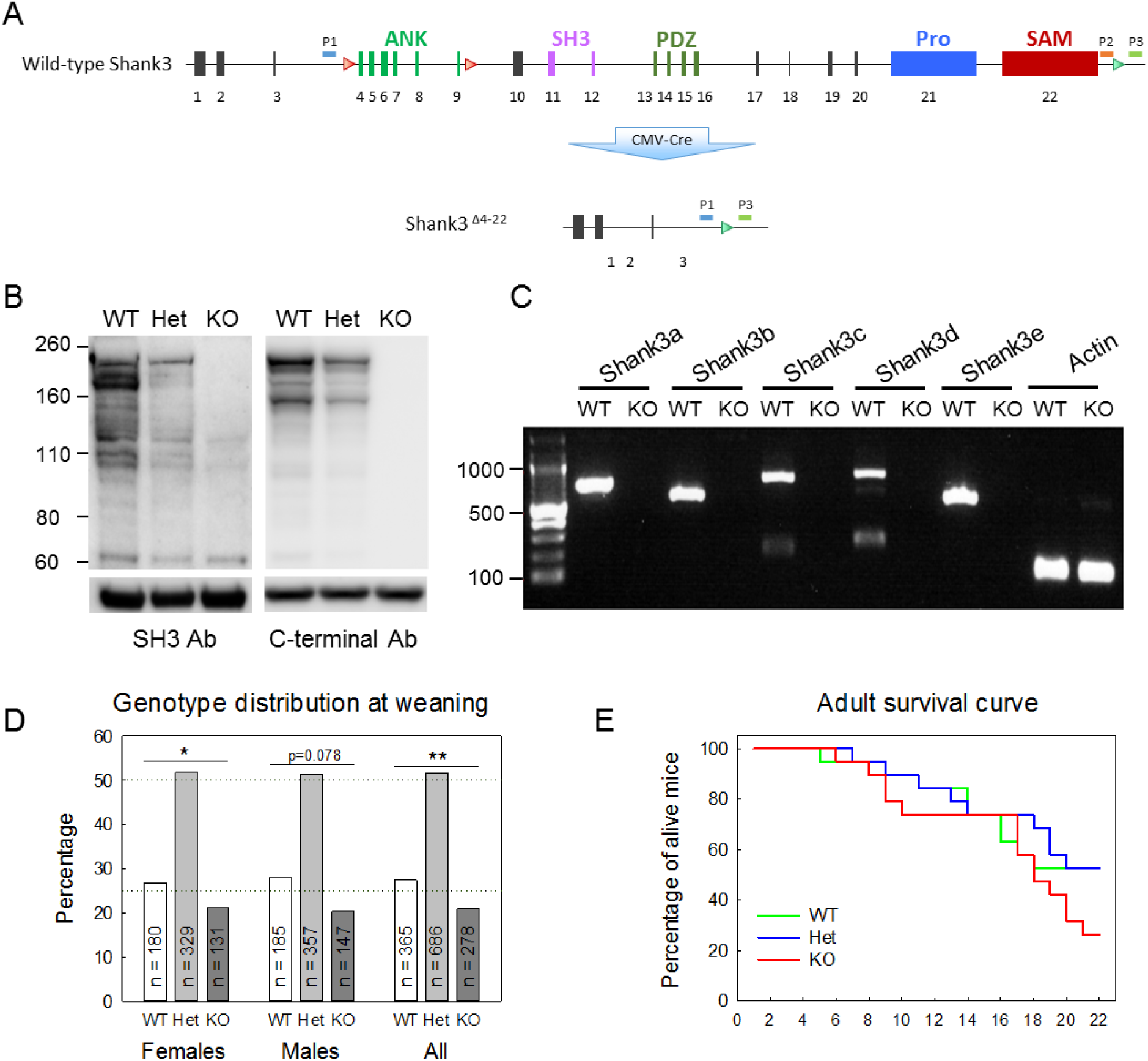
Generation and validation of a knockout mice with a complete deletion of Shank3. **(A)** Schematic design for generation of a *Shank3^Δ4-22^* complete knockout mouse using a Cre-loxP strategy. Bruce4 C57BL/6 embryonic stem cells from a previously generated mouse with two LoxP site located upstream exon 4 and downstream exon 9 (top, red triangles) were retargeted to insert an additional LoxP site 155 pb downstream of exon 22 (green triangle). Floxed mice were crossed with CMV-Cre mice to generate ubiquitous deletion of exons 4 to 22 (bottom). ANK, ankyrin repeats; SH3, Src homology 3 domain; PDZ, postsynaptic density protein, Pro, proline-rich domain; SAM, sterile a-motif domain. The positions of the PCR primers (P1, P2, P3) for genotyping are indicated. **(B)** Expression of Shank3 in postsynaptic density (PSD) fractions. PSD fractions from wild-type, heterozygous and homozygous mice were subjected to immunoblotting with either the N367/62 anti-Shank3 antibody directed against an epitope in the SH3 domain or the H160 C-terminal antibody. Immunoblots show that all Shank3 protein bands are absent in KO brains. The migration of molecular weight markers is shown on the left (in kilodaltons) and an immunoblot for βIII-tubulin as a loading control is shown below. Original full scans of immunoblots are displayed in Extended Figure 1-1. **(C)** RT-PCR analysis for specific Shank3 transcripts in *Shank3^Δ4-22^* mice. Brain-derived mRNAs from wild-type and homozygous mice were subjected to RT-PCR targeting different isoforms. All transcripts were absent in *Shank3^Δ4-22^* homozygous mice.

The colony was maintained on a pure C57BL/6Tac background (Taconic, Germantown, New York, USA). Heterozygous mice were mated to generate litters consisting of three genotypes, wild-type (WT), heterozygote (Het), and knock-out (KO). Mice were weaned at 21 days of age, and at least one littermate from each genotype were group housed in standard plastic cages of three to five littermates per cage. Standard rodent chow and tap water were available ad libitum. The colony room was maintained on a 12-hour light/dark cycle with lights on at 06:00 at a constant temperature of 21-22°C and 55% humidity. All animal procedures were performed in accordance with the [Author University] animal care committee’s regulations

### Genotyping

The confirmation of the deletions of all Shank3 isoforms was performed by RT-PCR. All the animals included in this study were genotyped using tail samples collected at the time of weaning. Additionally, the genotype of all the adult animals was confirmed using a supplementary biopsy at the end of the behavioral testing. Mouse tail snips were collected by dissecting 0.2 cm of tail between postnatal days 15 and 21. Tails were digested, genomic DNA isolated and purified using the Qiagen DNAeasy kit (Qiagen, Valencia, CA, USA) according to the manufacturer’s instructions. After the extraction, 2.0 µl of DNA in buffer containing ∼250–400 µg of DNA was amplified by PCR using standard PCR methods and a combination of three primers designed inside and outside the deleted region to identify both the wild-type and Δe4-22 alleles (Figure 1 and Extended Figure 1-1; P1-KO: TGAGACCAGAGTTGTTAGGATTTG, P2-WT: AGATGGCTCAGCCAGGTAAG, P3-Common AGATGGCTCAGCCAGGTAAG). The P1-P3 primer pair produced a 490 bp band identifying the Δe4-22 allele, while the P2-P3 primer pair amplified a 390 bp band from the wild-type allele. Denaturing, annealing, and extension steps were performed using 94 °C for 3 min, 35 cycles of 94 °C for 30 s, 62 °C for 45 s, 45 °C for 30 s, and for 1 cycle 72 °C for 4 min. The PCR products were run on a 1.5% agarose gel and stained with ethidium bromide.

### Immunoblotting

PSD fractions were prepared as follows. Hemibrains of wild-type, heterozygous and homozygous *Shank3^Δ4-22^* mice were homogenized in 2-[4-(2-hydroxyethyl)piperazin-1-yl]ethanesulfonic acid (HEPES)-A containing 4 mM HEPES, pH 7.4, 0.32 M sucrose, Protease Inhibitor Cocktail and PhoSTOP Phosphatase Inhibitor Cocktail (both from Roche, Indianapolis, IN, USA). Nuclear fractions were precipitated by centrifuging twice at 700 g for 15 min, and the resulting supernatants were further centrifuged at 21,000 g for 15 min. The precipitates were resuspended in HEPES-B containing 4 mM HEPES, pH 7.4, Protease Inhibitor Cocktail and PhoSTOP Phosphatase Inhibitor Cocktail, homogenized and rotated at 4°C for 1 hour. The lysates were centrifuged at 32,000 g for 20 min and washed twice with HEPES-C containing 50 mM HEPES, pH 7.4, 0.5% Triton X-100, Protease Inhibitor Cocktail and PhoSTOP Phosphatase Inhibitor Cocktail. Finally, postsynaptic density fractions were resuspended in HEPES-C containing 1.8% sodium dodecyl sulfate (SDS) and 2.5 M urea. Fifty micrograms of PSD fraction was loaded to 4-12% SDS-polyacrylamide gel electrophoresis (PAGE gel, Invitrogen, Carlsbad, CA, USA), transferred to polyvinylidene fluoride membrane and immunoblotted with either the N367/62 anti-Shank3 antibody directed against an epitope in the SH3 domain (UC Davis/NIH NeuroMab Facility, Davis, CA) or the H160 anti-Shank3 antibody directed against amino acids 1431-1590 mapping near the C-terminus of isoform 2 of Shank 3 (sc-30193, SantaCruz Biotech, Dallas, TX, USA). For βIII tubulin, the membrane was stripped and immunoblotted with an anti-βIII tubulin antibody (Abcam, Cambridge, MA, USA).

### RT-PCR isoform analysis

Total RNA from hemibrains of wild-type and homozygous *Shank3^Δ4-22^* mice was isolated using the TRIzol method (Invitrogen, ThermoFisher Scientific, CA, USA). Reverse transcription was performed with SuperScript^®^ III first-strand synthesis system (Invitrogen, ThermoFisher Scientific, CA, USA). DNA was amplified by PCR using standard PCR methods and the following primers ass described in (Wang et al., 2014). Shank3a Forward: ACGAAGTGCCTGCGTCTGGAC, Shank3a Reverse: CTCTTGCCAACCATTCTCATCAGTG, Shank3b Forward: GTAGCCACCTCTTGCTCACAT, Shank3b Reverse: TTGCCAACCATTCTCATCAGT, Shank3c Forward: CTTCTTCACTGGCAATCCTTG, Shank3c Reverse: CAGTGTAGTGGCGGAAGAGAC, Shank3d Forward: AGGGTCACGACTGTTTCTTAGC, Shank3d Reverse: TGTGGGTGTAAACTCCTCAATG, AACTGCCAGGATCTCATCCA. Shank3e Forward: GTACCTGGGTCTGGGTGCTTTA, Shank3e Reverse: AACTGCCAGGATCTCATCCA.

### Behavioral overview

Three cohorts were used for behavioral testing. The first cohort consisted of 54 newborn mice (14 WT, 30 Het and 10 KO) from 10 independent litters. The second cohort consisted of 57 newborn mice (16 WT, 32 Het and 9 KO) from 9 independent litters. Cohorts 3 (30 adult male mice, 11 WT, 10 Het and 9 KO) and 4 (27 adult male mice, 11 WT, 10 Het and 9 KO) were tested between 3 and 10 months of age according to the schedule described in Table 3. In each adult cohort, all mice were born within two weeks of each other, and generally only one triplet came from any given individual litter of mice. Behavioral experiments were conducted between 9:00 and 17:00 during the light phase of the 12:12 h light/dark cycle in dedicated testing sound-attenuated rooms. Mice were brought to the front room of the testing area at least half an hour prior to the start of experiments. All three genotypes were tested on the same day in randomized order by two investigators who were blind to the genotypes. Behavioral tests were conducted in the order and at the ages indicated in Table 3 and included developmental milestones, cage observation, neurological and motor reflexes, open field, elevated zero maze, Y-maze, beam walking, grip strength, gait analysis, rotarod, 3-chambered social interaction task, nest building, novel object recognition, fear conditioning, pre-pulse inhibition, tail flick, olfactory habituation/dishabituation, buried food, social transmission of food preference, marble burying, 4-object repetitive novel object contact task, male-female social interaction, and Barnes maze. Behavioral results are not described in the order they were tested in an effort to ease presentation and interpretation of the data.

### Newborn development

The physical, sensory and motor developmental milestones of neonates were assessed between postnatal days 1 and 21 using a battery of tests adapted from the Fox scale (Fox, 1965; Heyser, 2004). As we had previously observed a higher rate of postnatal mortality on the first litter, only dams that already had one litter were used for this experiment. To control for litter and avoid nutritional effects the litter size was homogenized and limited to 6 pups per dam by reducing larger litters and adding excess pups to smaller litters on the morning of postnatal day 1 where and when possible. At this time, pups were identified by paw tattoo using a non-toxic animal tattoo ink (Animal Identification & Marking Systems Inc, Piscataway, NY, USA) inserted subcutaneously through a 30-gauge hypodermic needle tip into the center of the paw. Individual pups were removed from the litter and placed on cotton pads in a heated cage under a heating lamp throughout the testing. Each subject was tested at approximately the same time of day. For all the timed tests, a 30-seconds cut-off was used and nonresponding animal received a score of 30 seconds. Most responses were considered positive only after they had been observed for 2 consecutive days.

The physical development was measured by following the weight (P1 to P21), eye opening (P9 to P20), tooth eruption (P7 to P18) the ear development (P1 to P9) and the fur development (P1 to P14) using the following scales. Eye opening, per eye: 0 = eye fully closed, 1 = eye partially opened, 2 = eye full opened, tooth eruption, scored separately for bottom and top incisors: 0 = incisors not visible, 1 = incisors visible but not erupted, 2 = incisors fully erupted. Ear development, per ear: 0 = ear bud not detached from the pinna, 1 = ear flap detached from the pinna, ear fully developed on the back of the ear). Fur development: 1 = bright red, 2 = nude, pink, 3 = nude, grey, 4 = grey, fuzzy on back and shoulder, 5 = black hair on back, grey fuzzy belly, 6 = body fully covered.

Sensory development was assessed using cliff aversion (P2 to P14), auditory startle (P6 to P18), rooting reflex (P2 to P10), ear twitch (P7 to P15) and forelimb grasp (P5 to P14) using the following measures. For cliff aversion, the subject was placed on the edge of a plexiglass platform with a 30-cm cliff with its nose and forefeet over the edge. The latency to move away from the edge was recorded. Auditory startle was measured in response to an 80 dB click 30 cm above the mouse and was considered present when the pup moved immediately after the presentation of the auditory stimulus. For the rooting reflex, the side of the pup’s face were bilaterally stimulated with two cotton swabs. The reflex was considered present when the pup crawled forwards pushing the head during the stimulation. For the ear twitch, the ear of the pup was stimulated with the tip of a cotton swab that was previously pulled to form a filament. Both ears were successively stimulated and the test was considered positive when the pup turned its head or jumped in response to the stimulation. The forelimb reflex was tested by gently stimulated the front paws with the loop of a small bended metallic wire. Each front paw was scored separately as follow: 0 = no response to stimulation. 1 = paw folding in response to the stimulation, 2 = paw grasping the wire in response to the stimulation, 3 = grasp strong enough to hold for at least one second when the wire was lifted up.

Motor development was studied using surface righting (P2 to P13), negative geotaxis (P2 to P14), air righting (P8 to P20), open field crossing (P8 to P20) and rod suspension (P11 to P20) using the following criteria. The surface righting was measured by the time for pups placed on their back to fully turn with all four paws on the ground. For negative geotaxis, pups were placed head down on a mesh covered plan that was slanted at a 45° angle and the latency to either roll down, stay or turn and move up the slope was recorded. For the air righting, the pup was dropped upside down at a height of 30 cm over a padded surface. Subjects received a score of 2 if they successfully righted themselves during the fall, 1 if they landed on the side and 0 if they did no turn. The open field crossing was measured by the time to exit a 13-cm diameter circle when place on the center of the circle. For the rod suspension, the pups were gently grabbed by the trunk, brought up close to a 3-mm wooden rod 30 cm above a padded surface and released once they grabbed the rod with their front paws. The latency to stay suspended was recorded.

### Physical factors, gross appearance and spontaneous activity

Adult animals were handled daily for one week before starting behavioral testing and general health, weight (grams), length (centimeters), physical factors, gross appearance, and spontaneous activity were recording during handling using the following scales.

*Physical factor and gross appearance.* Coat appearance: 0 = ungroomed, 1 = partially groomed, 2 = semi-groomed, 3 = groomed. Skin color (pinna and footpads): 0 = pink, 1= purple, 2 = other. Whisker barbering: 0 = normal, 1 = abnormally shortened. Patches of missing fur on face or body: 0 = none, 1 = some, 2 = extensive. Wounding: 0 = none, 1 = signs of previous wounding, 2 = slight wounds present, 3 = moderate wounds present, 4 = extensive wounds present. Body tone when both sides of the mouse are compressed between thumb and index finger: 0 = flaccid, no return of cavity to normal, 1 = slight resistance, 2 = extreme resistance. Palpebral closure: 0 = eyes wide open, 1 = eyes half open, 2 = eyes closed. Spontaneous piloerection: 0 = none, 1 - coat standing on end.

*Spontaneous general activity* in a 1000 mL jar and after transfer in a regular home cage for five minutes each. Body position: 0 = completely flat, 1 = lying on side, 2 = lying prone, 3 = sitting or standing, 4 = rearing on hind legs, 5 = Repeated vertical leaping. Spontaneous activity: 0 = none, resting, 1 = casual scratch, groom, slow movement, 2 = vigorous scratch, groom, moderate movement, 3 = vigorous, rapid/dart movement, 4 = extremely vigorous, rapid/dart movement. Respiration rate: 0 = gasping, irregular, 1 = slow, shallow, 2 = normal, 3 = hyperventilation. Tremor: 0 = none, 1 = mild, 2 = marked. Urination: 0 = none, 1 = little, 3 = moderate amount, 4 = extensive. Defecation: number of fecal boli. Transfer arousal: 0 = coma, 1 = prolonged freeze, then slight movement, 2 = brief freeze, then active movement, 3 = no freeze, stretch attends, 4 = no freeze, immediate movement (manic). Gait: 0 = normal, 1 = fluid but abnormal, 2 = slow and halting, 3 = limited movement only, 4 = incapacity. Pelvic Elevation: 0 = markedly flattened, 1 = barely touches, 2 = normal (3mm elevation), 3 = elevated (more than 3mm elevation). Tail Elevation: 0 = dragging, 1 = horizontally extended, 2 = less than 30° elevation, 3 = 30° − 60° elevation, 4 = 60° − 90° elevation.

### Motor testing

#### Gait analysis

Motor coordination and gait patterns was observed as the subject was allowed to run the length of an elevated runway (dimensions 152 cm long × 10 cm weight) lined with white paper (Carter et al., 2001). After three 3 training runs, the subject’s paws were coated in non-toxic paint (different colors for hind and front paws) to record paw prints on two consecutive runs. The record displaying the clearest prints and most consistent gait for analysis of 50 cm was chosen to measure sway (mean distance between left and right paws), stride (mean distance between same side front and hind paws) and diagonal stance (mean distance between diagonally opposed front and hind paws).

#### Open field

Mice were tested in an open field (45 × 45 cm) virtually divided into central and peripheral regions. Animal activity was recorded by video tracking (Noldus Ethovision, Leesburg, VA). Each mouse was allowed to explore the apparatus for 60 minutes. The distance travelled, the number of rears and revolutions, the number of grooming bouts and cumulative grooming time, the number of head shaking or twitches, the number of entries in the center and the time spent in the central and peripheral regions were recorded. Measures were recorded in 10-minute intervals.

#### Rotarod

Motor coordination, endurance and learning was assessed in the Rotarod test (Omnitech Electronics Inc, Columbus, OH, USA). Mice were placed on an elevated accelerating rod (3 cm diameter) for three trials per day on two consecutive days. Each trial lasted for a maximum of 5 minutes, during which the Rotarod underwent a linear acceleration from 4 to 40 rpm. A 20-minute interval was used between trials to avoid fatigue. Animals were scored for their latency to fall.

#### Beam walking

Subtle deficits in fine motor coordination and balance that might not be detected by other motor tests were assessed by the beam walking assay in which the mouse had to walk across an elevated horizontal wood beam (100 cm long, 1 m above bedding) to a safe dark box (Carter et al., 2001). Subjects were placed near one end in bright light, while the far end with the dark box was placed in darkness, providing motivation to cross. Performance was quantified by measuring the latency to start crossing, the time to reach the dark box or the time to fall, the total distance traveled and the number of paw slips or incomplete falls (mice able to climb back on the rod). Animals were successively trained on three different beams: 1 inch, ½ inch and ¼ inch diameter and scored on four consecutive trials per beam with one minute of rest between trials on the same beam and 20 to 30 min between each beam. Mice that did not reach the box after 2 minutes were gently placed inside the box and allowed to stay inside for one minute.

#### Righting Reflex

The subject was grasped by the nape of the neck and base of the tail, inverted so back faced down, and released 30 cm above subject’s home cage floor. Righting ability was scored as follow: 0 = no impairment, 1 = lands on side, 2 = lands on back, 3 = fails to right even when placed on back on the floor.

#### Hind limb placing

Subject was lowered by the base of the tail until it grasped a horizontal wire grid with both forepaws. The grid was rotated to vertical and the tail was released. Mice were evaluated over three trials, three minutes apart for their latency to fall or latency to pull body on the grid and the ability to place hind paws was scored as follow: 0 = grabs but falls, 1 = grabs but hangs, 3 = grabs and pulls body onto grid. Maximum cut-off was 60 seconds.

#### Hanging

The subject, held from the base of the tail, was allowed to grasp a wooden rod with both forepaws, rotated to horizontal and release. Test was repeated three times with a three-minute interval between trials and a 60-second maximum cut-off. Both the latency to fall and overall performance scored as follow were recorded: 0 = does not grasp, 1 = grasps but falls immediately, 2 = grasps but then falls off, 3 = grasps and stays on for 60 seconds.

#### Negative Geotaxis

The subject was placed on a wire mesh grid and the grid was lift vertically, with subject facing down. Test was repeated three times with a three-minute interval between trials and a 60-second maximum cut-off. Both the latency to fall and overall performance scored as follow were recorded: 0 = falls off, 1 = does not move, 2 = moves but does not turn, 3 = turns but does not climb, 4 - turns and climbs up.

#### Inverted screen

The subject was placed on a grid screen. The grid was waved lightly in the air, then inverted 60 cm over a cage with soft bedding material. Mice were tested only one time with a 60-second maximum cut-off and the latency to fall was recorded.

#### Grip strength

Forelimb muscle strength and function was evaluated with a strength meter (Ametek, Largo, FL, USA). This test relies on the instinctive tendency of mice to grasp an object with their forelimbs. The animal was pulled backward gently by the tail, while grasping a pull bar connected to a tension meter and the force at the moment when the mouse lost its grip was recorded as the peak tension. Test was repeated three times with a three-minute interval between trials. Each trial consisted in five attempts in quick successions for which the best value was recorded therefore increasing the chances that the measure will accurately reflect maximum strength. The mean of three trials and the largest value from all trials were used as parameters.

### Sensory testing

#### Sensory reflexes

Sensory abilities were evaluated through the reflex response to several sensory modalities using the following scales. Pinna reflex in response to a gentle touch of the auditory meatus with a cotton-tipped applicator repeated three times with a 10 to 15-second interval: 0 = none, 1 = active retraction, moderately brisk flick, 2 = hyperactive, repetitive flick. Corneal Reflex in response to a gentle puff of air repeated three times with a 10 to 15 seconds interval: 0 = no eye blink, 1 = active eye blink, 2 = multiple eye blink. Toe pinch normal retraction reflexes in all four limbs when lightly pinching each paw successively by applying a gentle lateral compression with fine forceps while the mouse is lifted by its tail so the hind limbs are clear of the table. Score is cumulative of four limbs: 0 = no retraction, 1 = active retraction, 2 = repetitive retractions. Preyer reflex in response to a 90 dB click 30 cm above mouse repeated three times with a 10 to 15-second interval: 0 = None, 1 = Preyer reflex (head twitch), 2 = jump less than 1 cm, 3 = Jump more than 1 cm.

#### Tail flick test

The automated Tail-Flick test (Omnitech Electronics Inc, Columbus, OH, USA) was used to assess nociceptive threshold. Awake mice were placed in a contention tube to limit movement with their tail resting on the groove of a heating panel. When the mice were calm, a narrow heat producing beam was directed at a small discrete spot about 15 mm from the tip of the tail. When the subject’s tail was removed from the beam, an automatic timer recorded the latency. The test was repeated five times with a three-minute interval between each trial. The latency of the mice to flick their tail was recorded and the two trials with the shorter latencies were discarded since the tail is not always fully in the beam and this is often an outlier.

#### Acoustic Startle Response and Pre-Pulse Inhibition of Startle

Subjects were placed in isolation boxes outfitted with accelerometers to measure magnitude of subject movement (Med Associates, Fairfax, VT, USA). After five minutes of acclimation mice were first tested for acoustic startle response. Mice were presented with six discrete blocks of six trials over 8 minutes, for a total of thirty-six trials. The trials consisted in six responses to no stimulus (baseline movement), six responses to 40 ms sound bursts of 74 dB, six responses to 40 ms sound bursts of 78 dB, six responses to 82 ms sound bursts of 100 dB, 5 responses to 40 ms sound bursts of 86 dB and six responses to 40 ms sound bursts of 92 dB. The six trials type were presented in pseudorandom order such that each trial type was presented once within a block of six trials. Mice were then tested for pre-pulse inhibition of startle. They were presented with seven discrete blocks of trials of six trials over 10.5 min for a total of forty-two trials. The trials consisted in six response to no stimulus (baseline movement), six startle response to a 40 ms, 110 dB sound burst, six prepulse inhibition trials where the 110 dB tone was preceded by a 20 ms 74 dB tone 100 ms earlier, six prepulse inhibition trials where the 110 dB tone was preceded by a 20 ms 78 dB tone 100 ms earlier, six prepulse inhibition trials where the 110 dB tone was preceded by a 20 ms 82 dB tone 100 ms earlier, six prepulse inhibition trials where the 110 dB tone was preceded by a 20 ms 86 dB tone 100 ms earlier and six prepulse inhibition trials where the 110 dB tone was preceded by a 20 ms 92 dB tone 100 ms earlier. The seven trial types were presented in pseudorandom order such that each trial type was presented once within a block of seven trials. Startle amplitude was measured every 1 ms over a 65 ms period, beginning at the onset of the startle stimulus. The inter-trial interval was 10 to 20 seconds. The maximum startle amplitude over this sampling period was taken as the dependent variable. A background noise level of 70 dB was maintained over the duration of the test session.

#### Visual acuity

Visual acuity was tested using the visual placing test that takes advantage of the forepaw-reaching reflex: the mouse was held by its tail about 20 cm above the surface and progressively lowered. As it approaches the surface, the mouse should expand its forepaws to reach the floor. The test was repeated three times with a 30-second interval and the forepaw reaching reflex was quantified as the percentage of forepaw-reaching episodes that did not involve the vibrissae and/or nose touching the surface before the forepaws.

#### Buried Food Test

The buried food test (Yang and Crawley, 2009) measures how quickly an overnight-fasted animal can find a small piece of familiar palatable food, that is hidden underneath a layer of bedding using olfactory clues. Fruit Loops (Kellog’s, Batle Creek, MI, USA) were used as familiar food. For three consecutive days before the test, 3-4 pieces were offered to the subjects to make sure it was highly palatable for all the subjects. 18 to 24 hours before the test, all chow pellets were removed from the subjects’ home cages. The water bottle was not removed. On the testing day, the subject was placed in a clean cage (28 cm L × 18 cm W × 12 cm H) containing 3 cm deep of clean bedding and the subject was allowed to acclimate to the cage for ten minutes. While the subject was temporary placed in an empty clean cage, 4-5 pieces of Fruit Loops were buried approximately 1 cm beneath the surface of the bedding, in a random corner of the cage and the bedding surface was smoothed out. The subject was placed back in the testing cage and given fifteen minutes to retrieve and eat the hidden food. Latency to find the food was recorded. If a subject did not find the food, fifteen minutes was recorded as its latency score and the food was unburied and presented to the mouse by the experimenter to make sure that it was palatable for the mouse. At the end of testing, subjects were hold in a temporary cage until all animals from the same home cage were tested.

#### Olfactory habituation and dishabituation

This test consisted of sequential presentations of different non-social and social odors in the following order: water, lemon extract (McCormick, Hunt Valley, MD; 1:100 dilution), banana extract (McCormick, Hunt Valley, MD; 1:100 dilution), unfamiliar males and unfamiliar females (Yang and Crawley, 2009). Lemon and banana solutions were freshly prepared everyday using distilled water. Social odors were obtained from cages of unfamiliar C56BL/6 mice of the same and opposite sex as the subject which have not been changed for at least three days and were maintained outside of the experimental testing room. Social odor stimuli were prepared by wiping a cotton swab in a zigzag motion across the cage. The subject was placed in a clean bedding-covered testing cage covered with the cage grid. A clean dry applicator (10 cm cotton swab) was inserted through the cage grid water bottle hole and the animal was allowed to acclimate for 30 minutes to reduce novelty-induced exploratory activity during the olfaction test. Each odor (or water) was presented in three consecutive trials for a duration of two minutes. The inter-trial interval was one minute, which is about the amount of time needed to change the odor stimulus. At the end of testing, subjects were hold in a temporary cage until all animals from the same home cage were tested. The test was videotaped and subsequently scored. Sniffing and direct interaction time (touching, biting, climbing the applicator) were quantified separately.

### Social tests

#### Three-chambered social approach test

Sociability and preference for social novelty and social recognition were tested in a three-chambered apparatus (Nadler et al., 2004). The subject mouse was first placed in the central, neutral chamber and allowed to explore for 10 minutes with all doors closed. Next, doors were opened and the mouse was allowed to freely explore the three empty chambers for an additional 10 minutes. Lack of side preference was confirmed during this habituation. The subject was then temporary placed in a holding cage while two empty wire cages which allow for olfactory, visual, auditory, and tactile contacts but not for sexual contact or fighting containing either an inanimate object (black cone) or a male mouse were placed in each of the testing chambers and the subject was returned to the apparatus for a 10-minute testing phase. Adult mice from the same strain that was previously habituated to the wire cup and did not exhibit aggressive behaviors but had no previous contact with the subject were used for unfamiliar mice. Unfamiliar mice were not used more than twice a day with at least two hours before two tests. At the end of testing, subjects were hold in a temporary cage until all animals from the same home cage have been tested. The side position of the interacting animal and the object was randomly determined. All the sessions were videotracked (Noldus Ethovision, Leesburg, VA, USA) and the amount of time spent in each chamber, close to the holding cages or in direct interaction with the holding cage was automatically calculated.

#### Male-female social interaction

Male-female social interactions were evaluated in in a regular clean cage during a 10-min test session as previously described (Scattoni et al., 2011). Each subject male was paired with an unfamiliar estrus C57BL/6J female under low light (10 lux) conditions. A total of twenty females were used for this test allowing to avoid to reuse the same female more than twice on the same day. The sessions were videotaped and ultrasonic vocalizations were recorded using an ultrasonic microphone with a 250 kHz sampling rate (Noldus Ultravox XT, Leesburg, VA) positioned 10 cm above the cage. The entire set-up was installed in a sound-attenuating room. Videos from the male subjects were subsequently manually scored to quantify (number of events and total time of male to female nose-to-nose sniffing, nose-to-anogenital sniffing and sniffing of other body regions. Ultrasonic vocalizations were played back and spectrograms were displayed using the Ultravox XT software and ultrasonic vocalizations were manually quantified.

#### Social transmission of food preference

The social transmission of food preference is a test of olfaction memory that involves a social component through the use of a demonstrator mouse (Wrenn et al., 2003). The demonstrator mouse is a conspecific mouse of same sex and similar age that was labeled by bleaching before testing. To minimize neophobia during the experiments, both subjects and demonstrator mice were habituated to eat powdered rodent chow (AIN-93M, Dyets, Inc., Bethlehem, PA) from 4-oz (113.40-g) glass food jar assemblies (Dyets, Inc., Bethlehem, PA). This habituation was performed for 48 hours in the mice home cage while the regular pellet chow was removed from the cages. After the habituation, both subject mice and demonstrator mice were food deprived for 18 to 24 hours before testing with free access to water. The test was divided into three phases.

Demonstrator exposition. During the first phase the demonstrator was presented with a jar of powder food mixed with either 1% cinnamon or 2% cocoa. The flavor was randomly assigned to the demonstrators so half of them received the cocoa flavored food while the other half received the cinnamon flavored food. Each demonstrator was used only once a day. The demonstrators were allowed to eat the flavored food for one hour. The jars were weighed before and after presentation to the demonstrators. The criterion for inclusion in the experiment was consumption of 0.2 g or more.

Interaction phase. After eating the flavored food, a demonstrator was placed in an interaction cage with the observer subject mouse and mice were allowed to freely interact for 30 minutes.

Choice phase. Immediately after the interaction phase, the observer mouse was placed in a clean cage and presented with one jar containing the flavor of food eaten by the demonstrator (cued) and another jar containing the other flavor and given one hour to freely explore the jar and eat. The demonstrator flavor and the position of the jar (front or back of the cage) was randomly assigned.

All phases were videotaped and food jars were weighed before and after the sessions to determine the amount of food eaten. At the end of testing, demonstrators and observers were hold in temporary cages until all animals from the same home cage had been tested. Video recordings from the interaction phase were used to score the number and total time of sniffing bouts from the observer to the nose or head of the demonstrator. Video recordings from the choice phase were used to score the total time spent in interaction with each food jar (mouse observed in the top of the jar with nose in jar hole).

### Avoidance, escape behavior and hyper-reactivity

Object avoidance and escape behavior was observed in several tests initially designed to assess other behaviors, including the novel object recognition, the marble burying and the nest building.

#### Novel object recognition

The novel object test for object recognition and memory takes place in an opacified open field arena (45 × 45 cm). The test involves a set of two unique novel objects, each about the size of a mouse, constructed from two different materials and non-uniform in shape. The test consisted of one 10-minute habituation session, a 5-minute familiarization session and a 5-minute recognition test, each videotracked (Noldus Ethovision). During the habituation, animals were allowed to freely explore an empty open field. At the end of the session, they were removed from the open field and place in a temporary clean holding cage for about two minutes. Two identical objects were placed on the median line at about 10 cm from each wall and the animal was returned to the open field and allowed to explore the objects for 5 minutes before being returned to its home cage. After one hour, one familiar object and one novel object were placed in the open field to the location where the identical objects were placed during the familiarization session and the mouse was allowed to explore them for a 5-minute recognition test. The side of the novel object position was randomly assigned so half of the animals were exposed to a novel object placed on the right of the open-field and half of the animals were exposed to a novel object placed on the left of the open-field.

Between each session, the open-field and the objects were carefully cleaned with 70% ethanol and let dry. Familiarization and recognition sessions were scored for total time spent investigating each object, the number of object interactions and the latency o the first object interaction. Time spend in each side during habituation and familiarization and time spent sniffing two identical objects during the familiarization phase were used to examine an innate side bias. Total time spent sniffing both objects was used as a measure of general exploration.

#### Marble burying test

The marble-burying assay is a tool for assessing either anxiety-like and/or repetitive-like behaviors in mice (Thomas et al., 2009). Subjects were tested in a regular clean cage (28 cm L × 18 cm W × 12 cm H) with 3 cm of fresh bedding. The subject was first placed in the empty cage for a 5-minute habituation. It was then temporary placed in an empty clean cage while 20 dark blue glass marbles (15 mm diameter) were positioned over the bedding equidistant in a 4×5 arrangement in order to cover the whole cage surface. The subject was then returned in the test cage and allowed to explore and bury the marbles during a 15-minute session that was videotaped. At the end of the session the subject was removed and the number of marbles buried (>50% marble covered by bedding material) was recorded.

#### Nest building

For small rodents, nests are important for heat conservation as well as for reproduction and shelter (Deacon, 2006). Mice were initially single housed in cages containing no environmental enrichment items such as bedding, cardboard houses or tunnels. To test their ability to build nests animals were temporarily single housed. One hour before the dark phase any building material present in the home cage was removed and replaced by two cotton nestlets (Ancare, NES3600 nestlets). The test was repeated twice and scored on the next morning of the second repeat using the following multi-criteria scale adapted from (Deacon, 2006) (maximum score= 11): nestlet shredding: 0 = not at all, 1 = partially, 2 = fully shredded; nestlet dispersion: 0 = nestlet dispersed all over the cage, 1 = mostly used to build nest, 2 = fully used to build a nest; nest density: 0: not dense, 1 = medium density, 2 = high density; nest shape: 0: no nest, 1 = ball shape, 2 = nest shape but no bottom, 3 = full nest; presence of walls: 0=no walls, 1 = partial walls, 2 = nest fully surrounded by walls; maximum score= 11.

#### Escape behavior

Escape behavior evaluated in three different tests all taking place in regular home cages (28 cm L × 18 cm W × 12 cm H) by counting the number of unsuccessful (mouse climbing on cage walls) or successful (mice jumping out of the cage) attempts. The three tests, selected for their increasing anxiogenic properties, were the habituation phase of the buried food test (first test in the home cage set-up, no object at the surface of the bedding), the repetitive novel object contact task (four objects visible at the surface of the bedding) and the marble burying test (twenty objects visible at the surface of the bedding). Each test was scored for ten minutes.

#### Hyper-reactivity

Hyper-reactivity was recorded by looking at touch escape response, positional passivity, trunk curl and catalepsy during the handling of the mice using the following scales. Touch escape to cotton-tipped applicator stroke from above starting light and slowly getting firmer recorded over five trials: 0 = no response, 1 = mild (escape response to firm stroke), 2 = moderate (rapid response to light stroke), 3 = vigorous (escape response to approach). Positional passivity or struggle response to sequential handling: 0 = struggles when restrained by tail, 1 = struggles when restrained by neck (finger grip, not scruffed), 2 = struggles when held supine (on back), 3 = struggles when restrained by hind legs, 4 = does not struggle. Trunk curl: 0 = absent, 1 = present. Catalepsy when subject front paws are positioned on a rod elevated 3 cm from floor, the amount of time the animal stayed immobile and kept its paws on rod was recorded, with a maximum cutoff of 120 seconds over three trials separated by 30 seconds. Hyper-reactivity was also observed in other tests such as the beam walking tests or the negative geotaxis test.

### Stereotypies, repetitive behavior, perseveration

#### Repetitive Novel Object Contact Task

This novel object investigation task looks for specific unfamiliar objects preference as well as patterned sequences of sequential investigations of those items (Pearson et al., 2011; Steinbach et al., 2016). Subjects were tested in a regular clean cage (28 cm L × 18 cm W × 12 cm H) with 1 cm of fresh bedding. The subject was first placed in the empty cage for a 20-minute habituation. It was then temporary placed in an empty clean cage while four unfamiliar objects (a Lego piece, 3 cm length; a jack, 4 cm length; a dice, 1.5 cm length; and a bowling pin,3.5 cm length) were place in the cage’s corners at approximately 3 cm from the edges. The subject was then able to investigate the environment and objects during a 10-minute session that was videotaped. The videos were manually scored for the occurrence of investigation of each of the four toys. Investigation was defined as clear facial or vibrissae contact with objects or burying of the objects. The number of contacts and the cumulative contact time was evaluated for each object. In order to determine if there was a genotype effect on the tendency to display preferences for particular toys, the frequencies of contact with each object were ranked in decreasing order from maximum to minimum preference for each subject and the frequencies were averaged by group, and compared. To assess the pattern of object investigation, each specific toy was given an arbitrary number (1–4) and all possible 3-digit and 4-digit combinations without repeat numbers were identified. For both three-and four-object sequences the total number of choice, the number of unique sequences, and the number of choices of the three most repeated sequence was calculated for each subject as described in (Steinbach et al., 2016). To take in account the overall mouse activity, the percentage of top, top two and top three preferred choices over the total number of choices were also calculated.

#### Barnes maze

The Barnes maze is a test of spatial memory comparable to a dry version of the Morris water maze (Barnes, 1979). In this assay, mice use spatial memory and navigation skills to orient themselves thanks to extra-maze cues placed in the test room, with the goal of locating one of twenty identical holes evenly spaced around the edge of a brightly-lit 100 cm diameter circular arena (Maze Engineers). While most of the holes (non-target) have nothing beneath them and lead nowhere, the target escape hole leads to shelter in a desirably darkened and enclosed goal box below the table. Two days before the beginning of the training, habituation was performed by allowing each subject to freely explore the arena (without escape box) under modest light for five minutes. At the end of the second habituation, subjects were pre-trained to learn of the presence of the escape hole by placing them for one minute in a clear box in the middle of the arena under bright light conditions. After one minute, the box was lifted up and the subject was gently guided near the escape hole selected randomly on the table, allowing it to enter the hole and remain inside for one minute. For the initial training, animals were trained for four days to locate the escape box (in a position different from the pre-training). All trials began with the subject in a clear box in the center of the table. The trial started when the box was lifted up. If the subject located and entered the escape box within three minutes, it was left in the box for one minute. If the subject failed to find the escape box within three minutes, it was gently guided to near the escape hole, and allowed to stay in the box for one minute. Animals received four trials per day with an inter-trial interval of twenty minutes for four days. After each trial, the maze and the escape box were cleaned using cleaning wipes to remove odors and fresh bedding was placed in the escape box. On the fifth day, animals were tested for three minutes without the escape box for a probe test. Time spent in the different quadrants was recorded. For the reversal training, the escape hole was moved to the opposite position on the maze and animals received four additional days of training followed by a reversal probe test on the fifth day. All trials were recorded by overhead camera (Noldus Ethovision) and scored for distance and latency to find escape box.

### Cognition

#### Y-maze test

Y-maze alternation is a test of working memory based on the natural tendency of mice to explore new territory whenever possible. Mice were placed in the center of a Y-maze (three 5 cm wide and 50 cm long arms, each set 130 degrees from each other) and given 15 min to freely explore the three arms of the maze. The number of arm entries and the number of triads were recorded in order to calculate the percentage of alternation. An entry occurs when all four limbs are within the arm. A successful score is defined by 3 successive choices that includes one instance of each arm by the total number of opportunities for alternation. A type 1 error is determined by three consecutive choices where the first and third choices are identical. A type 2 error is defined by three consecutive choices where the second and third choices are identical. Perseverance is defined as three or more repetitive entries in the same arm.

#### Contextual and cued fear conditioning

To isolate the effects of cued and contextual fear conditioning, a 3-day assay was employed. During the training session, the mice were placed in an ethanol cleaned contextual box with a bar floor, black and white striped walls in which all movements can be recorded (Med Associate fear conditioning boxes coupled with Noldus Ethovision for control an analysis) and given 5 minutes to habituate. Movements were then recorded for 540 seconds. At 120, 260 and 400 seconds after the beginning of the recording, mice were exposed to a 20-second tone (80dB, 2 KHz) and co-terminating shock (1 second, 0.7mA). Twenty-four hours after the training phase the animals were tested for contextual memory in the identical enclosure and movements were recorded for 240 seconds to assess the ability of the animal to remember the context in which the shocks had occurred the previous day. Forty-eight hours after the training phase animals were tested for cued memory in a different context (isopropanol cleaned, white wall insert over a mesh grid floor). They were recorded for 330 seconds and were presented with the identical tone from the training session at 120, and 260 seconds after the beginning of the recording session to assess the ability of each animal to remember the tone and pair it with the shock from training session. The three sessions were recorded using a camera located on the side of the boxes. Freezing, defined as lack of movement except for respiration, was scored using Noldus Ethovision software during each phase.

### Anxiety

#### Elevated zero-maze

Fear and anxiety were tested in an elevated zero-maze. The apparatus consisted of a circular black Plexiglas runway, 5 cm wide, 60 cm in diameter and raised 60 cm off the ground (Maze Engineers, Cambridge, MA, USA). The runway was divided equally into four alternating quadrants of open arcs, enclosed only by a 1 cm inch lip, and closed arcs, with 25 cm walls. All subjects received one 5-minute trial on two consecutive days starting in the center of a closed arm and were recorded by video-tracking (Noldus Ethovision, Leesburg, VA). Measures of cumulative open and closed arc times, latency to enter an open arc for the first time (for trials with a closed arc start), total open arm entries, latency to completely cross an open arc for the first time (for trials with a closed arc start) between two closed arcs, closed arc dipping (body in closed arc, head in open arc), open arc dipping (body in open arc, head outside of the maze) were calculated using the mean of the two trials.

#### Open field

The vertical activity in the open field was scored by counting the numbers of wall rears (while touching a side of the open field) and free-standing rears. The thigmotaxis was measured by quantifying the amount of time or distance travelled on the side of the open-field compared to the center of the open field.

### Statistical Analyses

*Shank3^Δ4-22^* wild-type, heterozygous and knock-out littermates were compared for each parameter. Statistical analyses were performed with SPSS 23.0 software using different types of ANOVA with or without repeated time measures with genotype as independent variable followed by Tukey pair-wise comparisons and correction for multiple comparisons if needed or equivalent non parametric tests when required. Newborn developmental milestones were analyzed by 2-way ANCOVA using genotype and gender as between-subject factors and litter number as co-variate to take in account possible gender and litter effects. As we did not observe a gender effect, males and females were grouped together in figures and tables. In order to account for possible cohort effects, cohorts 3 and 4 were analyzed either together using 2-way ANOVA with genotype and cohort as between-subject factors or separately using ANOVA or Kruskal-Wallis tests. Figures represent results for both cohorts analyzed together. Each cohort data and all statistical results including cohort effects are reported in Tables and corresponding Extended Tables. In tests comparing activity in two or more locations (open field thigmotaxis, social preference test, social transmission of food preference, novel object recognition, zero maze) genotype × zone interactions were assessed using repeated measures. When sphericity was found violated, the Greenhouse-Geisser values were reported. The distribution of the genotypes was compared to Mendelian expectation using Pearson’s chi-square test, the survival curves were analyzed using survival Kaplan-Meyer Chi-square. The comparison to chance level was evaluated using either one-sample T-test or Wilcoxon test. Normality was assessed using data visualization and Shapiro-Wilk test. All values are expressed as means ± s.e.m.

## Results

### Generation of a *Shank3^Δ4-22^* mouse with a complete deletion of the *Shank3* gene

A mouse line with a complete disruption of the *Shank3* gene was generated by retargeting ES cells previously used to disrupt exons 4 through 9. To do this, an additional loxP site was inserted directly after exon 22 while leaving intact the two existing loxP sites flanking exons 4 and 9 (Figure 1A). To generate the *Shank3^Δ4-22^* mouse line used in the present study, the floxed allele was then excised by breeding with a ubiquitous Cre transgenic line resulting in a deletion of exons 4 to 22 and therefore a constitutive disruption of all the *Shank3* murine isoforms.

Immunoblot analyses using antibodies which cross-react either with an epitope in the SH3 domain (antibody N367/62; Figure 1B left panel) or with the COOH terminal (Antibody H1160, Figure 1B right panel) showed no expression of Shank3 protein in post synaptic density fractions from *Shank3^Δ4-22^* homozygous mice and reduced expression consistent with haploinsuficiency in the heterozygotes. As in humans, in mice, the *Shank3* gene has 22 exons, spans ∼58 kb of genomic DNA, and undergoes complex transcriptional regulation controlled by a combination of five intragenic promoters and extensive alternative splicing resulting in in a complex pattern of mRNA and protein isoforms (Kouser et al., 2013; Speed et al., 2015; Waga et al., 2014; Wang et al., 2011; Wang et al., 2014). The loss of all known major *Shank3* mRNA isoforms was confirmed by RT-PCR (Figure 1C).

The *Shank3^Δ4-22^* mouse line was maintained on a C57BL/6 background by heterozygote × heterozygote mating, allowing for the production of all genotypes (wild-type, heterozygous and homozygous) as littermates. *Shank3^Δ4-22^* heterozygous and homozygous animals were viable, however abnormal Mendelian ratios were observed at the time of weaning, with a significant deficit for *Shank3^Δ4-22^* knockout mice (Figure 1D, Table 2). Adult survival curve between 1 and 22 months did not show a significant genotype difference with the current sample size, but there was evidence for higher numbers of deaths in *Shank3^Δ4-22^* homozygous mice between 18 and 22 months (Figure 1E, Table 2). Although the human clinical *SHANK3* mutation is hemizygous, for completeness, we have conducted our studies in *Shank3*-null mutant mice (homozygous knockout, KO), along with their heterozygous (Het) and wild-type (WT) littermates. The KO mice are instrumental to understand the function of Shank3, while the Het mice have significantly greater construct validity for PMS, a haploinsufficiency syndrome. To ensure the robustness of behavioral abnormalities in the adult mice, two cohorts representing all three genotypes were compared. All the cohorts used in the present study are described in Table 3.

**Table 2:** Genotype distribution at weaning and postnatal mortality.

| genotype distribution at weaning |  |  |  |  |  |  |  |  |
| --- | --- | --- | --- | --- | --- | --- | --- | --- |
|  | WT | Het | KO | %WT | %Het | %KO | Chi-square (df2) | Asymp p-value |
| All animals, observed N | 365 | 686 | 278 | 27.46 | 51.62 | 20.92 | 12.78 | 0.0017 |
| All animals, expected N | 332.25 | 664.5 | 332.25 | 25.00 | 50.00 | 25.00 |  |  |
| All animals, residual N | 32.75 | 21.5 | -54.25 | 2.46 | 1.62 | -4.08 |  |  |
| Males, observed N | 185 | 357 | 147 | 26.85 | 51.81 | 21.34 | 5.10 | 0.0781 |
| Males, expected N | 172.25 | 344.5 | 172.25 | 25.00 | 50.00 | 25.00 |  |  |
| Males, residual N | 12.75 | 12.5 | -25.25 | 1.85 | 1.81 | -3.66 |  |  |
| Females, observed N | 180 | 329 | 131 | 28.13 | 51.41 | 20.47 | 8.01 | 0.0182 |
| Females, expected N | 160 | 320 | 160 | 25.00 | 50.00 | 25.00 |  |  |
| Females, residual N | 20 | 9 | -29 | 3.13 | 1.41 | -4.53 |  |  |

**Table 3:** Cohorts used and order of behavioral testing. For adult animals, the age indicated corresponds to the average age of the cohort. For each cohort all mice were born within two weeks of each other. *: missing animals due to technical problems during startle recording.

| Cohort 1 (10 litters) - developmental milestones |  |  |  |  |
| --- | --- | --- | --- | --- |
|  | WT | Het | KO | Age at testing |
| All animals | 14 | 30 | 10 | P0-P21 |
| Males | 7 | 16 | 5 | P0-P21 |
| Females | 7 | 14 | 5 | P0-P21 |

| Cohort 2 (10 litters) - ultrasonic vocalizations |  |  |  |  |
| --- | --- | --- | --- | --- |
|  | WT | Het | KO | Age at testing |
| All animals | 16 | 32 | 9 | P6 |
| Males | 4 | 15 | 6 | P6 |
| Females | 12 | 17 | 3 | P6 |

| Cohorts 3 and 4 - adult behavior |  |  |  |  |  |  |  |  |
| --- | --- | --- | --- | --- | --- | --- | --- | --- |
|  | Cohort 2 |  |  |  | Cohort 3 |  |  |  |
|  | WT | Het | KO | Age at testing | WT | Het | KO | Age at testing |
| Handling, cage observation, neurological and motor reflexes | 11 | 10 | 9 | P86-P90 | 8 | 9 | 10 | P103-P107 |
| 15-month weight | 8 | 8 | 6 | P460 | 5 | 7 | 4 | P455 |
| 20-month weight | 7 | 7 | 2 | P610 | 4 | 5 | 3 | P600 |
| open field | 11 | 10 | 9 | 93-94 | 8 | 9 | 10 | P106-P108 |
| zero maze | 11 | 10 | 9 | P95-P96 | 8 | 9 | 10 | P109-P110 |
| Y-maze | 11 | 10 | 9 | P99-P101 | 8 | 9 | 10 | P114-P122 |
| beam walking | 11 | 10 | 9 | P102-P103 | 8 | 9 | 10 | P124-P125 |
| grip strength | 11 | 10 | 9 | P104 | 8 | 9 | 10 | P125 |
| gait analysis | 11 | 10 | 9 | P105 | 8 | 9 | 10 | P126 |
| rotarod | 11 | 10 | 9 | P107-P108 | 8 | 9 | 10 | P127 |
| 3-chambered social interaction task | 11 | 10 | 9 | P113-P114 | 8 | 9 | 10 | P130-P131 |
| nest building | 11 | 10 | 9 | P120 | 8 | 9 | 10 | P137 |
| novel object | 11 | 10 | 9 | P123-125 | 8 | 9 | 10 | P139-140 |
| Fear conditioning | 11 | 10 | 9 | P126-P128 | 8 | 9 | 10 | P141-P143 |
| Startle response * | 11 | 10 | 9 | P137-P139 | 3* | 4* | 4* | P155-P157 |
| Prepulse inhibition | 11 | 10 | 9 | P137-P139 | 8 | 9 | 10 | P155-P157 |
| Tail flick | 11 | 10 | 9 | P144-P145 | 8 | 9 | 10 | P158-P159 |
| Olfactory habituation/dishabituation | 11 | 10 | 9 | P149-P157 | 8 | 9 | 10 | P162-P165 |
| buried food | 11 | 10 | 9 | P163-P164 | 8 | 9 | 10 | P178 |
| social transmission of food preference | 11 | 10 | 9 | P206-P215 | 8 | 9 | 10 | P185-P192 |
| Marble burying | 11 | 10 | 8 | P227-P228 | 8 | 9 | 10 | P197 |
| 4-object repetitivenovelobject contact task | 11 | 10 | 8 | P232 | 7 | 9 | 9 | P215 |
| Male-female social interaction | 11 | 10 | 8 | P240-241 | 7 | 9 | 9 | P217-219 |
| Barnes maze | 11 | 10 | 7 | P247-P274 | 7 | 9 | 8 | P222-P250 |

### Developmental milestones in *Shank3^Δ4-22^* neonates

Ten litters were used to study developmental milestones. The average litter size was 7.2 pups (ranging from 5 to 9), with 54 surviving passed postnatal day 2 (28 males and 26 females). As very limited gender effects were observed (see Table 4 for detailed analysis), males and females were analyzed together using both genotype and gender as fixed factors and the litter number as a covariate.

**Table 4:**
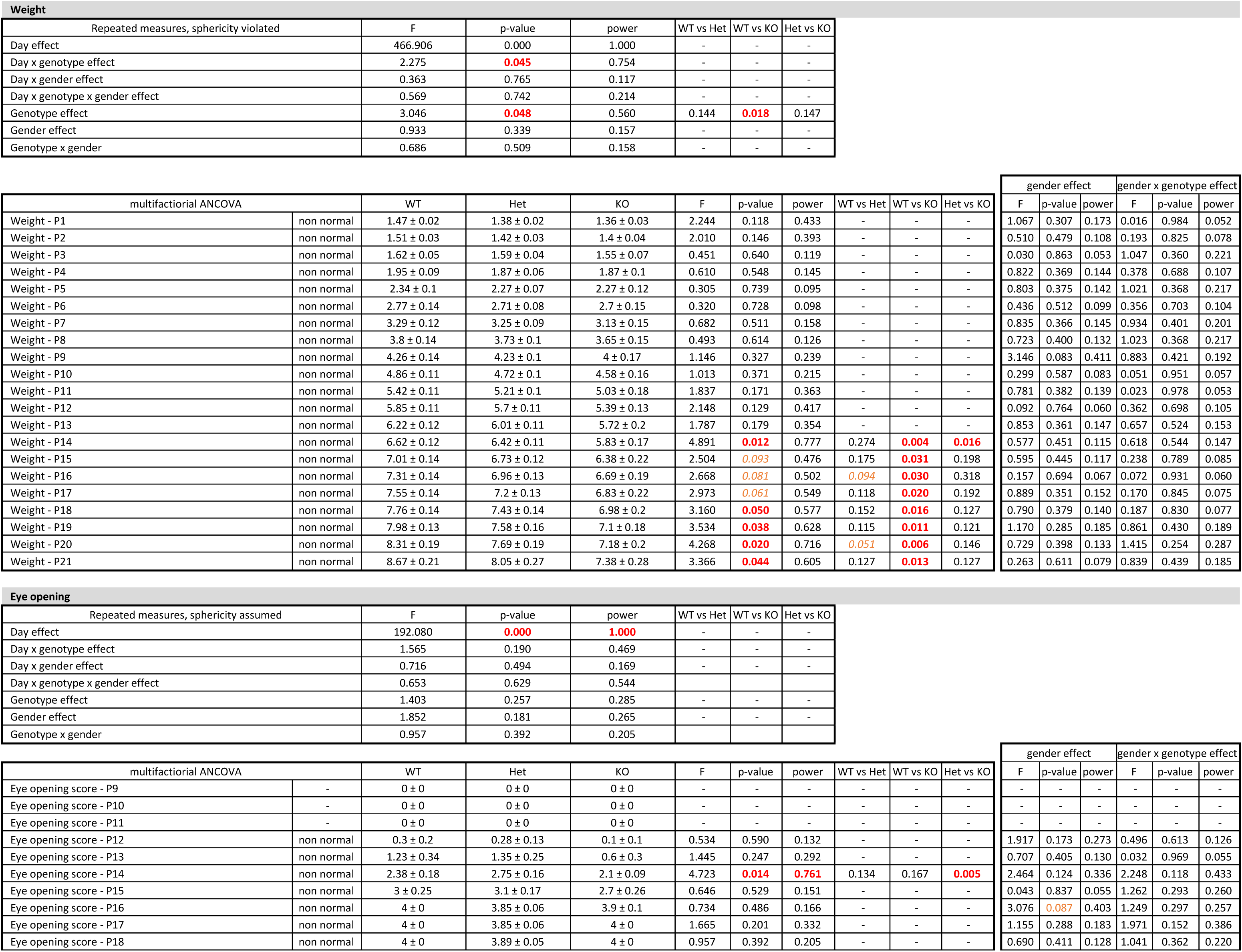
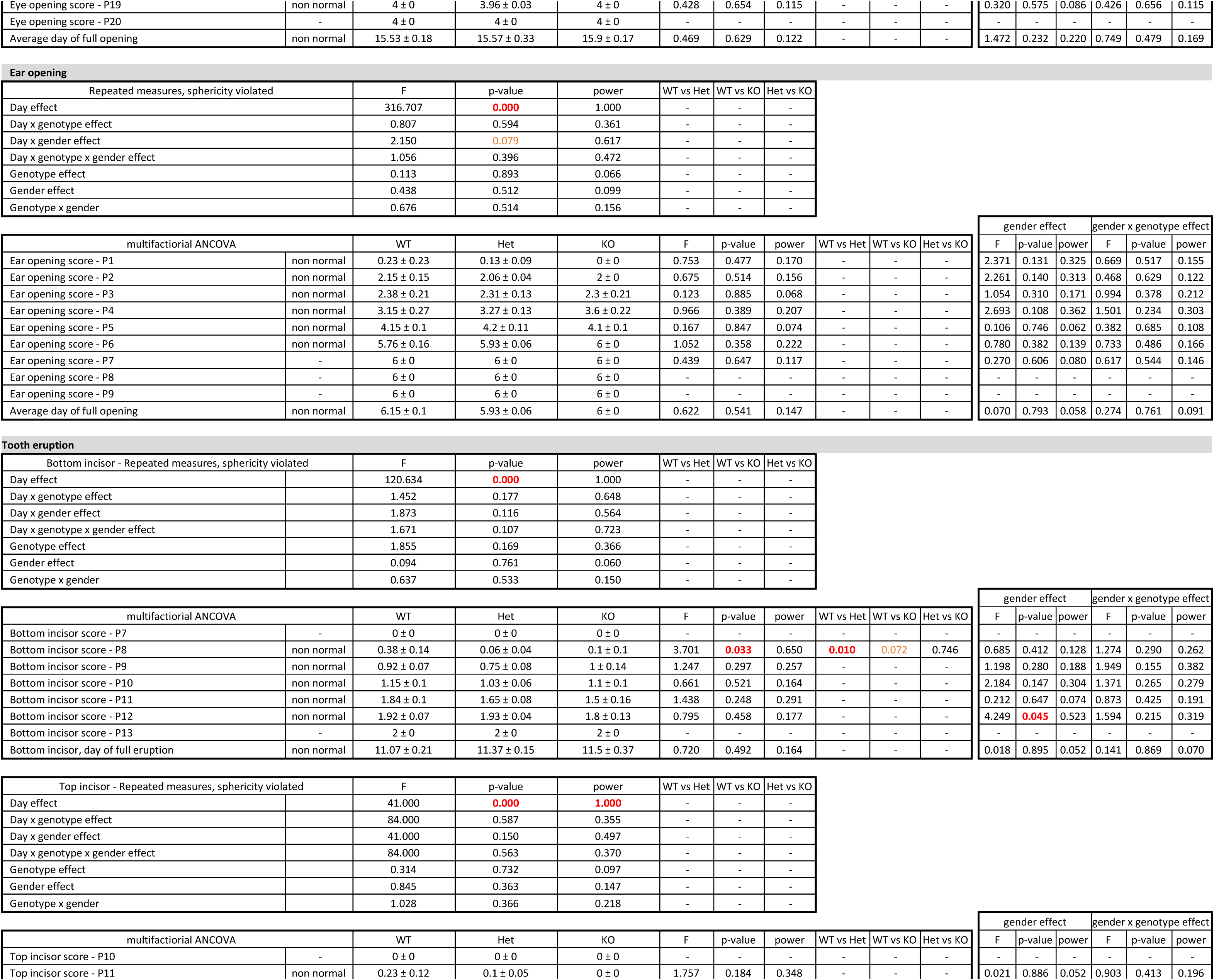
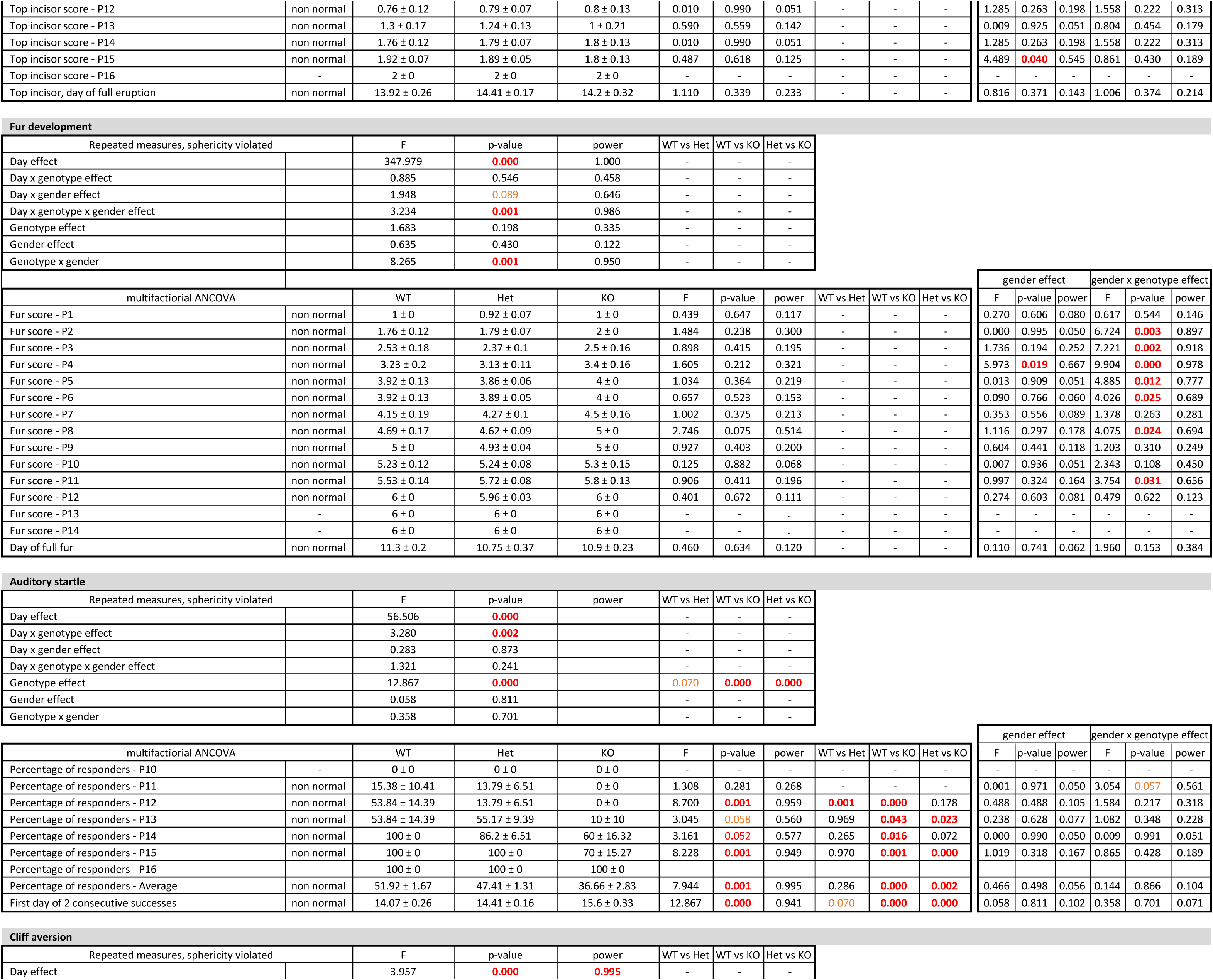
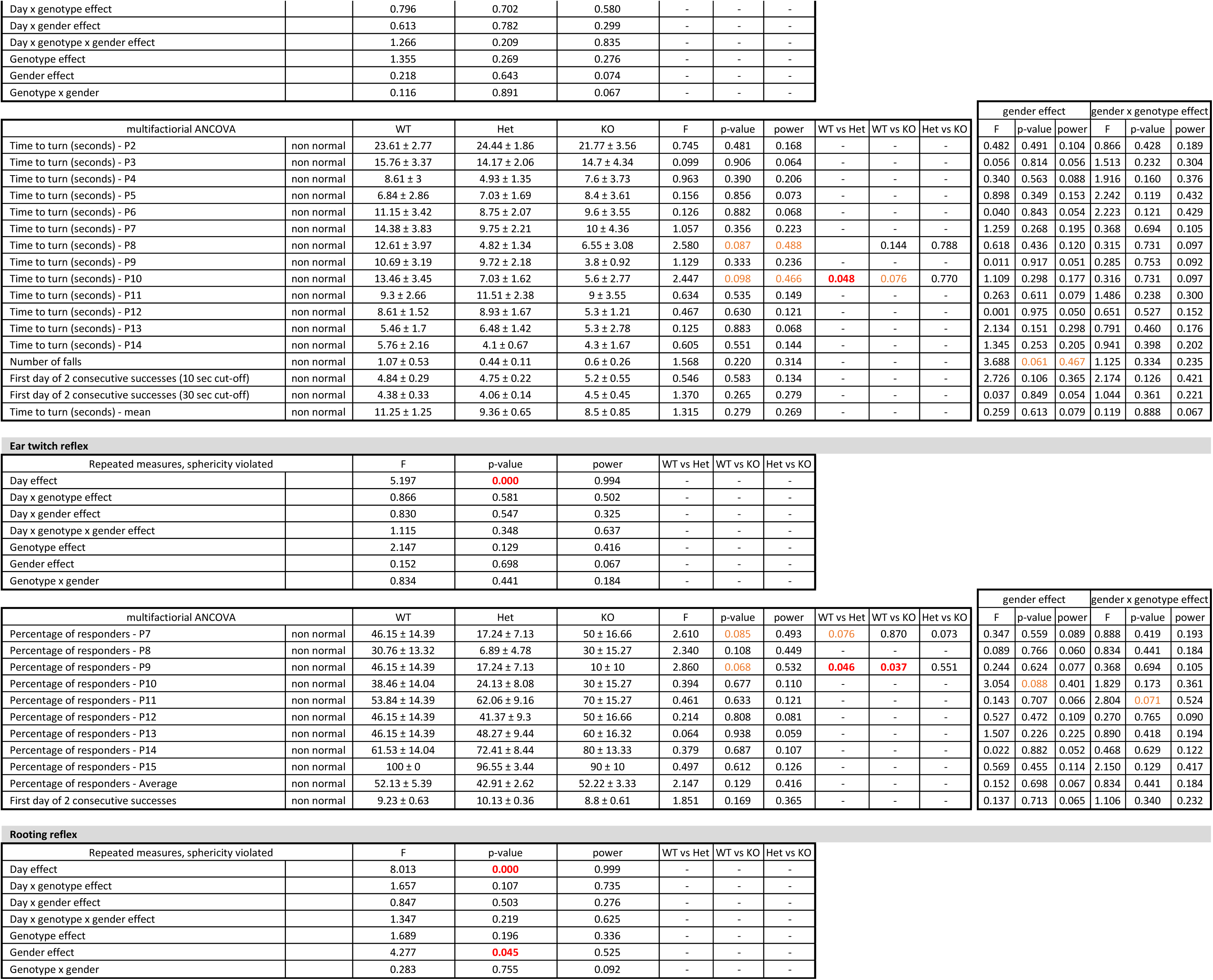
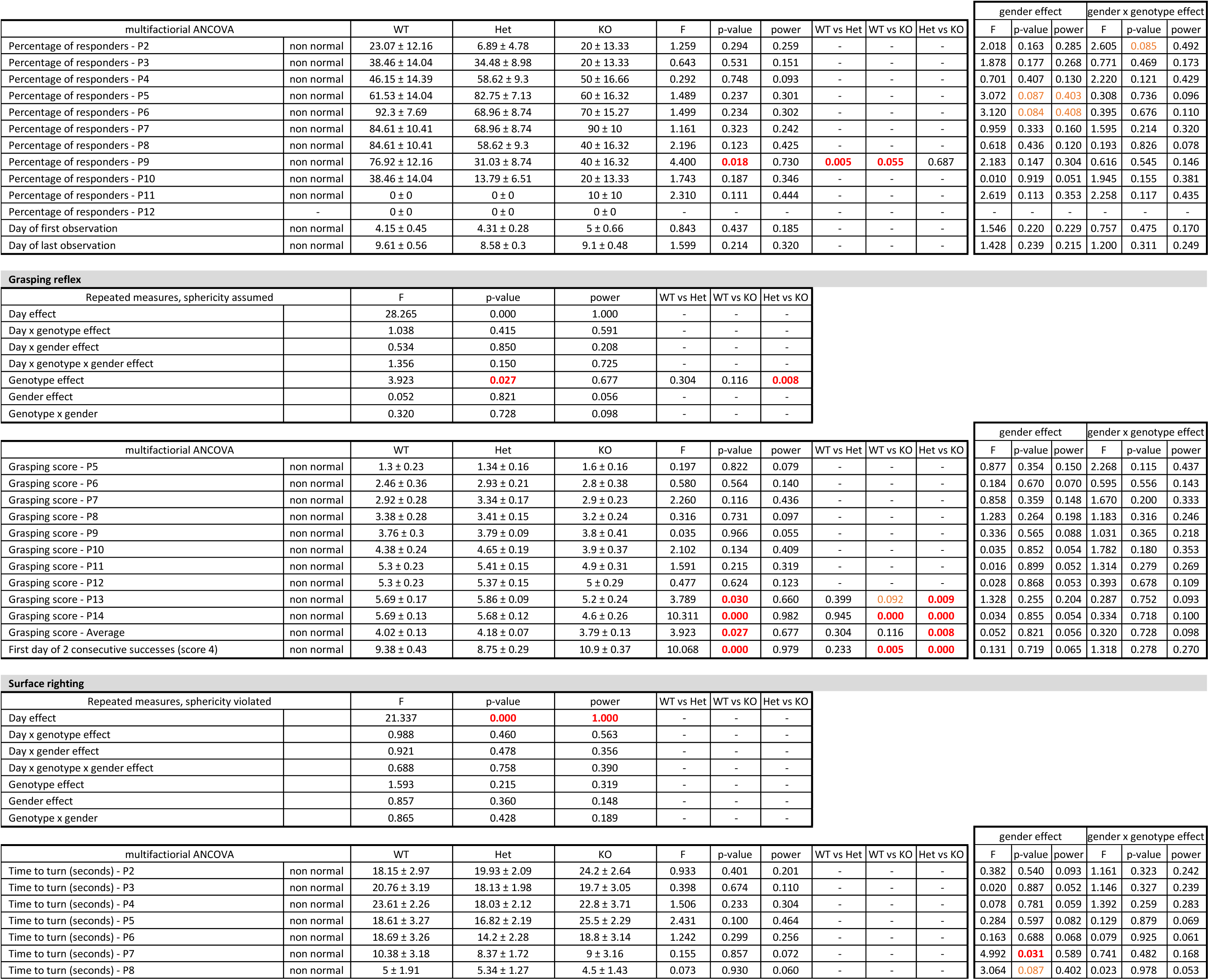
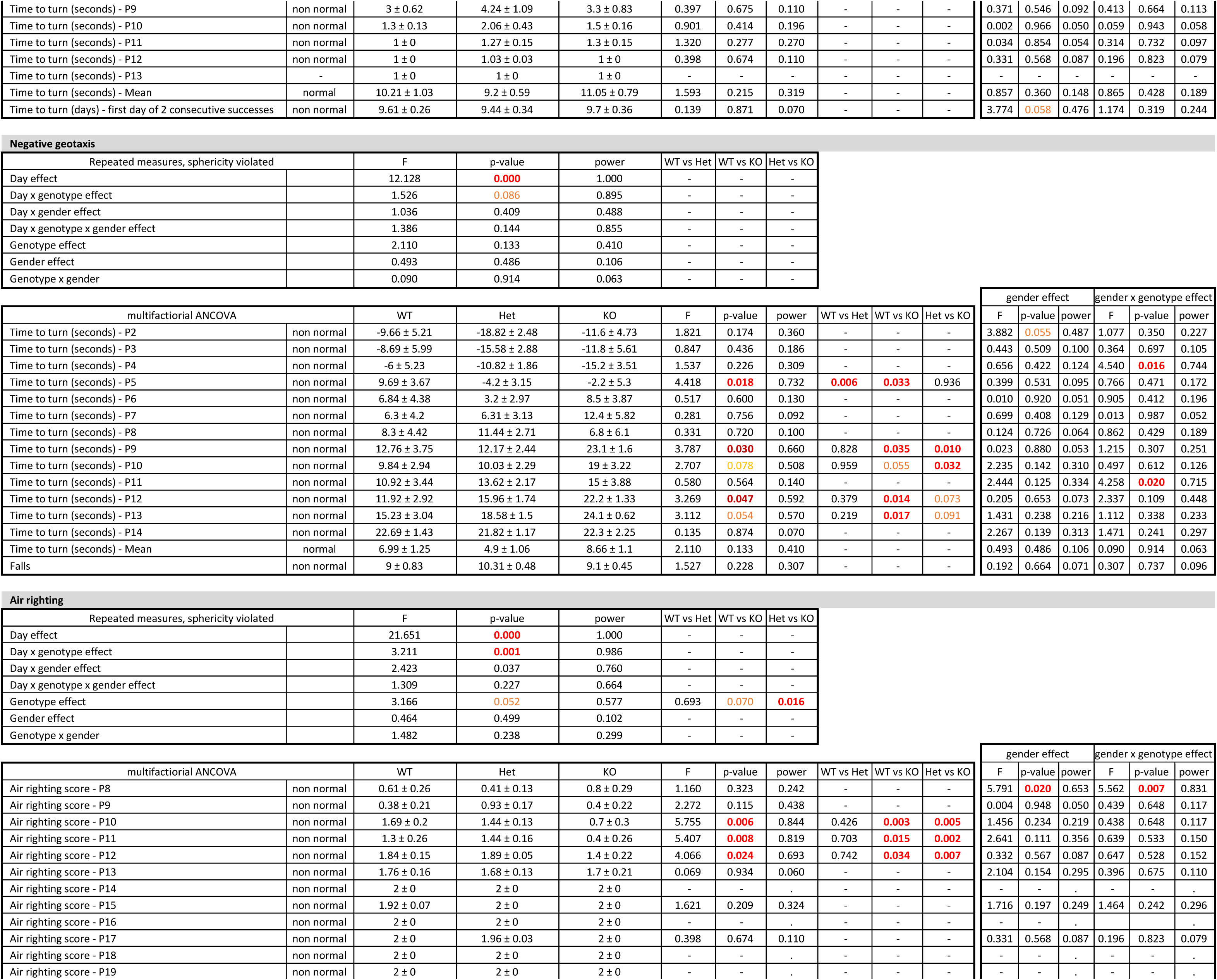
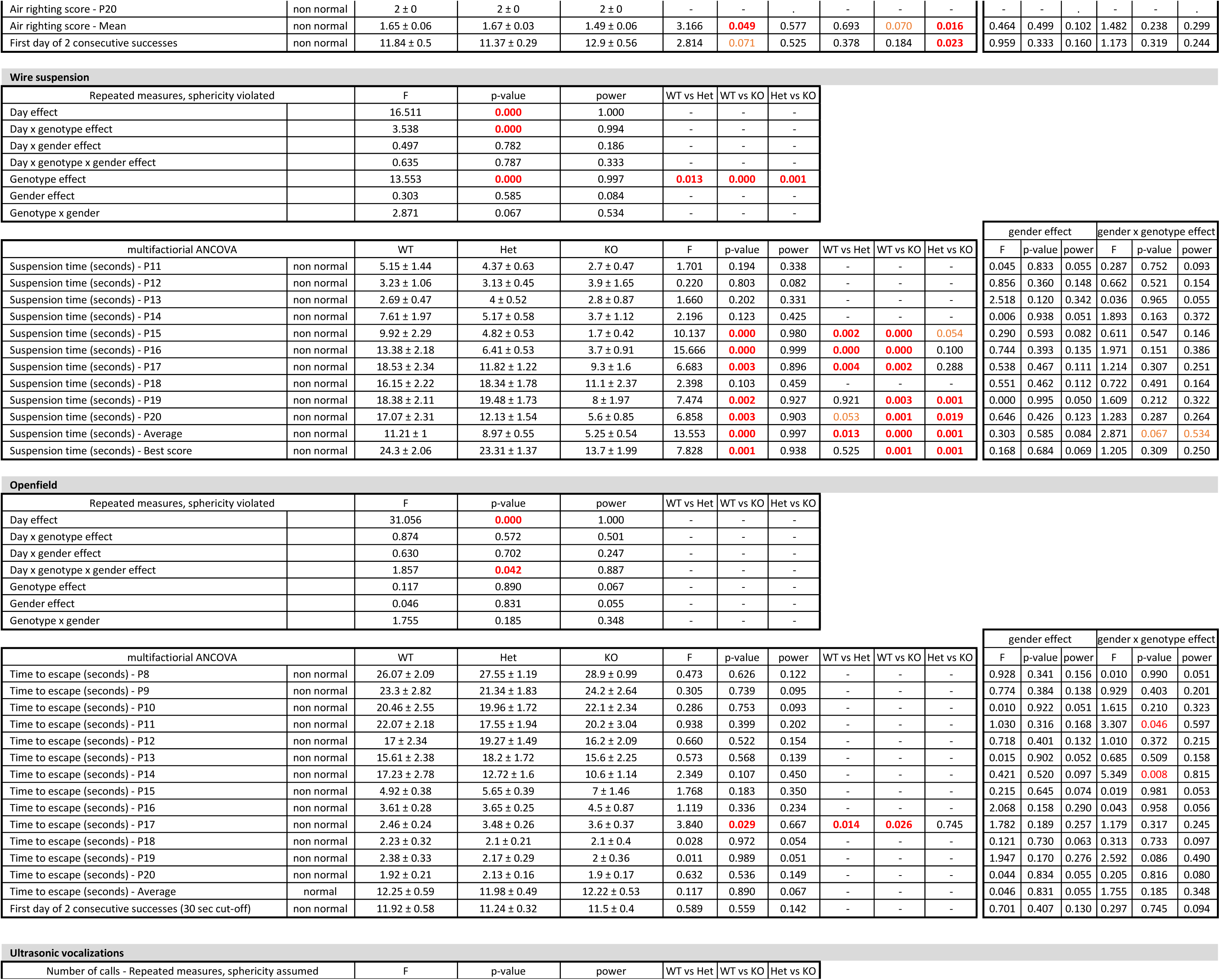
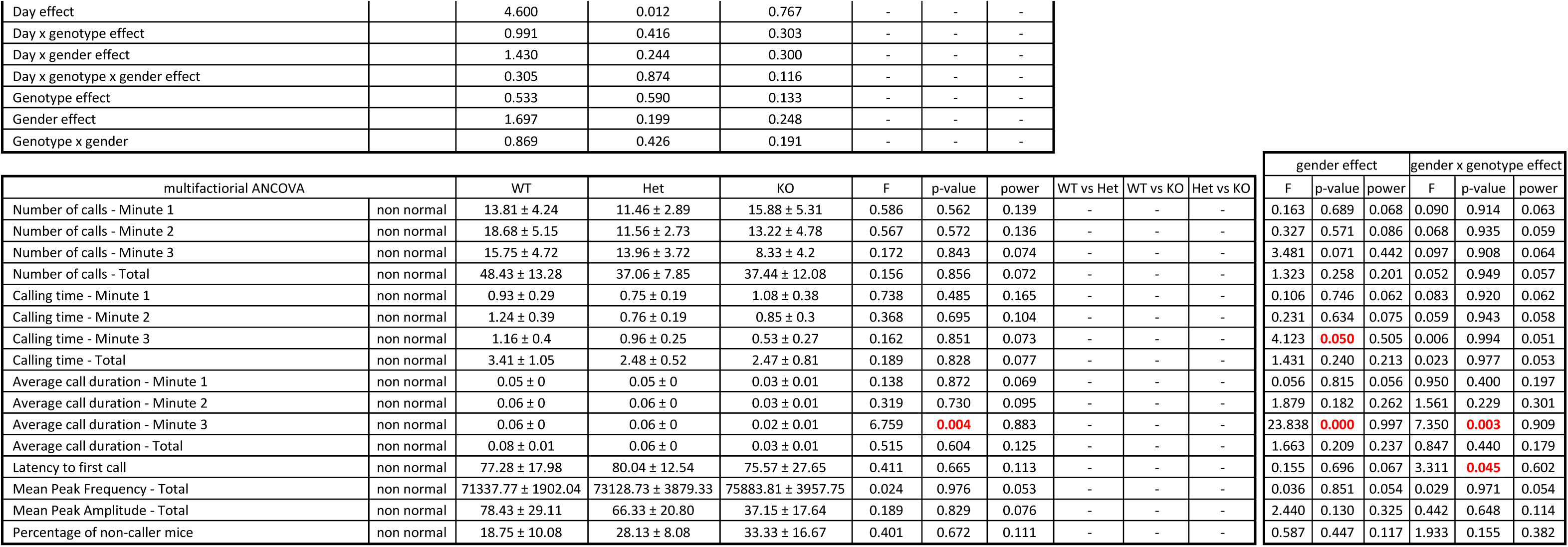
Detailed results and statistical analyses related to developmental milestones. WT, wild-type mice; Het, heterozygous mice; KO, homozygous knockout mice. Group values are reported as means ± s.e.m. Red font indicates significant results (p<0.05), orange font indicates trends (0.1<p<0.05).

Developmental delays were observed in the *Shank3^Δ4-22^* homozygote neonates in several of the parameters studied (Figure 2, Extended Figure 2-1 and Table 4). While the birth weight was not significantly different, the growth rate of *Shank3^Δ4-22^* homozygote pups was slower and by P14, the weight of *Shank3^Δ4-22^* homozygous mice was significantly lower than the weight of their wild-type littermates (Figure 2A). No differences were observed in any of the other physical developmental milestones, including eye opening, ear opening, tooth eruption or fur development (Extended Figure 2-1 A-D and Table 4).

**Figure 2:**
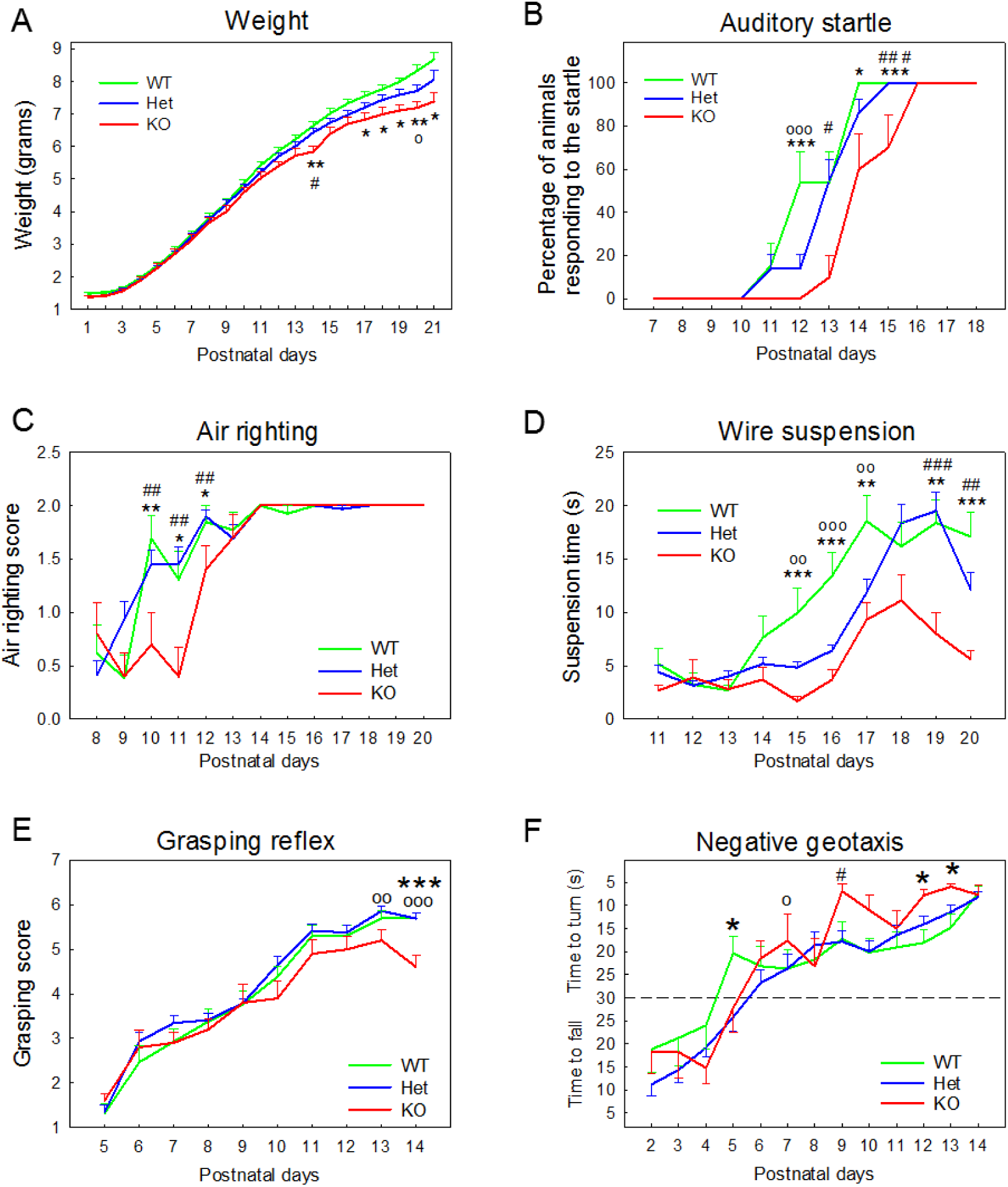
Delayed developmental milestones of in *Shank3^Δ4-22^*-deficient mice. Analysis of markers of developmental milestones revealed genotype differences in *Shank3^Δ4-22^* wild-type, heterozygous and homozygous pups between postnatal days 1 and 21 on measures of **(A)** body weight, **(B)** auditory startle, **(C)** air righting, **(D)** wire suspension, **(E)** grasping reflex and **(F)** negative geotaxis. Additional milestones (jar opening, tooth eruption, fur development, eye opening, rooting reflex, cliff aversion, ear twitch, surface righting, open field crossing and ultrasonic vocalizations are displayed in Extended Figure 2-1) WT, wild-type mice; Het, heterozygous mice; KO, homozygous knockout mice. *: WT vs KO; o: WT vs Het, #: Het vs KO. *: p<0.05, **: p<0.1, ***: p<0.001.

A significant delay was observed for *Shank3*^Δ4-22^ homozygotes in the response to auditory startle (Figure 2B) and in the mid-air righting task (Figure 2C) although all the mice were able to properly respond at the end of the observation period. In the wire suspension (Figure 2D) and grasping reflex (Figure 2E) tasks, however, not only was the acquisition of the response delayed, but *Shank3^Δ4-22^* homozygous animals remained significantly impaired until the time of weaning. In the negative geotaxis test, an initial delay was observed at P5 were most wild-type animals were able to turn while homozygous and heterozygous *Shank3^Δ4-22^* animals were still falling or staying in the starting position (Figure 2F). Moreover, after P9 when most of the animals were able to master the task, higher reactivity (characterized by a shorter latency to turn) was observed for the *Shank3^Δ4-22^* homozygous mice. The acquisition of the rooting reflex was similar for the three groups however a premature disappearance of the reflex was observed in both the *Shank3^Δ4-22^* heterozygous and homozygous pups (Extended Figure 2-1 E and Table 4).

Other sensory-motor and neurological milestones such as cliff aversion, ear twitch, surface righting, negative geotaxis and open-field crossing (Extended Figure 2-1 F-I and Table 4) were not significantly affected by the disruption of the *Shank3* gene.

Ultrasonic vocalizations were recorded at postnatal day on an independent cohort of mice and a genotype difference was detected in the number and quality of ultrasonic vocalizations emitted by the pups (Table 4). *Shank3^Δ4-22^* heterozygous and homozygous mice emitted fewer ultrasonic vocalizations than wild-type littermates (Extended Figure 2-1 K and Table 4). The total calling time was also affected with *Shank3^Δ4-22^*-deficient mice both spending less time calling and having shorter calls than wild-type littermates. Additionally, the peak amplitude was shorter in *Shank3^Δ4-22^*-deficient mice. However, none of these parameters were significantly different probably due to a high interindividual variability within each group with some animals emitting no vocalizations during the three-minute recording. The percentage of non-callers was higher, although not significantly, in *Shank3^Δ4-22^*-deficient animals. Genotype did not affect the latency to the first call nor the peak frequency of calls and no difference was observed in the time course of the emission of ultrasonic vocalizations.

### Adult general health in *Shank3^Δ4-22^*-deficient mice

Adult *Shank3^Δ4-22^* mice were evaluated for general health at three months of age (Table 5). The three genotypes did not differ on physical measure of weight and length. Additional weight measures at the age of fifteen and twenty months showed a trend in reduced weight of *Shank3^Δ4-22^* homozygous mice compared to their littermates. Genotypes scored similarly and in the normal range for other physical characteristics including coat appearance (grooming, piloerection, patches of missing fur on face or body), skin pigmentation, whisker appearance, wounding and palpebral closure. Observation in a beaker or after transfer to a housing cage revealed no abnormalities in term of spontaneous general activity, stereotypies (rears, jumps, circling, wild running), transfer arousal, gait, pelvic and tail elevation.

**Table 5:** Detailed results and statistical analyses related to general health, physical factors, gross appearance and spontaneous activity. WT, wild-type mice; Het, heterozygous mice; KO, homozygous knockout mice. Group values are reported as means ± s.e.m. Red font indicates significant results (p<0.05), orange font indicates trends (0.1<p<0.05). Individual results and statistical analyses for cohorts 1 and 2 are available in Extended Table 5-1

| Physical factors and gross appearance |  |  |  |  |  |  |  |  |  |  |  |  |  |  |  |  |  |
| --- | --- | --- | --- | --- | --- | --- | --- | --- | --- | --- | --- | --- | --- | --- | --- | --- | --- |
|  | Cohorts 1 and 2 |  |  |  |  |  |  |  |  |  |  |  |  |  |  |  |  |
|  | test | data structure | WT | Het | KO | genotype |  |  | cohort |  |  | genotype x cohort |  |  | pairwise comparisons |  |  |
|  |  |  |  |  |  | F | p-value | power | F | p-value | power | F | p-value | power | WT vs Het | WT vs KO | Het vs KO |
| Weight at 3 months (grams) | 2-way ANOVA | normal | 26.33 ± 0.87 | 27.18 ± 0.57 | 26.01 ± 0.54 | 1.241 | 0.298 | 0.258 | 2.546 | 0.117 | 0.347 | 2.000 | 0.146 | 0.394 | - | - | - |
| Weight at 15 months (grams) | 2-way ANOVA | normal | 33.41 ± 2 | 31.06 ± 1.21 | 29.36 ± 1.73 | 1.578 | 0.222 | 0.310 | 0.269 | 0.608 | 0.079 | 0.491 | 0.617 | 0.123 | - | - | - |
| Weight at 20 months (grams) | 2-way ANOVA | normal | 32.9 ± 1.86 | 31.43 ± 1.2 | 28.84 ± 1.5 | 0.982 | 0.390 | 0.199 | 0.034 | 0.856 | 0.054 | 0.018 | 0.982 | 0.052 | - | - | - |
| Length | 2-way ANOVA | non normal | 16.45 ± 0.24 | 16.84 ± 0.24 | 16.73 ± 0.25 | 1.062 | 0.353 | 0.226 | 91.207 | 0.000 | 1.000 | 0.471 | 0.627 | 0.123 | - | - | - |
| Coat appearance | 2-way ANOVA | non normal | 2.63 ± 0.13 | 2.89 ± 0.07 | 2.94 ± 0.05 | 2.558 | 0.087 | 0.489 | 2.615 | 0.112 | 0.355 | 1.424 | 0.250 | 0.291 | 0.119 | 0.050 | 0.915 |
| Skin color | - | - | 0 ± 0 | 0 ± 0 | 0 ± 0 | - | - | - | - | - | - | - | - | - | - | - | - |
| Whisker barbering | - | - | 0 ± 0 | 0 ± 0 | 0 ± 0 | - | - | - | - | - | - | - | - | - | - | - | - |
| Patches of missing fur on face | - | - | 0 ± 0 | 0 ± 0 | 0 ± 0 | - | - | - | - | - | - | - | - | - | - | - | - |
| Patches of missing fur on body | - | - | 0 ± 0 | 0 ± 0 | 0 ± 0 | - | - | - | - | - | - | - | - | - | - | - | - |
| Wounding | 2-way ANOVA | non normal | 0.15 ± 0.11 | 0 ± 0 | 0.05 ± 0.05 | 1.078 | 0.348 | 0.229 | 0.028 | 0.869 | 0.053 | 0.279 | 0.758 | 0.092 | - | - | - |
| Body tone | 2-way ANOVA | non normal | 1.1 ± 0.07 | 1.1 ± 0.07 | 1 ± 0.07 | 0.563 | 0.573 | 0.138 | 2.225 | 0.142 | 0.310 | 0.563 | 0.573 | 0.138 | - | - | - |
| Palpebral closure | - | - | 0 ± 0 | 0 ± 0 | 0 ± 0 | - | - | - | - | - | - | - | - | - | - | - | - |
| Spontaneous piloerection | - | - | 0 ± 0 | 0 ± 0 | 0 ± 0 | - | - | - | - | - | - | - | - | - | - | - | - |

| Jar observation |  |  |  |  |  |  |  |  |  |  |  |  |  |  |  |  |  |
| --- | --- | --- | --- | --- | --- | --- | --- | --- | --- | --- | --- | --- | --- | --- | --- | --- | --- |
|  | Cohorts 1 and 2 |  |  |  |  |  |  |  |  |  |  |  |  |  |  |  |  |
|  | test | data structure | WT | Het | KO | genotype |  |  | cohort |  |  | genotype x cohort |  |  | pairwise comparisons |  |  |
|  |  |  |  |  |  | F | p-value | power | F | p-value | power | F | p-value | power | WT vs Het | WT vs KO | Het vs KO |
| Body position | 2-way ANOVA | non normal | 4.15 ± 0.08 | 4.1 ± 0.07 | 4.26 ± 0.1 | 0.702 | 0.500 | 0.162 | 4.949 | 0.031 | 0.588 | 0.139 | 0.871 | 0.070 | - | - | - |
| Spontaneous activity | 2-way ANOVA | non normal | 1.68 ± 0.1 | 1.57 ± 0.11 | 1.63 ± 0.11 | 0.223 | 0.801 | 0.083 | 0.282 | 0.598 | 0.082 | 0.279 | 0.758 | 0.092 | - | - | - |
| Latency to sit/stand (seconds) | - | - | 0 ± 0 | 0 ± 0 | 0 ± 0 | - | - | - | - | - | - | - | - | - | - | - | - |
| Latency to rear (seconds) | 2-way ANOVA | non normal | 8.94 ± 1.36 | 8.05 ± 1.03 | 5.63 ± 1.02 | 2.137 | 0.128 | 0.418 | 0.036 | 0.850 | 0.054 | 0.046 | 0.955 | 0.057 | - | - | - |
| Repeated jumps (percentage of mice) | 2-way ANOVA | non normal | 15.78 ± 8.59 | 10.52 ± 7.23 | 26.31 ± 10.37 | 0.702 | 0.500 | 0.162 | 4.949 | 0.031 | 0.588 | 0.139 | 0.871 | 0.070 | - | - | - |
| Circling (percentage of mice) | 2-way ANOVA | non normal | 5.26 ± 5.26 | 10.52 ± 7.23 | 10.52 ± 7.23 | 0.289 | 0.750 | 0.093 | 5.177 | 0.027 | 0.607 | 0.289 | 0.750 | 0.093 | - | - | - |
| Urination | 2-way ANOVA | non normal | 0.47 ± 0.19 | 0.1 ± 0.1 | 0.15 ± 0.11 | 1.540 | 0.224 | 0.312 | 0.023 | 0.881 | 0.052 | 1.213 | 0.306 | 0.253 | - | - | - |
| Defecation (number) | 2-way ANOVA | non normal | 1.57 ± 0.35 | 0.94 ± 0.29 | 0.94 ± 0.27 | 1.065 | 0.352 | 0.226 | 0.003 | 0.958 | 0.050 | 2.592 | 0.085 | 0.494 | - | - | - |
| respiration | - | - | 2 ± 0 | 2 ± 0 | 2 ± 0 | - | - | - | - | - | - | - | - | - | - | - | - |
| tremor | - | - | 0 ± 0 | 0 ± 0 | 0 ± 0 | - | - | - | - | - | - | - | - | - | - | - | - |

| Cage transfer |  |  |  |  |  |  |  |  |  |  |  |  |  |  |  |  |  |
| --- | --- | --- | --- | --- | --- | --- | --- | --- | --- | --- | --- | --- | --- | --- | --- | --- | --- |
|  | Cohorts 1 and 2 |  |  |  |  |  |  |  |  |  |  |  |  |  |  |  |  |
|  | test | data structure | WT | Het | KO | genotype |  |  | cohort |  |  | genotype x cohort |  |  | pairwise comparisons |  |  |
|  |  |  |  |  |  | F | p-value | power | F | p-value | power | F | p-value | power | WT vs Het | WT vs KO | Het vs KO |
| Transfer arousal | 2-way ANOVA | non normal | 3.21 ± 0.22 | 3.21 ± 0.22 | 3.15 ± 0.2 | 0.037 | 0.964 | 0.055 | 1.470 | 0.231 | 0.221 | 2.055 | 0.139 | 0.404 | - | - | - |
| Gait | 2-way ANOVA | non normal | 0 ± 0 | 0.15 ± 0.08 | 0.05 ± 0.05 | 1.783 | 0.178 | 0.356 | 0.003 | 0.957 | 0.050 | 0.637 | 0.533 | 0.151 | - | - | - |
| Pelvic elevation | 2-way ANOVA | non normal | 2 ± 0 | 2.15 ± 0.08 | 2 ± 0.07 | 1.730 | 0.188 | 0.347 | 0.141 | 0.709 | 0.066 | 0.141 | 0.869 | 0.071 | - | - | - |
| Tail elevation | 2-way ANOVA | non normal | 1.89 ± 0.15 | 1.73 ± 0.18 | 1.21 ± 0.18 | 4.003 | 0.024 | 0.691 | 4.469 | 0.039 | 0.545 | 0.204 | 0.816 | 0.080 | 0.530 | 0.009 | 0.043 |

### Motor functions in *Shank3^Δ4-22^*-deficient mice

Motor functions were examined using several different paradigms (Table 6). Footprint gait analysis showed normal stance and sway but increased stride in *Shank3^Δ4-22^* homozygous mice compared to wild-type and heterozygous animals (Figure 3A) and reduced spontaneous locomotion was observed during a one-hour open field session in both *Shank3^Δ4-22^* heterozygous and homozygous mice (Figure 3B). Across the 60-minute session, the time course for total distance traversed by all three genotypes declined as expected, representing habituation to the open-field. However, while the distance traveled during the first ten minutes was similar for the three groups, the decline was faster for *Shank3^Δ4-22^* homozygous mice, possibly reflecting a higher fatigability. Similarly, in the accelerating rotarod test, which assay for gait, balance, motor coordination and endurance, shorter latencies to fall where observed in *Shank3^Δ4-22^-*deficient mice after the first trial, with a milder phenotype observed in the heterozygotes compared to homozygotes. When examining learning in this paradigm, characterized by an improvement of performance (latency to fall) over the trials, *Shank3^Δ4-22^* heterozygous and homozygous animals failed to improve over time, in contrast to wild-type animals which showed typical learning (Figure 3C).

**Table 6:**
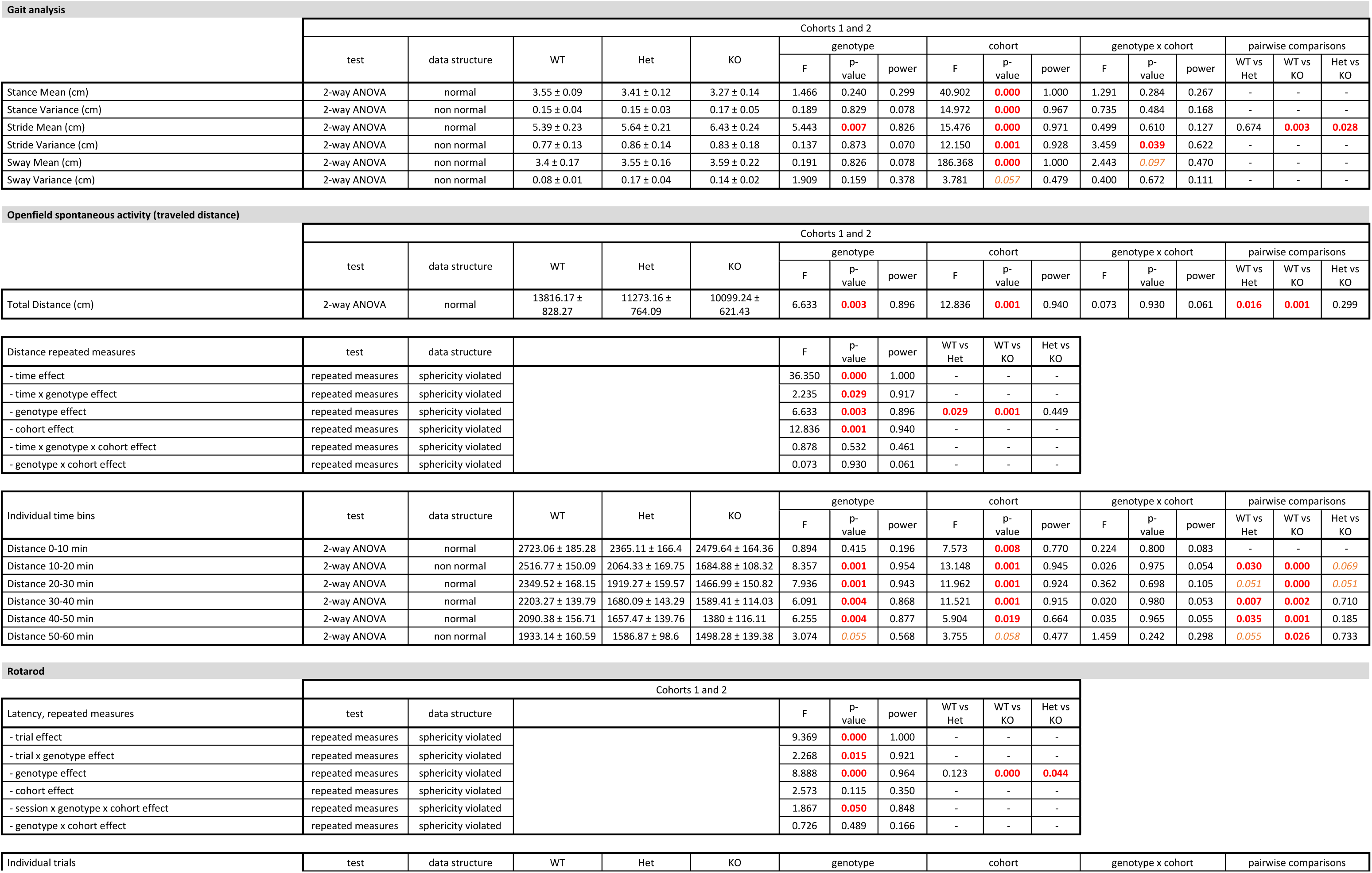
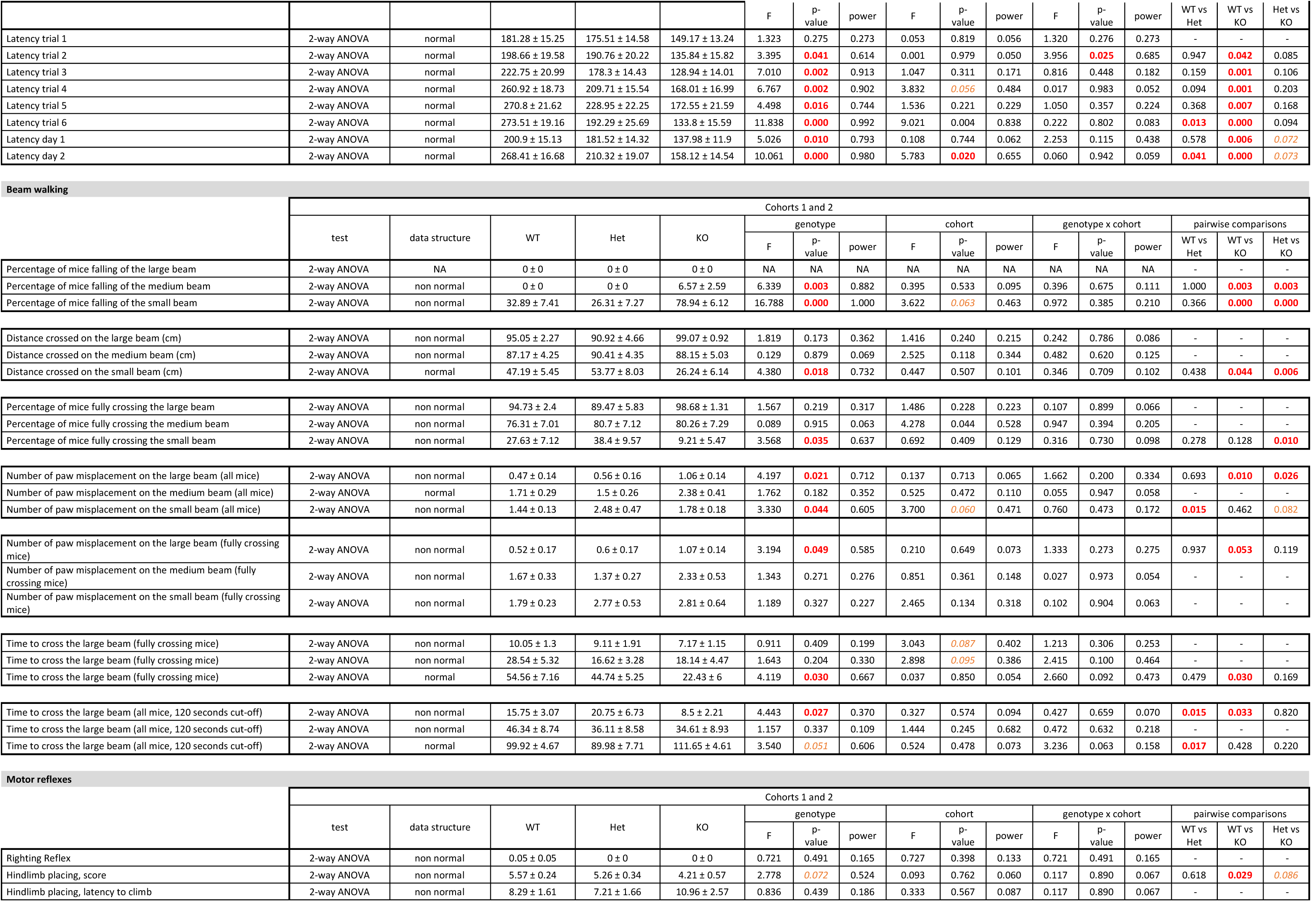
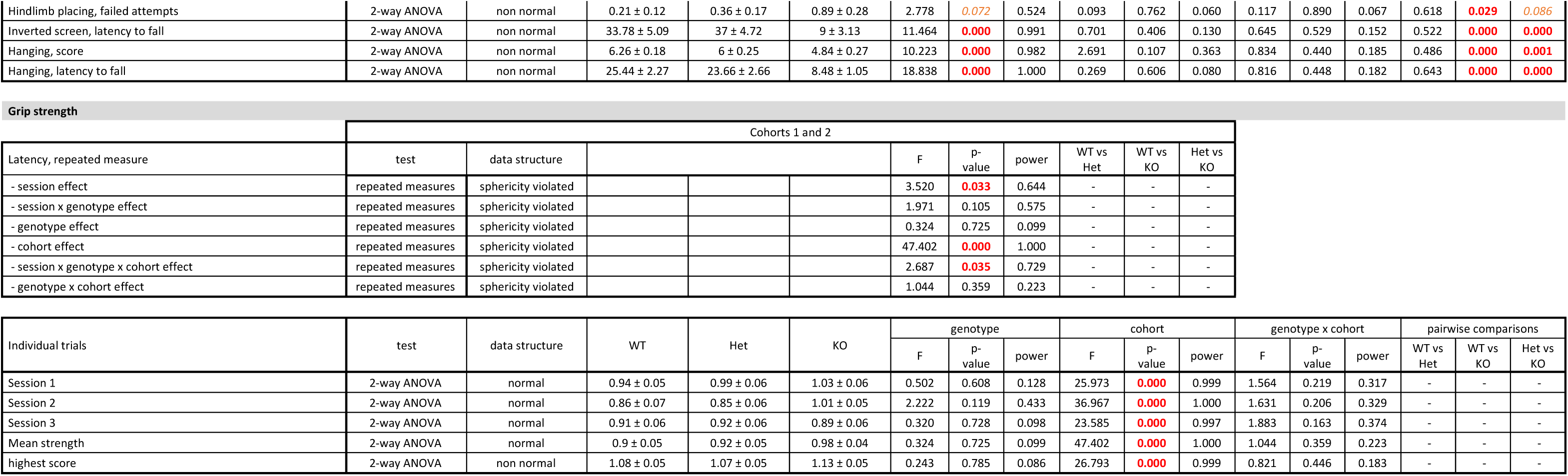
Detailed results and statistical analyses related to motor functions. WT, wild-type mice; Het, heterozygous mice; KO, homozygous knockout mice. Group values are reported as means ± s.e.m. Red font indicates significant results (p<0.05), orange font indicates trends (0.1<p<0.05). Individual results and statistical analyses for cohorts 1 and 2 are available in Extended Table 6-1

**Figure 3:**
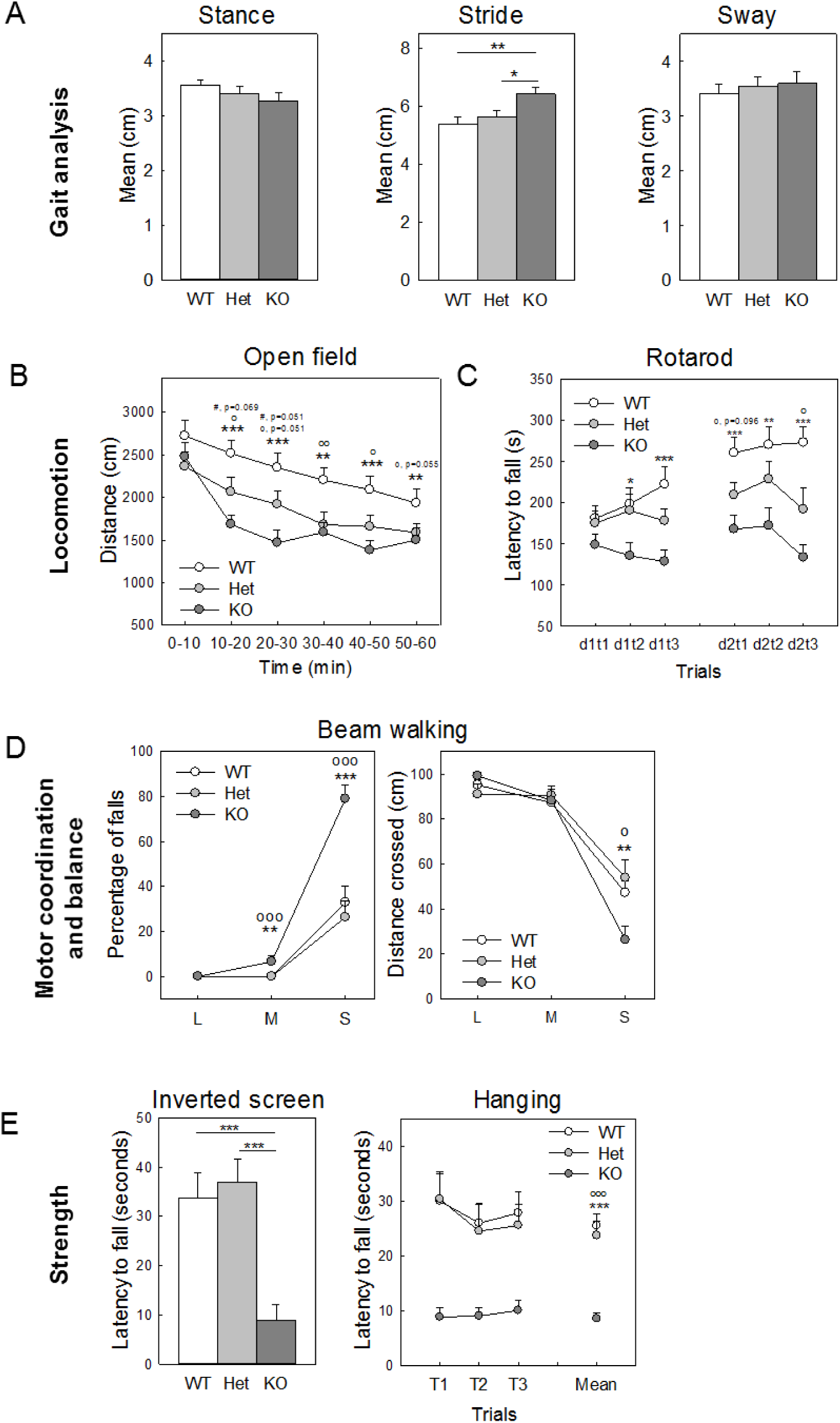
Impaired motor performances in in *Shank3^Δ4-22^*-deficient mice. **(A)** Average stance, stride and sway. Gait analysis showed an increase stride length in *Shank3^Δ4-22^* homozygous mice. **(B)** Distance travelled during a 60-minute session in an open field. Spontaneous locomotor activity in the open field was reduced in *Shank3^Δ4-22^* homozygous mice relative to other genotypes. **(C)** Latency to fall over 6 trials (3 trials per day for 2 consecutive days) in the accelerating rotarod task. Motor learning on the accelerating rotarod was deficient in *Shank3^Δ4-22^* homozygous mice compared to wild-type animals as they failed to improve over time. Heterozygous mice had an intermediate phenotype. **(D)** Percentage of falls and distance crossed during the beam walking test. While not different on the large (L, 1 inch) and medium (M, 1/2 inch) beams, *Shank3^Δ4-22^* homozygous mice were strongly impaired in the small (S, 1/4 inch) beam walking test as shown by a significant increase of the number of falls and a decrease of the distance crossed. **(E)** Strength and endurance measured in the inverted screen and hanging tests. Endurance strength was significantly impaired in *Shank3^Δ4-22^* homozygous mice as they exhibited significantly shorter latency to fall in both the inverted screen and hanging tests. Additional results of motor tests (Hind limb placing and grip strength are available in Extended Figure 3-1. WT, wild-type mice; Het, heterozygous mice; KO, homozygous knockout mice. *: WT vs KO; o: WT vs Het, #: Het vs KO. *: p<0.05, **: p<0.1, ***: p<0.001.

Impairment of motor coordination and balance was also observed in *Shank3^Δ4-22^* homozygous in the beam walking test (Figure 3D, Table 6) as well by reduced strength and endurance in both the inverted screen and hanging tests (Figure 3E), but with no differences in forelimb grip strength (Extended Figure 3-1 A). There was also a trend toward an increased number of failed attempt in the hind limb placing for *Shank3^Δ4-22^* homozygous mice, compared to their littermates (Extended Figure 3-1 B).

### Sensory abilities in *Shank3^Δ4-22^*-deficient mice

For all sensory-related assays, detailed results are reported in Table 7.

**Table 7:**
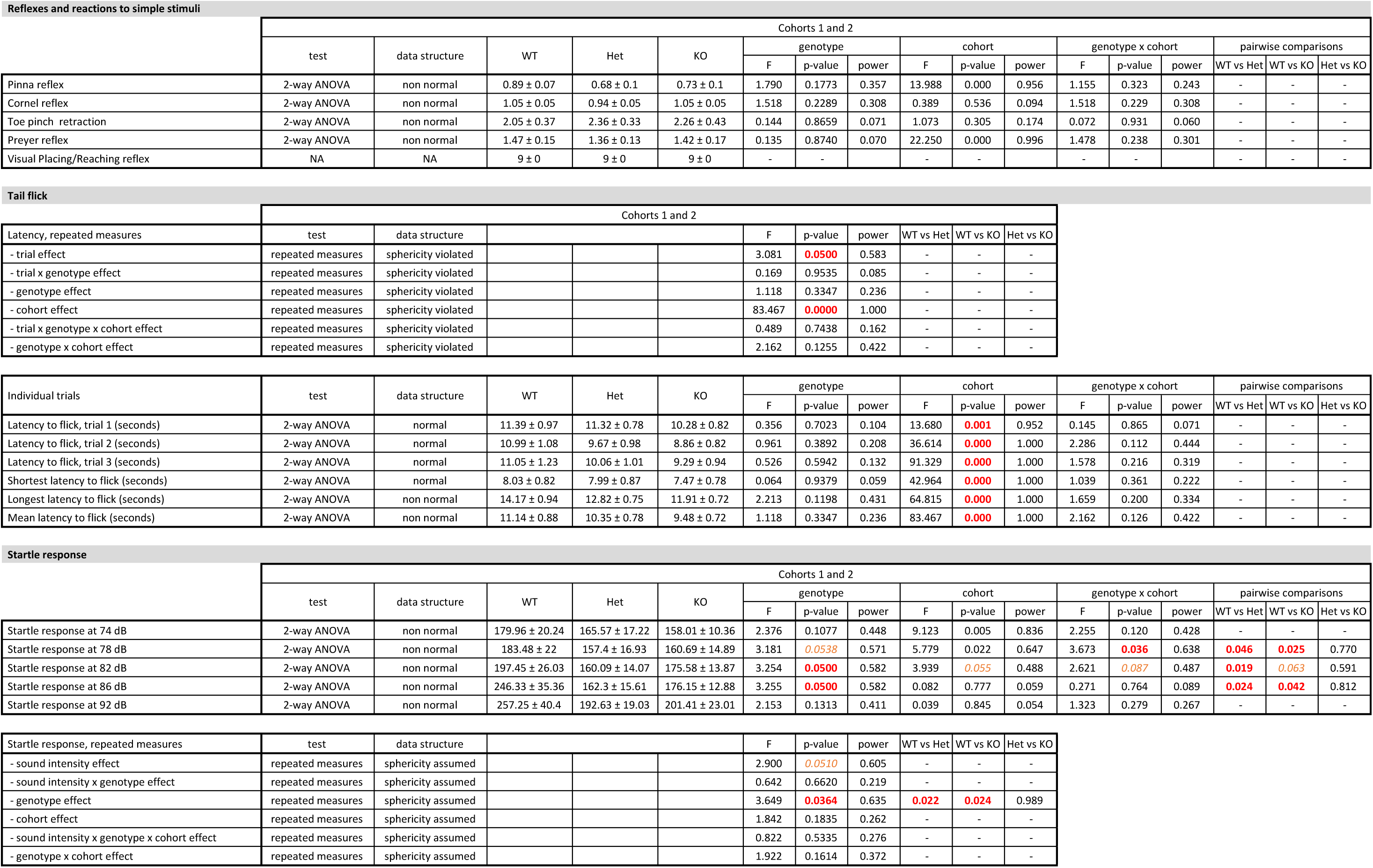
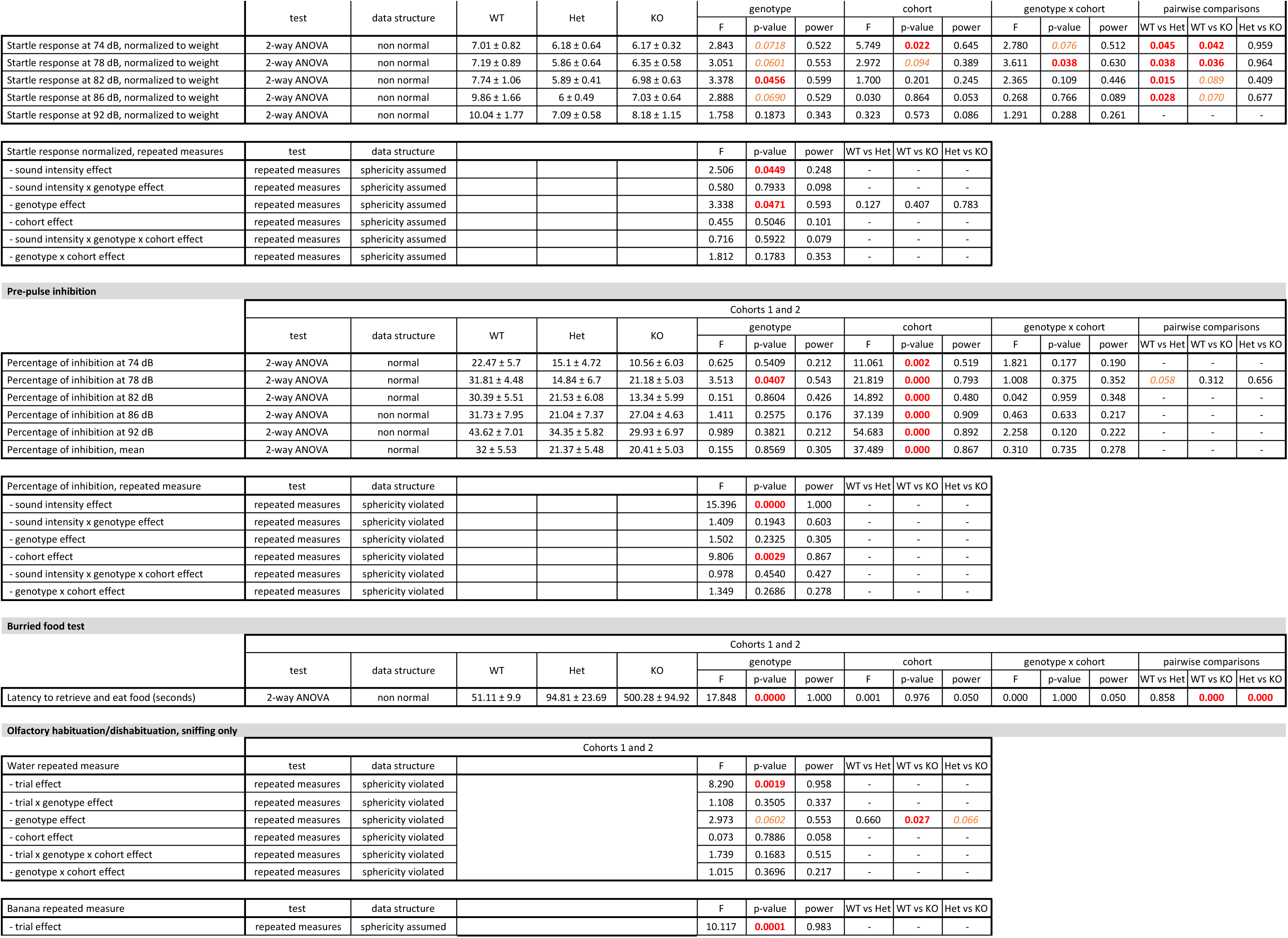
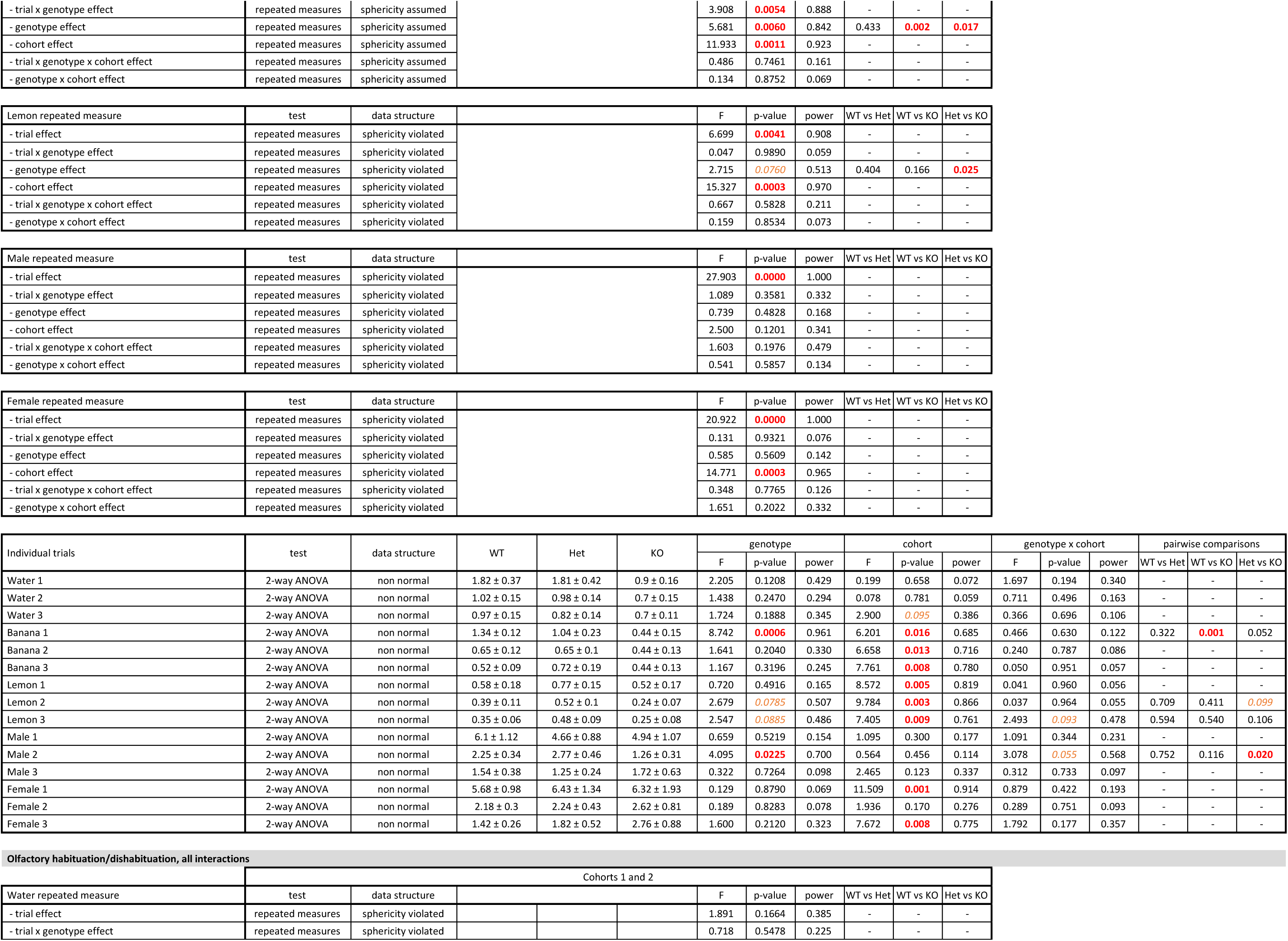
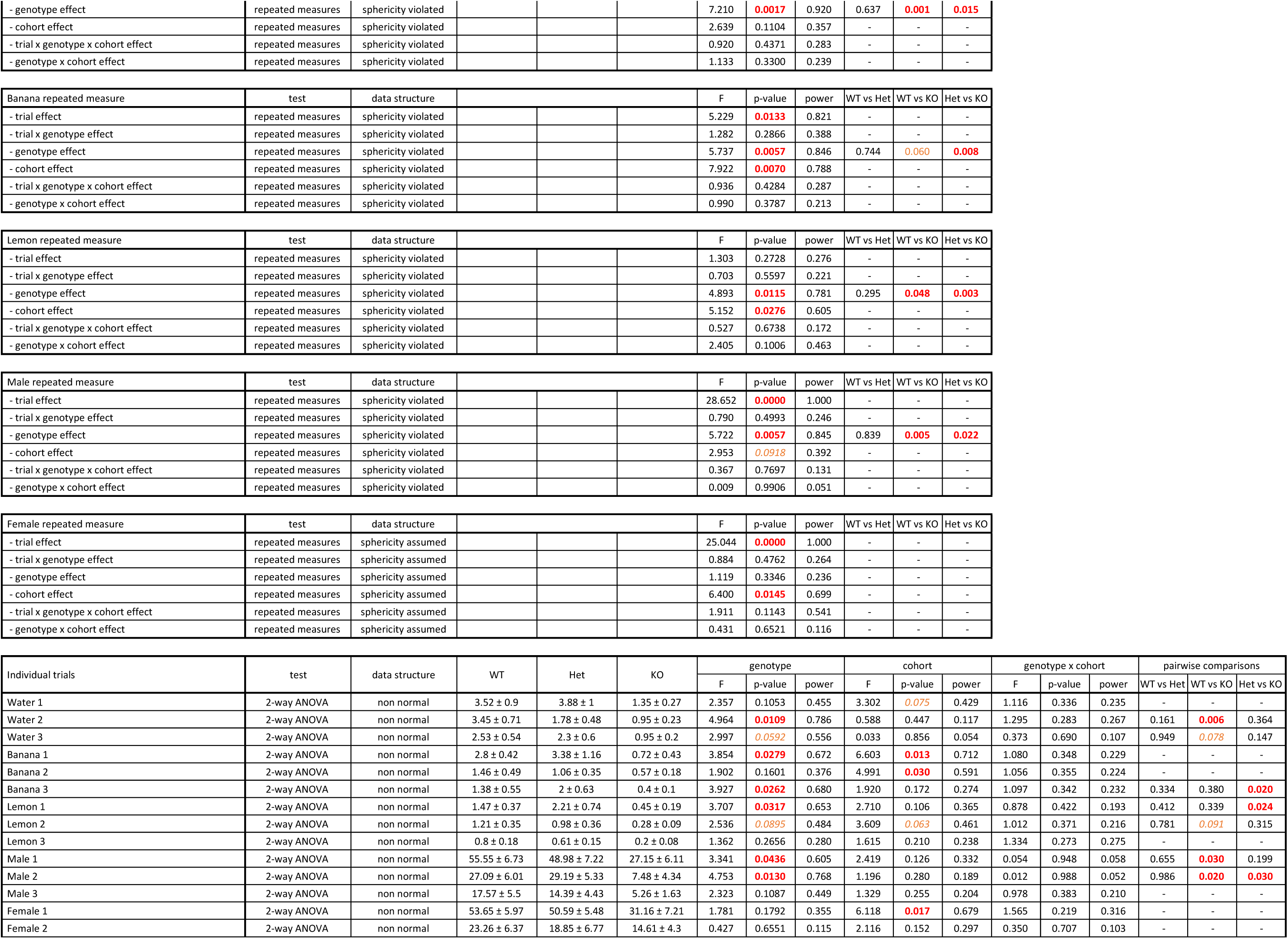
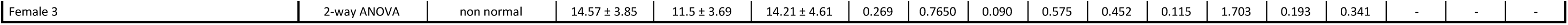
Detailed results and statistical analyses related to the sensory profile. WT, wild-type mice; Het, heterozygous mice; KO, homozygous knockout mice. Group values are reported as means ± s.e.m. Red font indicates significant results (p<0.05), orange font indicates trends (0.1<p<0.05). Individual results and statistical analyses for cohorts 1 and 2 are available in Extended Table 7-1

| Reflexes and reactions to simple stimuli |  |  |  |  |  |  |  |  |  |  |  |  |  |  |  |  |  |
| --- | --- | --- | --- | --- | --- | --- | --- | --- | --- | --- | --- | --- | --- | --- | --- | --- | --- |
|  | Cohorts 1 and 2 |  |  |  |  |  |  |  |  |  |  |  |  |  |  |  |  |
|  | test | data structure | WT | Het | KO | genotype |  |  | cohort |  |  | genotype x cohort |  |  | pairwise comparisons |  |  |
|  |  |  |  |  |  | F | p-value | power | F | p-value | power | F | p-value | power | WT vs Het | WT vs KO | Het vs KO |
| Pinna reflex | 2-way ANOVA | non normal | 0.89 ± 0.07 | 0.68 ± 0.1 | 0.73 ± 0.1 | 1.790 | 0.1773 | 0.357 | 13.988 | 0.000 | 0.956 | 1.155 | 0.323 | 0.243 | - | - | - |
| Cornel reflex | 2-way ANOVA | non normal | 1.05 ± 0.05 | 0.94 ± 0.05 | 1.05 ± 0.05 | 1.518 | 0.2289 | 0.308 | 0.389 | 0.536 | 0.094 | 1.518 | 0.229 | 0.308 | - | - | - |
| Toe pinch retraction | 2-way ANOVA | non normal | 2.05 ± 0.37 | 2.36 ± 0.33 | 2.26 ± 0.43 | 0.144 | 0.8659 | 0.071 | 1.073 | 0.305 | 0.174 | 0.072 | 0.931 | 0.060 | - | - | - |
| Preyer reflex | 2-way ANOVA | non normal | 1.47 ± 0.15 | 1.36 ± 0.13 | 1.42 ± 0.17 | 0.135 | 0.8740 | 0.070 | 22.250 | 0.000 | 0.996 | 1.478 | 0.238 | 0.301 | - | - | - |
| Visual Placing/Reaching reflex | NA | NA | 9 ± 0 | 9 ± 0 | 9 ± 0 | - | - | - | - | - | - | - | - | - | - | - | - |

| Tail flick |  |  |  |  |  |  |  |  |  |  |  |
| --- | --- | --- | --- | --- | --- | --- | --- | --- | --- | --- | --- |
|  | Cohorts 1 and 2 |  |  |  |  |  |  |  |  |  |  |
|  | test | data structure |  |  |  | F | p-value | power | WT vs Het | WT vs KO | Het vs KO |
| Latency, repeated measures |  |  |  |  |  |  |  |  |  |  |  |
| - trial effect | repeated measures | sphericity violated |  |  |  | 3.081 | 0.0500 | 0.583 | - | - | - |
| - trial x genotype effect | repeated measures | sphericity violated |  |  |  | 0.169 | 0.9535 | 0.085 | - | - | - |
| - genotype effect | repeated measures | sphericity violated |  |  |  | 1.118 | 0.3347 | 0.236 | - | - | - |
| - cohort effect | repeated measures | sphericity violated |  |  |  | 83.467 | 0.0000 | 1.000 | - | - | - |
| - trial x genotype x cohort effect | repeated measures | sphericity violated |  |  |  | 0.489 | 0.7438 | 0.162 | - | - | - |
| - genotype x cohort effect | repeated measures | sphericity violated |  |  |  | 2.162 | 0.1255 | 0.422 | - | - | - |

| Individual trials | test | data structure | WT | Het | KO | genotype |  |  | cohort |  |  | genotype x cohort |  |  | pairwise comparisons |  |  |
| --- | --- | --- | --- | --- | --- | --- | --- | --- | --- | --- | --- | --- | --- | --- | --- | --- | --- |
|  |  |  |  |  |  | F | p-value | power | F | p-value | power | F | p-value | power | WT vs Het | WT vs KO | Het vs KO |
| Latency to flick, trial 1 (seconds) | 2-way ANOVA | normal | 11.39 ± 0.97 | 11.32 ± 0.78 | 10.28 ± 0.82 | 0.356 | 0.7023 | 0.104 | 13.680 | 0.001 | 0.952 | 0.145 | 0.865 | 0.071 | - | - | - |
| Latency to flick, trial 2 (seconds) | 2-way ANOVA | normal | 10.99 ± 1.08 | 9.67 ± 0.98 | 8.86 ± 0.82 | 0.961 | 0.3892 | 0.208 | 36.614 | 0.000 | 1.000 | 2.286 | 0.112 | 0.444 | - | - | - |
| Latency to flick, trial 3 (seconds) | 2-way ANOVA | normal | 11.05 ± 1.23 | 10.06 ± 1.01 | 9.29 ± 0.94 | 0.526 | 0.5942 | 0.132 | 91.329 | 0.000 | 1.000 | 1.578 | 0.216 | 0.319 | - | - | - |
| Shortest latency to flick (seconds) | 2-way ANOVA | normal | 8.03 ± 0.82 | 7.99 ± 0.87 | 7.47 ± 0.78 | 0.064 | 0.9379 | 0.059 | 42.964 | 0.000 | 1.000 | 1.039 | 0.361 | 0.222 | - | - | - |
| Longest latency to flick (seconds) | 2-way ANOVA | non normal | 14.17 ± 0.94 | 12.82 ± 0.75 | 11.91 ± 0.72 | 2.213 | 0.1198 | 0.431 | 64.815 | 0.000 | 1.000 | 1.659 | 0.200 | 0.334 | - | - | - |
| Mean latency to flick (seconds) | 2-way ANOVA | non normal | 11.14 ± 0.88 | 10.35 ± 0.78 | 9.48 ± 0.72 | 1.118 | 0.3347 | 0.236 | 83.467 | 0.000 | 1.000 | 2.162 | 0.126 | 0.422 | - | - | - |

| Startle response |  |  |  |  |  |  |  |  |  |  |  |  |  |  |  |  |  |
| --- | --- | --- | --- | --- | --- | --- | --- | --- | --- | --- | --- | --- | --- | --- | --- | --- | --- |
|  | Cohorts 1 and 2 |  |  |  |  |  |  |  |  |  |  |  |  |  |  |  |  |
|  | test | data structure | WT | Het | KO | genotype |  |  | cohort |  |  | genotype x cohort |  |  | pairwise comparisons |  |  |
|  |  |  |  |  |  | F | p-value | power | F | p-value | power | F | p-value | power | WT vs Het | WT vs KO | Het vs KO |
| Startle response at 74 dB | 2-way ANOVA | non normal | 179.96 ± 20.24 | 165.57 ± 17.22 | 158.01 ± 10.36 | 2.376 | 0.1077 | 0.448 | 9.123 | 0.005 | 0.836 | 2.255 | 0.120 | 0.428 | - | - | - |
| Startle response at 78 dB | 2-way ANOVA | non normal | 183.48 ± 22 | 157.4 ± 16.93 | 160.69 ± 14.89 | 3.181 | 0.0538 | 0.571 | 5.779 | 0.022 | 0.647 | 3.673 | 0.036 | 0.638 | 0.046 | 0.025 | 0.770 |
| Startle response at 82 dB | 2-way ANOVA | non normal | 197.45 ± 26.03 | 160.09 ± 14.07 | 175.58 ± 13.87 | 3.254 | 0.0500 | 0.582 | 3.939 | 0.055 | 0.488 | 2.621 | 0.087 | 0.487 | 0.019 | 0.063 | 0.591 |
| Startle response at 86 dB | 2-way ANOVA | non normal | 246.33 ± 35.36 | 162.3 ± 15.61 | 176.15 ± 12.88 | 3.255 | 0.0500 | 0.582 | 0.082 | 0.777 | 0.059 | 0.271 | 0.764 | 0.089 | 0.024 | 0.042 | 0.812 |
| Startle response at 92 dB | 2-way ANOVA | non normal | 257.25 ± 40.4 | 192.63 ± 19.03 | 201.41 ± 23.01 | 2.153 | 0.1313 | 0.411 | 0.039 | 0.845 | 0.054 | 1.323 | 0.279 | 0.267 | - | - | - |

| Startle response, repeated measures | Cohorts 1 and 2 |  |  |  |  |  |  |  |  |  |  |
| --- | --- | --- | --- | --- | --- | --- | --- | --- | --- | --- | --- |
|  | test | data structure |  |  |  | F | p-value | power | WT vs Het | WT vs KO | Het vs KO |
| - sound intensity effect | repeated measures | sphericity assumed |  |  |  | 2.900 | 0.0510 | 0.605 | - | - | - |
| - sound intensity x genotype effect | repeated measures | sphericity assumed |  |  |  | 0.642 | 0.6620 | 0.219 | - | - | - |
| - genotype effect | repeated measures | sphericity assumed |  |  |  | 3.649 | 0.0364 | 0.635 | 0.022 | 0.024 | 0.989 |
| - cohort effect | repeated measures | sphericity assumed |  |  |  | 1.842 | 0.1835 | 0.262 | - | - | - |
| - sound intensity x genotype x cohort effect | repeated measures | sphericity assumed |  |  |  | 0.822 | 0.5335 | 0.276 | - | - | - |
| - genotype x cohort effect | repeated measures | sphericity assumed |  |  |  | 1.922 | 0.1614 | 0.372 | - | - | - |

|  | test | data structure | WT | Het | KO | genotype |  |  | cohort |  |  | genotype x cohort |  |  | pairwise comparisons |  |  |
| --- | --- | --- | --- | --- | --- | --- | --- | --- | --- | --- | --- | --- | --- | --- | --- | --- | --- |
|  |  |  |  |  |  | F | p-value | power | F | p-value | power | F | p-value | power | WT vs Het | WT vs KO | Het vs KO |
| Startle response at 74 dB, normalized to weight | 2-way ANOVA | non normal | 7.01 ± 0.82 | 6.18 ± 0.64 | 6.17 ± 0.32 | 2.843 | 0.0718 | 0.522 | 5.749 | 0.022 | 0.645 | 2.780 | 0.076 | 0.512 | 0.045 | 0.042 | 0.959 |
| Startle response at 78 dB, normalized to weight | 2-way ANOVA | non normal | 7.19 ± 0.89 | 5.86 ± 0.64 | 6.35 ± 0.58 | 3.051 | 0.0601 | 0.553 | 2.972 | 0.094 | 0.389 | 3.611 | 0.038 | 0.630 | 0.038 | 0.036 | 0.964 |
| Startle response at 82 dB, normalized to weight | 2-way ANOVA | non normal | 7.74 ± 1.06 | 5.89 ± 0.41 | 6.98 ± 0.63 | 3.378 | 0.0456 | 0.599 | 1.700 | 0.201 | 0.245 | 2.365 | 0.109 | 0.446 | 0.015 | 0.089 | 0.409 |
| Startle response at 86 dB, normalized to weight | 2-way ANOVA | non normal | 9.86 ± 1.66 | 6 ± 0.49 | 7.03 ± 0.64 | 2.888 | 0.0690 | 0.529 | 0.030 | 0.864 | 0.053 | 0.268 | 0.766 | 0.089 | 0.028 | 0.070 | 0.677 |
| Startle response at 92 dB, normalized to weight | 2-way ANOVA | non normal | 10.04 ± 1.77 | 7.09 ± 0.58 | 8.18 ± 1.15 | 1.758 | 0.1873 | 0.343 | 0.323 | 0.573 | 0.086 | 1.291 | 0.288 | 0.261 | - | - | - |

| Startle response normalized, repeated measures | test | data structure |  |  |  | F | p-value | power | WT vs Het | WT vs KO | Het vs KO |
| --- | --- | --- | --- | --- | --- | --- | --- | --- | --- | --- | --- |
| - sound intensity effect | repeated measures | sphericity assumed |  |  |  | 2.506 | 0.0449 | 0.248 | - | - | - |
| - sound intensity x genotype effect | repeated measures | sphericity assumed |  |  |  | 0.580 | 0.7933 | 0.098 | - | - | - |
| - genotype effect | repeated measures | sphericity assumed |  |  |  | 3.338 | 0.0471 | 0.593 | 0.127 | 0.407 | 0.783 |
| - cohort effect | repeated measures | sphericity assumed |  |  |  | 0.455 | 0.5046 | 0.101 | - | - | - |
| - sound intensity x genotype x cohort effect | repeated measures | sphericity assumed |  |  |  | 0.716 | 0.5922 | 0.079 | - | - | - |
| - genotype x cohort effect | repeated measures | sphericity assumed |  |  |  | 1.812 | 0.1783 | 0.353 | - | - | - |

Pre-pulse inhibition
|  | test | data structure | WT | Het | KO | Cohorts 1 and 2 |  |  |  |  |  |  |  |  |  |  |  |
| --- | --- | --- | --- | --- | --- | --- | --- | --- | --- | --- | --- | --- | --- | --- | --- | --- | --- |
|  |  |  |  |  |  | genotype |  |  | cohort |  |  | genotype x cohort |  |  | pairwise comparisons |  |  |
|  |  |  |  |  |  | F | p-value | power | F | p-value | power | F | p-value | power | WT vs Het | WT vs KO | Het vs KO |
| Percentage of inhibition at 74 dB | 2-way ANOVA | normal | 22.47 ± 5.7 | 15.1 ± 4.72 | 10.56 ± 6.03 | 0.625 | 0.5409 | 0.212 | 11.061 | 0.002 | 0.519 | 1.821 | 0.177 | 0.190 | - | - | - |
| Percentage of inhibition at 78 dB | 2-way ANOVA | normal | 31.81 ± 4.48 | 14.84 ± 6.7 | 21.18 ± 5.03 | 3.513 | 0.0407 | 0.543 | 21.819 | 0.000 | 0.793 | 1.008 | 0.375 | 0.352 | 0.058 | 0.312 | 0.656 |
| Percentage of inhibition at 82 dB | 2-way ANOVA | normal | 30.39 ± 5.51 | 21.53 ± 6.08 | 13.34 ± 5.99 | 0.151 | 0.8604 | 0.426 | 14.892 | 0.000 | 0.480 | 0.042 | 0.959 | 0.348 | - | - | - |
| Percentage of inhibition at 86 dB | 2-way ANOVA | non normal | 31.73 ± 7.95 | 21.04 ± 7.37 | 27.04 ± 4.63 | 1.411 | 0.2575 | 0.176 | 37.139 | 0.000 | 0.909 | 0.463 | 0.633 | 0.217 | - | - | - |
| Percentage of inhibition at 92 dB | 2-way ANOVA | non normal | 43.62 ± 7.01 | 34.35 ± 5.82 | 29.93 ± 6.97 | 0.989 | 0.3821 | 0.212 | 54.683 | 0.000 | 0.892 | 2.258 | 0.120 | 0.222 | - | - | - |
| Percentage of inhibition, mean | 2-way ANOVA | normal | 32 ± 5.53 | 21.37 ± 5.48 | 20.41 ± 5.03 | 0.155 | 0.8569 | 0.305 | 37.489 | 0.000 | 0.867 | 0.310 | 0.735 | 0.278 | - | - | - |

| Percentage of inhibition, repeated measure | test | data structure |  |  |  | F | p-value | power | WT vs Het | WT vs KO | Het vs KO |
| --- | --- | --- | --- | --- | --- | --- | --- | --- | --- | --- | --- |
| - sound intensity effect | repeated measures | sphericity violated |  |  |  | 15.396 | 0.0000 | 1.000 | - | - | - |
| - sound intensity x genotype effect | repeated measures | sphericity violated |  |  |  | 1.409 | 0.1943 | 0.603 | - | - | - |
| - genotype effect | repeated measures | sphericity violated |  |  |  | 1.502 | 0.2325 | 0.305 | - | - | - |
| - cohort effect | repeated measures | sphericity violated |  |  |  | 9.806 | 0.0029 | 0.867 | - | - | - |
| - sound intensity x genotype x cohort effect | repeated measures | sphericity violated |  |  |  | 0.978 | 0.4540 | 0.427 | - | - | - |
| - genotype x cohort effect | repeated measures | sphericity violated |  |  |  | 1.349 | 0.2686 | 0.278 | - | - | - |

Buried food test
|  | test | data structure | WT | Het | KO | Cohorts 1 and 2 |  |  |  |  |  |  |  |  |  |  |  |
| --- | --- | --- | --- | --- | --- | --- | --- | --- | --- | --- | --- | --- | --- | --- | --- | --- | --- |
|  |  |  |  |  |  | genotype |  |  | cohort |  |  | genotype x cohort |  |  | pairwise comparisons |  |  |
|  |  |  |  |  |  | F | p-value | power | F | p-value | power | F | p-value | power | WT vs Het | WT vs KO | Het vs KO |
| Latency to retrieve and eat food (seconds) | 2-way ANOVA | non normal | 51.11 ± 9.9 | 94.81 ± 23.69 | 500.28 ± 94.92 | 17.848 | 0.0000 | 1.000 | 0.001 | 0.976 | 0.050 | 0.000 | 1.000 | 0.050 | 0.858 | 0.000 | 0.000 |

Olfactory habituation/dishabituation, sniffing only
|  | test | data structure | Cohorts 1 and 2 |  |  |  |  |  |
| --- | --- | --- | --- | --- | --- | --- | --- | --- |
|  |  |  | F | p-value | power | WT vs Het | WT vs KO | Het vs KO |
| Water repeated measure |  |  |  |  |  |  |  |  |
| - trial effect | repeated measures | sphericity violated | 8.290 | 0.0019 | 0.958 | - | - | - |
| - trial x genotype effect | repeated measures | sphericity violated | 1.108 | 0.3505 | 0.337 | - | - | - |
| - genotype effect | repeated measures | sphericity violated | 2.973 | 0.0602 | 0.553 | 0.660 | 0.027 | 0.066 |
| - cohort effect | repeated measures | sphericity violated | 0.073 | 0.7886 | 0.058 | - | - | - |
| - trial x genotype x cohort effect | repeated measures | sphericity violated | 1.739 | 0.1683 | 0.515 | - | - | - |
| - genotype x cohort effect | repeated measures | sphericity violated | 1.015 | 0.3696 | 0.217 | - | - | - |

| Banana repeated measure | test | data structure |  |  |  | F | p-value | power | WT vs Het | WT vs KO | Het vs KO |
| --- | --- | --- | --- | --- | --- | --- | --- | --- | --- | --- | --- |
| - trial effect | repeated measures | sphericity assumed |  |  |  | 10.117 | 0.0001 | 0.983 | - | - | - |

|  |  |  |  |  |  |  |  |  |  |
| --- | --- | --- | --- | --- | --- | --- | --- | --- | --- |
| - trial x genotype effect | repeated measures | sphericity assumed |  | 3.908 | **0.0054** | 0.888 | - | - | - |
| - genotype effect | repeated measures | sphericity assumed |  | 5.681 | **0.0060** | 0.842 | 0.433 | **0.002** | **0.017** |
| - cohort effect | repeated measures | sphericity assumed |  | 11.933 | **0.0011** | 0.923 | - | - | - |
| - trial x genotype x cohort effect | repeated measures | sphericity assumed |  | 0.486 | 0.7461 | 0.161 | - | - | - |
| - genotype x cohort effect | repeated measures | sphericity assumed |  | 0.134 | 0.8752 | 0.069 | - | - | - |

| Lemon repeated measure | test | data structure |  | F | p-value | power | WT vs Het | WT vs KO | Het vs KO |
| --- | --- | --- | --- | --- | --- | --- | --- | --- | --- |
| - trial effect | repeated measures | sphericity violated |  | 6.699 | **0.0041** | 0.908 | - | - | - |
| - trial x genotype effect | repeated measures | sphericity violated |  | 0.047 | 0.9890 | 0.059 | - | - | - |
| - genotype effect | repeated measures | sphericity violated |  | 2.715 | **0.0760** | 0.513 | 0.404 | 0.166 | **0.025** |
| - cohort effect | repeated measures | sphericity violated |  | 15.327 | **0.0003** | 0.970 | - | - | - |
| - trial x genotype x cohort effect | repeated measures | sphericity violated |  | 0.667 | 0.5828 | 0.211 | - | - | - |
| - genotype x cohort effect | repeated measures | sphericity violated |  | 0.159 | 0.8534 | 0.073 | - | - | - |

| Male repeated measure | test | data structure |  | F | p-value | power | WT vs Het | WT vs KO | Het vs KO |
| --- | --- | --- | --- | --- | --- | --- | --- | --- | --- |
| - trial effect | repeated measures | sphericity violated |  | 27.903 | **0.0000** | 1.000 | - | - | - |
| - trial x genotype effect | repeated measures | sphericity violated |  | 1.089 | 0.3581 | 0.332 | - | - | - |
| - genotype effect | repeated measures | sphericity violated |  | 0.739 | 0.4828 | 0.168 | - | - | - |
| - cohort effect | repeated measures | sphericity violated |  | 2.500 | 0.1201 | 0.341 | - | - | - |
| - trial x genotype x cohort effect | repeated measures | sphericity violated |  | 1.603 | 0.1976 | 0.479 | - | - | - |
| - genotype x cohort effect | repeated measures | sphericity violated |  | 0.541 | 0.5857 | 0.134 | - | - | - |

| Female repeated measure | test | data structure |  | F | p-value | power | WT vs Het | WT vs KO | Het vs KO |
| --- | --- | --- | --- | --- | --- | --- | --- | --- | --- |
| - trial effect | repeated measures | sphericity violated |  | 20.922 | **0.0000** | 1.000 | - | - | - |
| - trial x genotype effect | repeated measures | sphericity violated |  | 0.131 | 0.9321 | 0.076 | - | - | - |
| - genotype effect | repeated measures | sphericity violated |  | 0.585 | 0.5609 | 0.142 | - | - | - |
| - cohort effect | repeated measures | sphericity violated |  | 14.771 | **0.0003** | 0.965 | - | - | - |
| - trial x genotype x cohort effect | repeated measures | sphericity violated |  | 0.348 | 0.7765 | 0.126 | - | - | - |
| - genotype x cohort effect | repeated measures | sphericity violated |  | 1.651 | 0.2022 | 0.332 | - | - | - |

| Individual trials | test | data structure | WT | Het | KO | genotype |  |  | cohort |  |  | genotype x cohort |  |  | pairwise comparisons |  |  |
| --- | --- | --- | --- | --- | --- | --- | --- | --- | --- | --- | --- | --- | --- | --- | --- | --- | --- |
|  |  |  |  |  |  | F | p-value | power | F | p-value | power | F | p-value | power | WT vs Het | WT vs KO | Het vs KO |
| Water 1 | 2-way ANOVA | non normal | 1.82 ± 0.37 | 1.81 ± 0.42 | 0.9 ± 0.16 | 2.205 | 0.1208 | 0.429 | 0.199 | 0.658 | 0.072 | 1.697 | 0.194 | 0.340 | - | - | - |
| Water 2 | 2-way ANOVA | non normal | 1.02 ± 0.15 | 0.98 ± 0.14 | 0.7 ± 0.15 | 1.438 | 0.2470 | 0.294 | 0.078 | 0.781 | 0.059 | 0.711 | 0.496 | 0.163 | - | - | - |
| Water 3 | 2-way ANOVA | non normal | 0.97 ± 0.15 | 0.82 ± 0.14 | 0.7 ± 0.11 | 1.724 | 0.1888 | 0.345 | 2.900 | **0.095** | 0.386 | 0.366 | 0.696 | 0.106 | - | - | - |
| Banana 1 | 2-way ANOVA | non normal | 1.34 ± 0.12 | 1.04 ± 0.23 | 0.44 ± 0.15 | 8.742 | **0.0006** | 0.961 | 6.201 | **0.016** | 0.685 | 0.466 | 0.630 | 0.122 | 0.322 | **0.001** | 0.052 |
| Banana 2 | 2-way ANOVA | non normal | 0.65 ± 0.12 | 0.65 ± 0.1 | 0.44 ± 0.13 | 1.641 | 0.2040 | 0.330 | 6.658 | **0.013** | 0.716 | 0.240 | 0.787 | 0.086 | - | - | - |
| Banana 3 | 2-way ANOVA | non normal | 0.52 ± 0.09 | 0.72 ± 0.19 | 0.44 ± 0.13 | 1.167 | 0.3196 | 0.245 | 7.761 | **0.008** | 0.780 | 0.050 | 0.951 | 0.057 | - | - | - |
| Lemon 1 | 2-way ANOVA | non normal | 0.58 ± 0.18 | 0.77 ± 0.15 | 0.52 ± 0.17 | 0.720 | 0.4916 | 0.165 | 8.572 | **0.005** | 0.819 | 0.041 | 0.960 | 0.056 | - | - | - |
| Lemon 2 | 2-way ANOVA | non normal | 0.39 ± 0.11 | 0.52 ± 0.1 | 0.24 ± 0.07 | 2.679 | **0.0785** | 0.507 | 9.784 | **0.003** | 0.866 | 0.037 | 0.964 | 0.055 | 0.709 | 0.411 | **0.099** |
| Lemon 3 | 2-way ANOVA | non normal | 0.35 ± 0.06 | 0.48 ± 0.09 | 0.25 ± 0.08 | 2.547 | **0.0885** | 0.486 | 7.405 | **0.009** | 0.761 | 2.493 | **0.093** | 0.478 | 0.594 | 0.540 | 0.106 |
| Male 1 | 2-way ANOVA | non normal | 6.1 ± 1.12 | 4.66 ± 0.88 | 4.94 ± 1.07 | 0.659 | 0.5219 | 0.154 | 1.095 | 0.300 | 0.177 | 1.091 | 0.344 | 0.231 | - | - | - |
| Male 2 | 2-way ANOVA | non normal | 2.25 ± 0.34 | 2.77 ± 0.46 | 1.26 ± 0.31 | 4.095 | **0.0225** | 0.700 | 0.564 | 0.456 | 0.114 | 3.078 | **0.055** | 0.568 | 0.752 | 0.116 | **0.020** |
| Male 3 | 2-way ANOVA | non normal | 1.54 ± 0.38 | 1.25 ± 0.24 | 1.72 ± 0.63 | 0.322 | 0.7264 | 0.098 | 2.465 | 0.123 | 0.337 | 0.312 | 0.733 | 0.097 | - | - | - |
| Female 1 | 2-way ANOVA | non normal | 5.68 ± 0.98 | 6.43 ± 1.34 | 6.32 ± 1.93 | 0.129 | 0.8790 | 0.069 | 11.509 | **0.001** | 0.914 | 0.879 | 0.422 | 0.193 | - | - | - |
| Female 2 | 2-way ANOVA | non normal | 2.18 ± 0.3 | 2.24 ± 0.43 | 2.62 ± 0.81 | 0.189 | 0.8283 | 0.078 | 1.936 | 0.170 | 0.276 | 0.289 | 0.751 | 0.093 | - | - | - |
| Female 3 | 2-way ANOVA | non normal | 1.42 ± 0.26 | 1.82 ± 0.52 | 2.76 ± 0.88 | 1.600 | 0.2120 | 0.323 | 7.672 | **0.008** | 0.775 | 1.792 | 0.177 | 0.357 | - | - | - |

| Olfactory habituation/dishabituation, all interactions |  |  |  |  |  |  |  |  |  |  |  |
| --- | --- | --- | --- | --- | --- | --- | --- | --- | --- | --- | --- |
| Cohorts 1 and 2 |  |  |  |  |  |  |  |  |  |  |  |
| Water repeated measure | test | data structure |  |  |  | F | p-value | power | WT vs Het | WT vs KO | Het vs KO |
| - trial effect | repeated measures | sphericity violated |  |  |  | 1.891 | 0.1664 | 0.385 | - | - | - |
| - trial x genotype effect | repeated measures | sphericity violated |  |  |  | 0.718 | 0.5478 | 0.225 | - | - | - |

|  |  |  |  |  |  |  |  |  |  |  |  |
| --- | --- | --- | --- | --- | --- | --- | --- | --- | --- | --- | --- |
| - genotype effect | repeated measures | sphericity violated |  |  |  | 7.210 | **0.0017** | 0.920 | 0.637 | **0.001** | **0.015** |
| - cohort effect | repeated measures | sphericity violated |  |  |  | 2.639 | 0.1104 | 0.357 | - | - | - |
| - trial x genotype x cohort effect | repeated measures | sphericity violated |  |  |  | 0.920 | 0.4371 | 0.283 | - | - | - |
| - genotype x cohort effect | repeated measures | sphericity violated |  |  |  | 1.133 | 0.3300 | 0.239 | - | - | - |

| Banana repeated measure | test | data structure |  |  |  | F | p-value | power | WT vs Het | WT vs KO | Het vs KO |
| --- | --- | --- | --- | --- | --- | --- | --- | --- | --- | --- | --- |
| - trial effect | repeated measures | sphericity violated |  |  |  | 5.229 | **0.0133** | 0.821 | - | - | - |
| - trial x genotype effect | repeated measures | sphericity violated |  |  |  | 1.282 | 0.2866 | 0.388 | - | - | - |
| - genotype effect | repeated measures | sphericity violated |  |  |  | 5.737 | **0.0057** | 0.846 | 0.744 | **0.060** | **0.008** |
| - cohort effect | repeated measures | sphericity violated |  |  |  | 7.922 | **0.0070** | 0.788 | - | - | - |
| - trial x genotype x cohort effect | repeated measures | sphericity violated |  |  |  | 0.936 | 0.4284 | 0.287 | - | - | - |
| - genotype x cohort effect | repeated measures | sphericity violated |  |  |  | 0.990 | 0.3787 | 0.213 | - | - | - |

| Lemon repeated measure | test | data structure |  |  |  | F | p-value | power | WT vs Het | WT vs KO | Het vs KO |
| --- | --- | --- | --- | --- | --- | --- | --- | --- | --- | --- | --- |
| - trial effect | repeated measures | sphericity violated |  |  |  | 1.303 | 0.2728 | 0.276 | - | - | - |
| - trial x genotype effect | repeated measures | sphericity violated |  |  |  | 0.703 | 0.5597 | 0.221 | - | - | - |
| - genotype effect | repeated measures | sphericity violated |  |  |  | 4.893 | **0.0115** | 0.781 | 0.295 | **0.048** | **0.003** |
| - cohort effect | repeated measures | sphericity violated |  |  |  | 5.152 | **0.0276** | 0.605 | - | - | - |
| - trial x genotype x cohort effect | repeated measures | sphericity violated |  |  |  | 0.527 | 0.6738 | 0.172 | - | - | - |
| - genotype x cohort effect | repeated measures | sphericity violated |  |  |  | 2.405 | 0.1006 | 0.463 | - | - | - |

| Male repeated measure | test | data structure |  |  |  | F | p-value | power | WT vs Het | WT vs KO | Het vs KO |
| --- | --- | --- | --- | --- | --- | --- | --- | --- | --- | --- | --- |
| - trial effect | repeated measures | sphericity violated |  |  |  | 28.652 | **0.0000** | 1.000 | - | - | - |
| - trial x genotype effect | repeated measures | sphericity violated |  |  |  | 0.790 | 0.4993 | 0.246 | - | - | - |
| - genotype effect | repeated measures | sphericity violated |  |  |  | 5.722 | **0.0057** | 0.845 | 0.839 | **0.005** | **0.022** |
| - cohort effect | repeated measures | sphericity violated |  |  |  | 2.953 | **0.0918** | 0.392 | - | - | - |
| - trial x genotype x cohort effect | repeated measures | sphericity violated |  |  |  | 0.367 | 0.7697 | 0.131 | - | - | - |
| - genotype x cohort effect | repeated measures | sphericity violated |  |  |  | 0.009 | 0.9906 | 0.051 | - | - | - |

| Female repeated measure | test | data structure |  |  |  | F | p-value | power | WT vs Het | WT vs KO | Het vs KO |
| --- | --- | --- | --- | --- | --- | --- | --- | --- | --- | --- | --- |
| - trial effect | repeated measures | sphericity assumed |  |  |  | 25.044 | **0.0000** | 1.000 | - | - | - |
| - trial x genotype effect | repeated measures | sphericity assumed |  |  |  | 0.884 | 0.4762 | 0.264 | - | - | - |
| - genotype effect | repeated measures | sphericity assumed |  |  |  | 1.119 | 0.3346 | 0.236 | - | - | - |
| - cohort effect | repeated measures | sphericity assumed |  |  |  | 6.400 | **0.0145** | 0.699 | - | - | - |
| - trial x genotype x cohort effect | repeated measures | sphericity assumed |  |  |  | 1.911 | 0.1143 | 0.541 | - | - | - |
| - genotype x cohort effect | repeated measures | sphericity assumed |  |  |  | 0.431 | 0.6521 | 0.116 | - | - | - |

| Individual trials | test | data structure | WT | Het | KO | genotype |  |  | cohort |  |  | genotype x cohort |  |  | pairwise comparisons |  |  |
| --- | --- | --- | --- | --- | --- | --- | --- | --- | --- | --- | --- | --- | --- | --- | --- | --- | --- |
|  |  |  |  |  |  | F | p-value | power | F | p-value | power | F | p-value | power | WT vs Het | WT vs KO | Het vs KO |
| Water 1 | 2-way ANOVA | non normal | 3.52 ± 0.9 | 3.88 ± 1 | 1.35 ± 0.27 | 2.357 | 0.1053 | 0.455 | 3.302 | **0.075** | 0.429 | 1.116 | 0.336 | 0.235 | - | - | - |
| Water 2 | 2-way ANOVA | non normal | 3.45 ± 0.71 | 1.78 ± 0.48 | 0.95 ± 0.23 | 4.964 | **0.0109** | 0.786 | 0.588 | 0.447 | 0.117 | 1.295 | 0.283 | 0.267 | 0.161 | **0.006** | 0.364 |
| Water 3 | 2-way ANOVA | non normal | 2.53 ± 0.54 | 2.3 ± 0.6 | 0.95 ± 0.2 | 2.997 | **0.0592** | 0.556 | 0.033 | 0.856 | 0.054 | 0.373 | 0.690 | 0.107 | 0.949 | **0.078** | 0.147 |
| Banana 1 | 2-way ANOVA | non normal | 2.8 ± 0.42 | 3.38 ± 1.16 | 0.72 ± 0.43 | 3.854 | **0.0279** | 0.672 | 6.603 | **0.013** | 0.712 | 1.080 | 0.348 | 0.229 | - | - | - |
| Banana 2 | 2-way ANOVA | non normal | 1.46 ± 0.49 | 1.06 ± 0.35 | 0.57 ± 0.18 | 1.902 | 0.1601 | 0.376 | 4.991 | **0.030** | 0.591 | 1.056 | 0.355 | 0.224 | - | - | - |
| Banana 3 | 2-way ANOVA | non normal | 1.38 ± 0.55 | 2 ± 0.63 | 0.4 ± 0.1 | 3.927 | **0.0262** | 0.680 | 1.920 | 0.172 | 0.274 | 1.097 | 0.342 | 0.232 | 0.334 | 0.380 | **0.020** |
| Lemon 1 | 2-way ANOVA | non normal | 1.47 ± 0.37 | 2.21 ± 0.74 | 0.45 ± 0.19 | 3.707 | **0.0317** | 0.653 | 2.710 | 0.106 | 0.365 | 0.878 | 0.422 | 0.193 | 0.412 | 0.339 | **0.024** |
| Lemon 2 | 2-way ANOVA | non normal | 1.21 ± 0.35 | 0.98 ± 0.36 | 0.28 ± 0.09 | 2.536 | **0.0895** | 0.484 | 3.609 | **0.063** | 0.461 | 1.012 | 0.371 | 0.216 | 0.781 | **0.091** | 0.315 |
| Lemon 3 | 2-way ANOVA | non normal | 0.8 ± 0.18 | 0.61 ± 0.15 | 0.2 ± 0.08 | 1.362 | 0.2656 | 0.280 | 1.615 | 0.210 | 0.238 | 1.334 | 0.273 | 0.275 | - | - | - |
| Male 1 | 2-way ANOVA | non normal | 55.55 ± 6.73 | 48.98 ± 7.22 | 27.15 ± 6.11 | 3.341 | **0.0436** | 0.605 | 2.419 | 0.126 | 0.332 | 0.054 | 0.948 | 0.058 | 0.655 | **0.030** | 0.199 |
| Male 2 | 2-way ANOVA | non normal | 27.09 ± 6.01 | 29.19 ± 5.33 | 7.48 ± 4.34 | 4.753 | **0.0130** | 0.768 | 1.196 | 0.280 | 0.189 | 0.012 | 0.988 | 0.052 | 0.986 | **0.020** | **0.030** |
| Male 3 | 2-way ANOVA | non normal | 17.57 ± 5.5 | 14.39 ± 4.43 | 5.26 ± 1.63 | 2.323 | 0.1087 | 0.449 | 1.329 | 0.255 | 0.204 | 0.978 | 0.383 | 0.210 | - | - | - |
| Female 1 | 2-way ANOVA | non normal | 53.65 ± 5.97 | 50.59 ± 5.48 | 31.16 ± 7.21 | 1.781 | 0.1792 | 0.355 | 6.118 | **0.017** | 0.679 | 1.565 | 0.219 | 0.316 | - | - | - |
| Female 2 | 2-way ANOVA | non normal | 23.26 ± 6.37 | 18.85 ± 6.77 | 14.61 ± 4.3 | 0.427 | 0.6551 | 0.115 | 2.116 | 0.152 | 0.297 | 0.350 | 0.707 | 0.103 | - | - | - |
| Female 3 | 2-way ANOVA | non normal | 14.57 ± 3.85 | 11.5 ± 3.69 | 14.21 ± 4.61 | 0.269 | 0.7650 | 0.090 | 0.575 | 0.452 | 0.115 | 1.703 | 0.193 | 0.341 | - | - | - |

No genotype differences were detected in tactile tests including the pinna reflex, the palpebral reflex and the toe pinch retraction test. In the tail flick pain sensitivity test, a trend toward a decreased latency to flick the tail in response to a noxious thermal stimulation a was observed in *Shank3^Δ4-22^* homozygous animals (Figure 4A).

**Figure 4:**
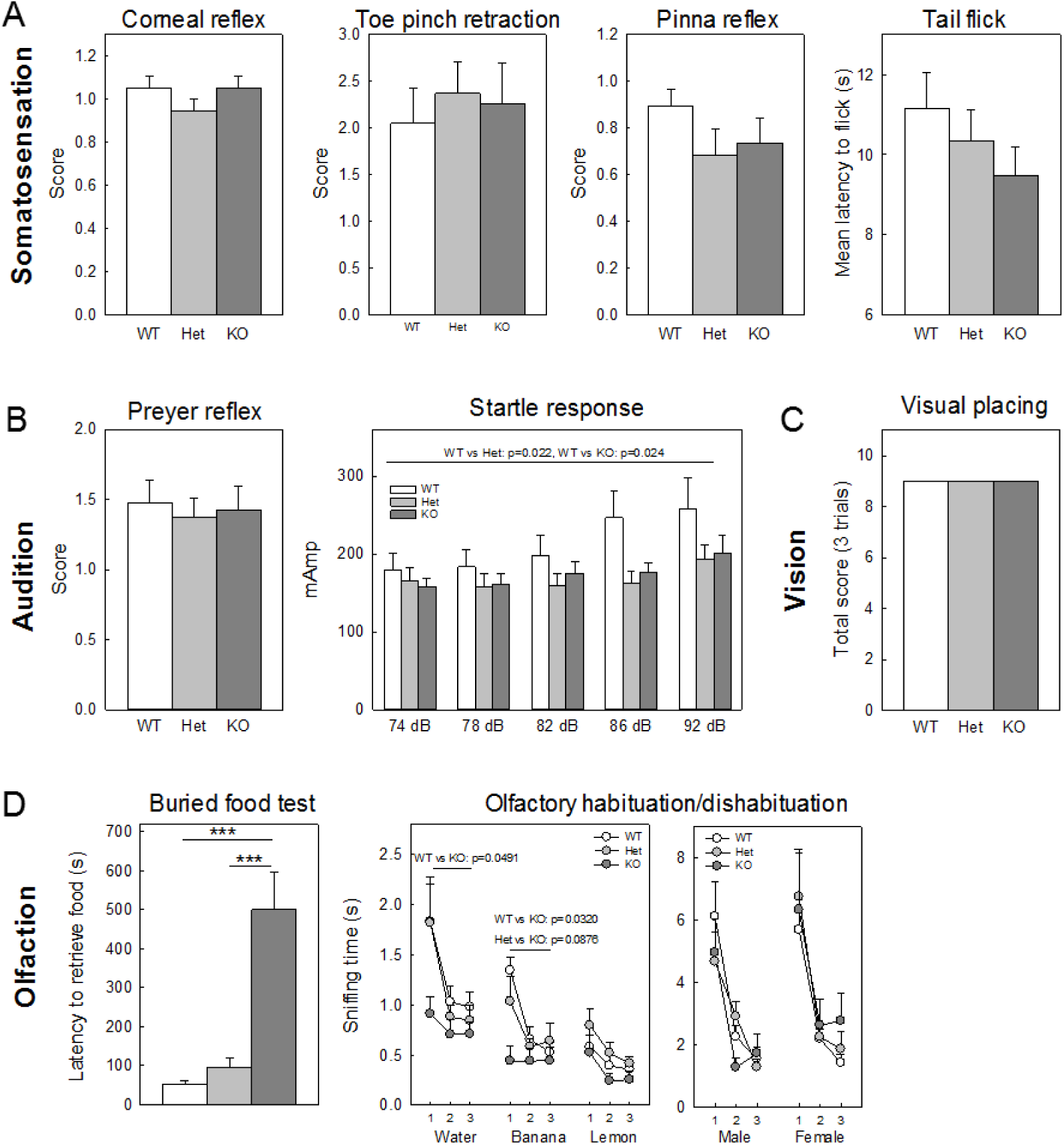
Altered sensory profile in *Shank3^Δ4-22^*-deficient mice. **(A)** Somatosensation evaluated with corneal reflex, toe pinch retraction, pinna reflex and tail flick. Normal tactile and pain responses were observed in *Shank3^Δ4-22^-*deficient mice. **(B)** Auditory functions measured with the Preyer reflex and startle response to increasing sound intensities. No genotype difference was observed for Preyer reflex however a startle response was decreased in both heterozygous and homozygous *Shank3^Δ4-22^* mice compared to their wild-type littermate with genotype differences being more marked for the higher startle intensities. Pre-pulse inhibition results are displayed in Extended Figure 4-1 A. **(C)** Gross visual function assessed by the visual placing test. Normal visual placing was observed for all genotypes. **(D)** Olfactory abilities evaluated by the time to find hidden food in buried food test and the cumulative time sniffing the applicator without direct interactions during olfactory habituation and dishabituation to nonsocial and social odors. Strong impairments were observed in the buried food test for *Shank3^Δ4-22^* homozygous mice as shown by a significant increase of the latency to retrieve the buried food compared to their heterozygous and wild-type littermates. Individual performances are available in Extended Figure 4-1 B. Similarly, a significant lack of interest for non-social scents (water, banana and lemon) was observed in *Shank3^Δ4-22^* homozygous mice but not in heterozygotes and wild-type during olfactory habituation/dishabituation while they still displayed normal habituation/dishabituation for social scents (unfamiliar male and female bedding). The olfactory habituation and dishabituation to nonsocial and social odors was measured as cumulative time spent sniffing a sequence of identical and novel odors delivered on cotton swabs inserted into a clean cage. WT, wild-type mice; Het, heterozygous mice; KO, homozygous knockout mice. *: p<0.05, **: p<0.1, ***: p<0.001.

Normal Preyer reflexes were observed in all genotypes, however, *Shank3^Δ4-22^* heterozygous and homozygous mice showed a reduced startle response throughout all the sound intensities (74 to 92 dB, analyzed as repeated measures) indicating an impaired sound discrimination (Figure 4B). Changes in pre-pulse inhibition of acoustic startle in *Shank3^Δ4-22^-* deficient mice are consistent with abnormalities in auditory processing, rather than sensorimotor gating deficits (Extended Figure 4-1 A).

Normal visual placing/reaching reflexes were observed for all the mice, thus ruling out strong visual impairments (Figure 4C).

*Shank3^Δ4-22^* homozygous mice demonstrated strong deficits in the buried food test (Figure 4D, left panel) with only 7 out of 19 mice able to retrieve the food in less than two minutes and 9 out of 19 mice not being able to find the food at all (Extended Figure 4-1 B). However, all animals showed interest for the food and ate it when it was made visible. To further investigate olfactory function, animals were subjected to the olfactory habituation/dishabituation paradigm using three non-social scents (water, banana and lemon) and two social scents (unfamiliar males and unfamiliar females). Wild-type and *Shank3^Δ4-22^* heterozygous animals displayed a normal response, characterized by a robust sniffing elicited by the first scent presentation of each non-social and social scent that declined over the second and third presentation of the same scent. In contrast, *Shank3^Δ4-22^* homozygous animals had little response to any of the non-social scents, even upon their first presentation (Figure 4D, middle panel), thus confirming the results of the buried food test. Interestingly the lack of interest for olfactory stimuli does not appear to be the consequence of anosmia as a normal response to both social scents was observed in *Shank3^Δ4-22^* homozygous mice (Figure 4D, right panel).

### Social interactions in *Shank3^Δ4-22^*-deficient mice

Mice were evaluated for social abilities during male-female dyadic social interaction, in the 3-chambered social interaction task, and in the social transmission of food preference test and detailed results are reported in Table 8. In freely moving male-female dyads of male mice paired with unfamiliar wild-type estrous C57BL6 females, sniffing time was generally similar across genotypes (Figure 5A, left panel). A significant increase for the first event of anogenital sniffing was found in male *Shank3^Δ4-22^* homozygous mice (Figure 5A, right panel) and we can note that this latency may contribute to trend towards reduced anogenital sniffing time in those animals. Ultrasonic vocalizations did not show significant difference across genotypes (Extended Figure 5-1 A).

**Table 8:**
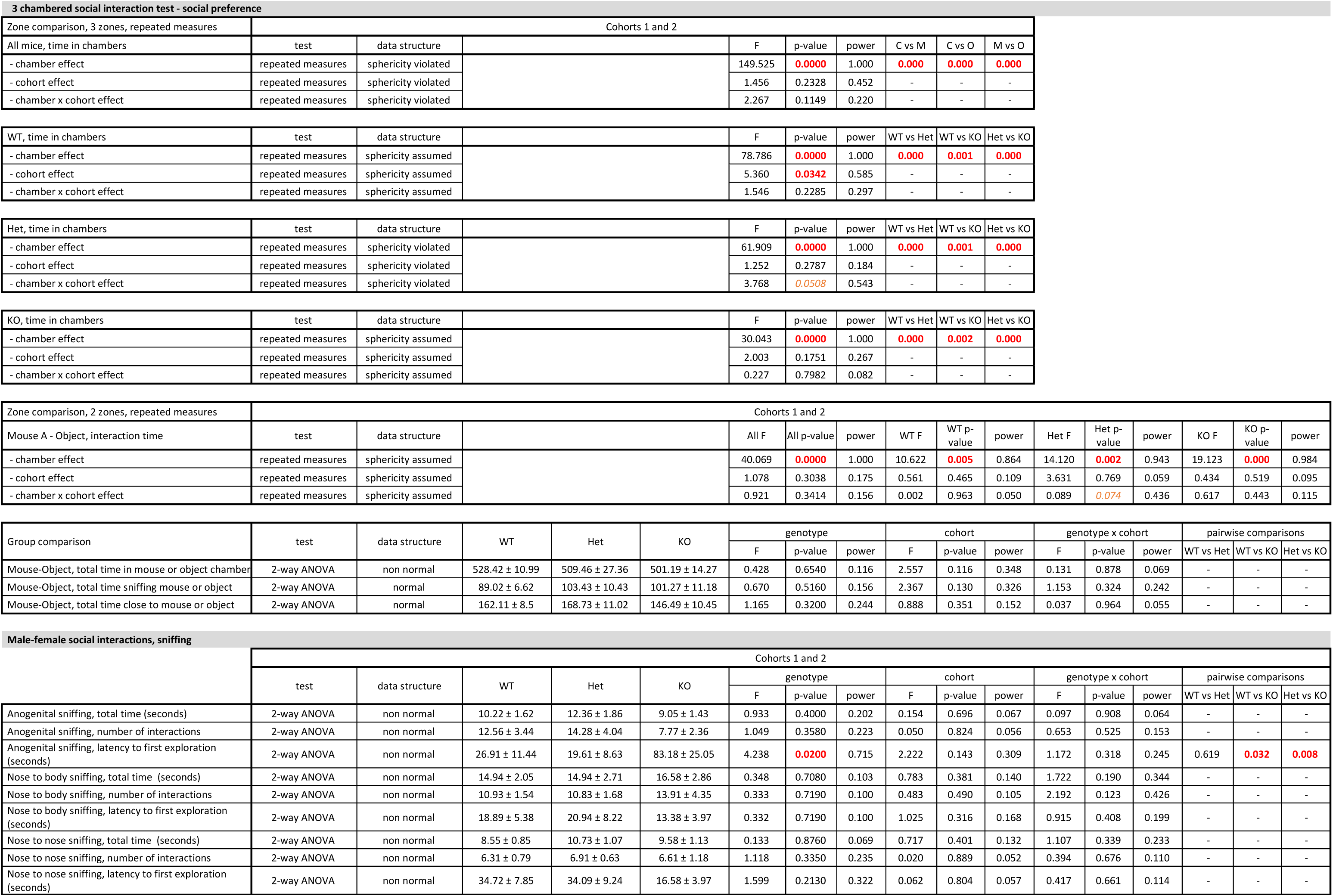
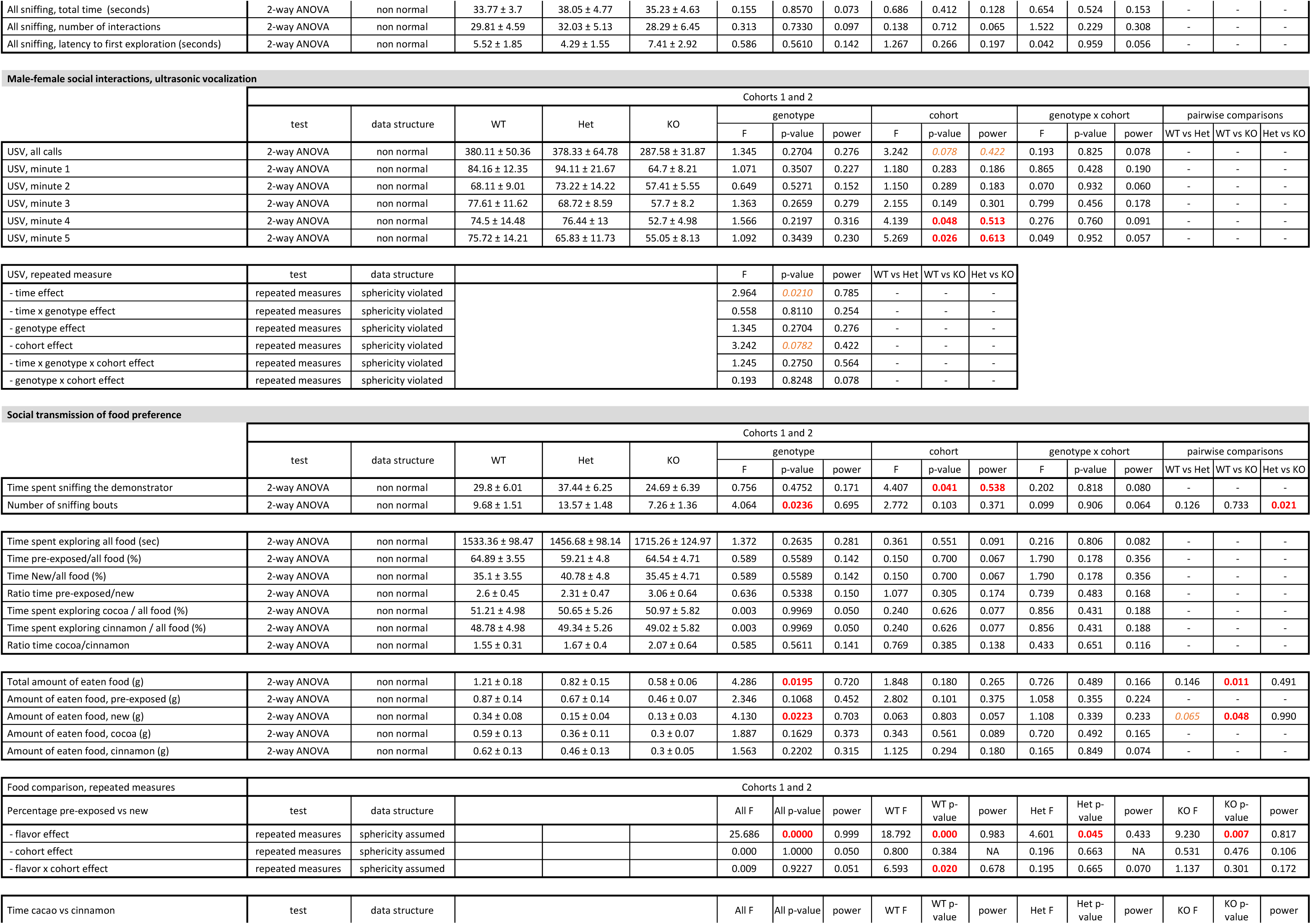
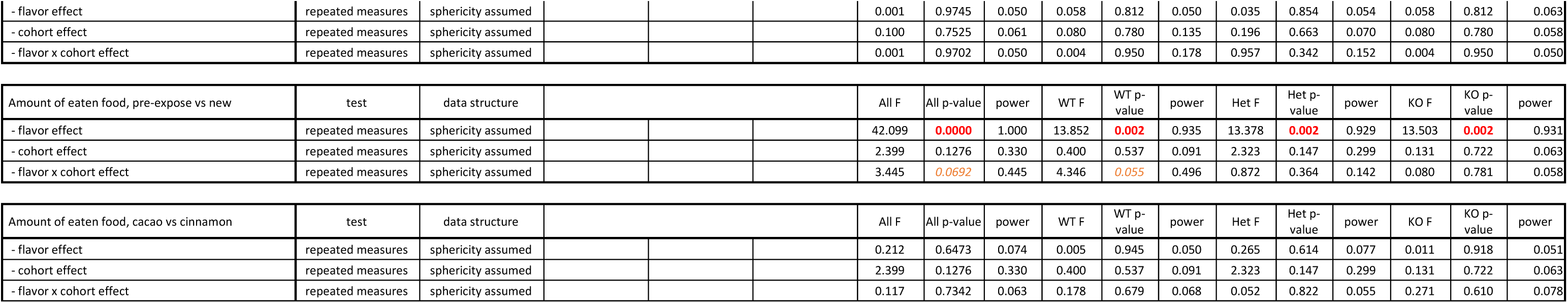
Detailed results and statistical analyses related to social behavior. WT, wild-type mice; Het, heterozygous mice; KO, homozygous knockout mice. C, center chamber; M, mouse chamber; O, object chamber. Group values are reported as means ± s.e.m. Red font indicates significant results (p<0.05), orange font indicates trends (0.1<p<0.05). Individual results and statistical analyses for cohorts 1 and 2 are available in Extended Table 8-1

**Figure 5:**
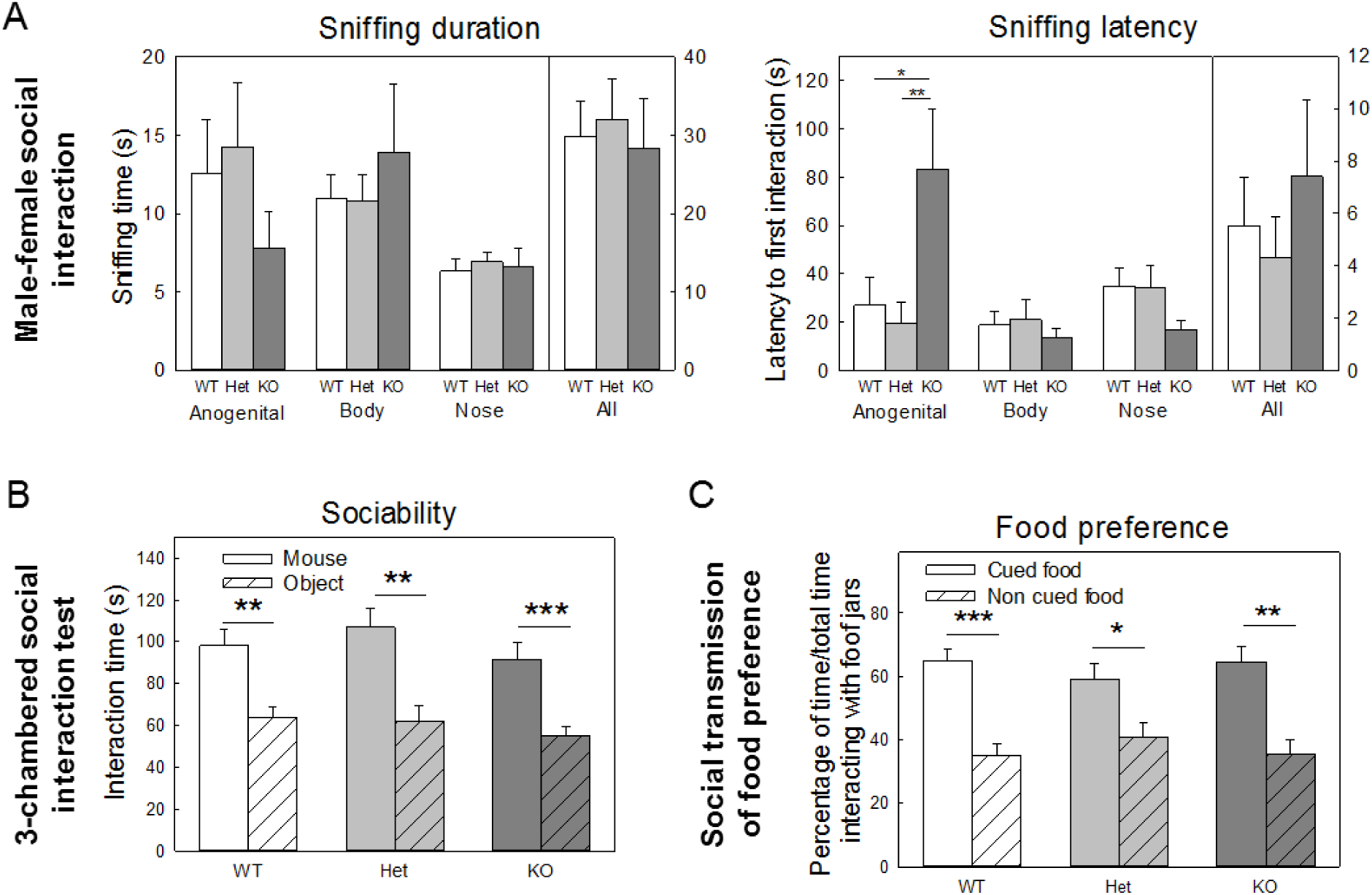
Social behavior of *Shank3^Δ4-22^-*deficient mice. **(A)** Male social interaction in response to the presentation of an unfamiliar conspecific female in estrus and scored by the cumulative sniffing time and latency from the male toward different body regions of the female and the number of ultrasonic vocalizations (USV). No genotype differences were evident in the dyadic male-female social interaction for the overall sniffing time from the male toward the female however a trend toward a decrease of anogenital sniffing as well as a significant increase of the latency to initiate the first anogenital sniffing event was observed in *Shank3^Δ4-22^* homozygous mice. A non-significant decrease of the number of ultrasonic vocalization was also seen in males *Shank3^Δ4-22^* homozygous mice upon exposure to an estrus female. **(B)** Preference for social stimulus in the 3-chambered social interaction test measured by cumulative time interacting with either a mouse or an unanimated object. Normal social preference in *Shank3^Δ4-22^-*deficient mice in the 3-chambered sociability test. All three genotypes demonstrated a significant preference for an unfamiliar mouse over a non-social object. **(C)** Social transmission of food preference measured by the time spent by the test mouse sniffing the demonstrator mouse and the time spent interacting with both cured and non-cued food. All genotypes had a strong preference for the food flavor presented by the demonstrator mouse. Ultrasonic vocalizations and time spent sniffing the demonstrator during the demonstrator interaction phase are displayed on Extended Figure 5-1. WT, wild-type mice; Het, heterozygous mice; KO, homozygous knockout mice. *: p<0.05, **: p<0.1, ***: p<0.001.

Similarly, In the 3-chambered test for social preference, sociability, defined as spending more time interacting with the mouse than with the object, was found in all genotypes. Hence, in all groups, significantly more time was spent in the chamber containing the novel mouse than in the chamber containing the novel object, and more time was spent sniffing the novel mouse than the novel object (Figure 5B). All genotypes showed the normal absence of innate chamber side bias during the 10-minute habituation phase before the start of the sociability test.

Finally, mice were tested in the social transmission of food preference test that combines social behavior, olfactory recognition and memory skills. A modest decrease of the number of sniffing bouts initiated by the observer mouse towards the demonstrator mouse was observed during the observer-demonstrator interaction phase in *Shank3^Δ4-22^* homozygous mice but not in heterozygotes (Extended Figure 5-1 B). All genotypes showed a strong preference for the cued food flavor that was exposed to them through the demonstrator, as compared to the non-cued food flavor, as shown both by significantly more time spent interacting with the jar containing the cued food than the non-cued food (Figure 5C) or by eating significantly more cued food than non-cued food during the choice phase (Table 8). Note that two flavors were randomly used as cued and non-cued food flavor and all genotypes showed an absence of flavor preference. However, the total amount of food (cued and non-cued) eaten by *Shank3^Δ4-22^* homozygous mice was significantly lower than the total amount of food eaten by their wild-type and homozygous littermates.

### Object avoidance in *Shank*^*Δ4-22*^-deficient mice

While testing mice in different set-ups involving object interactions, a strong avoidance toward inanimate objects was observed in *Shank3^Δ4-22^* homozygous mice (Table 9).

**Table 9:**
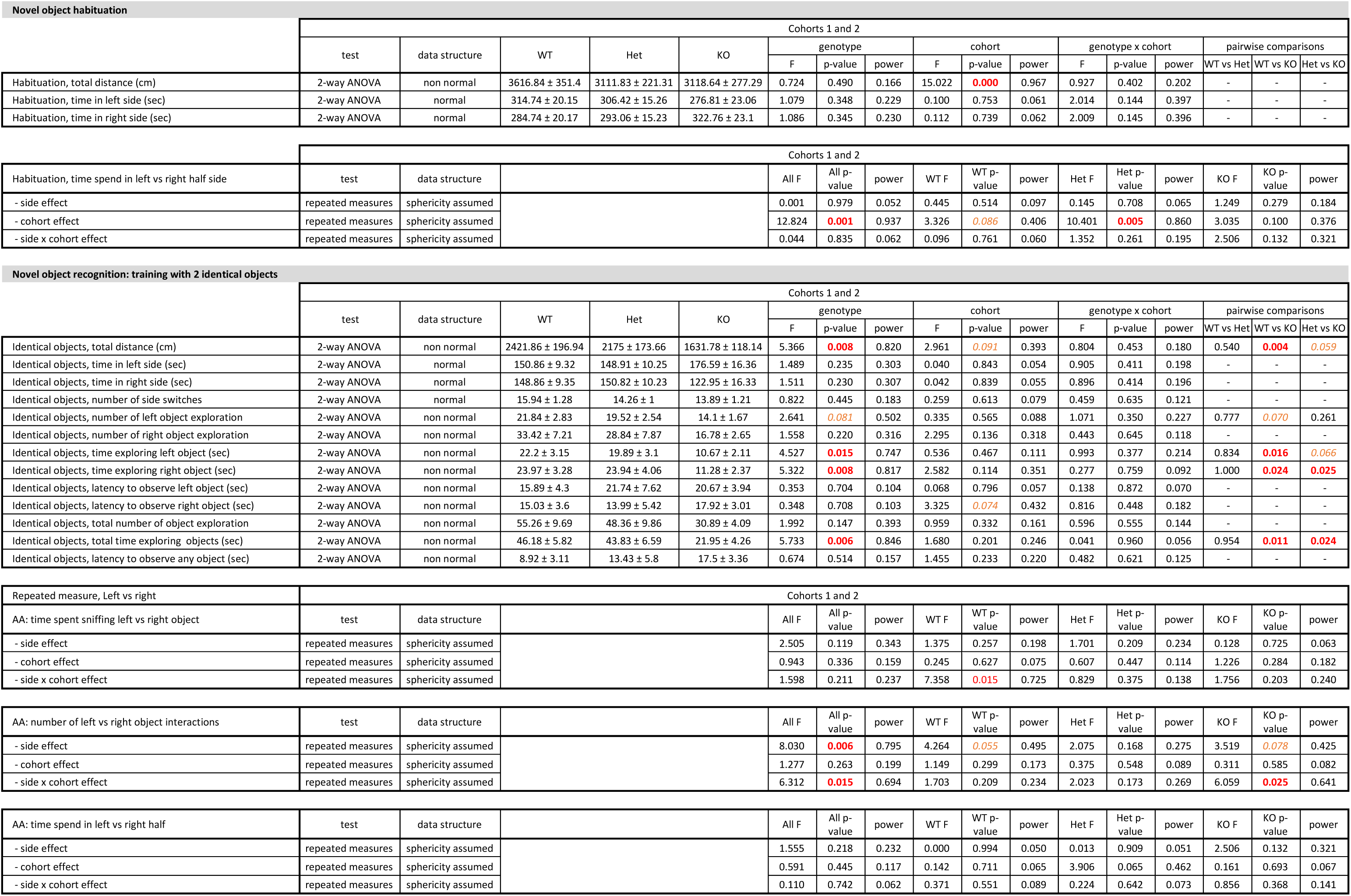
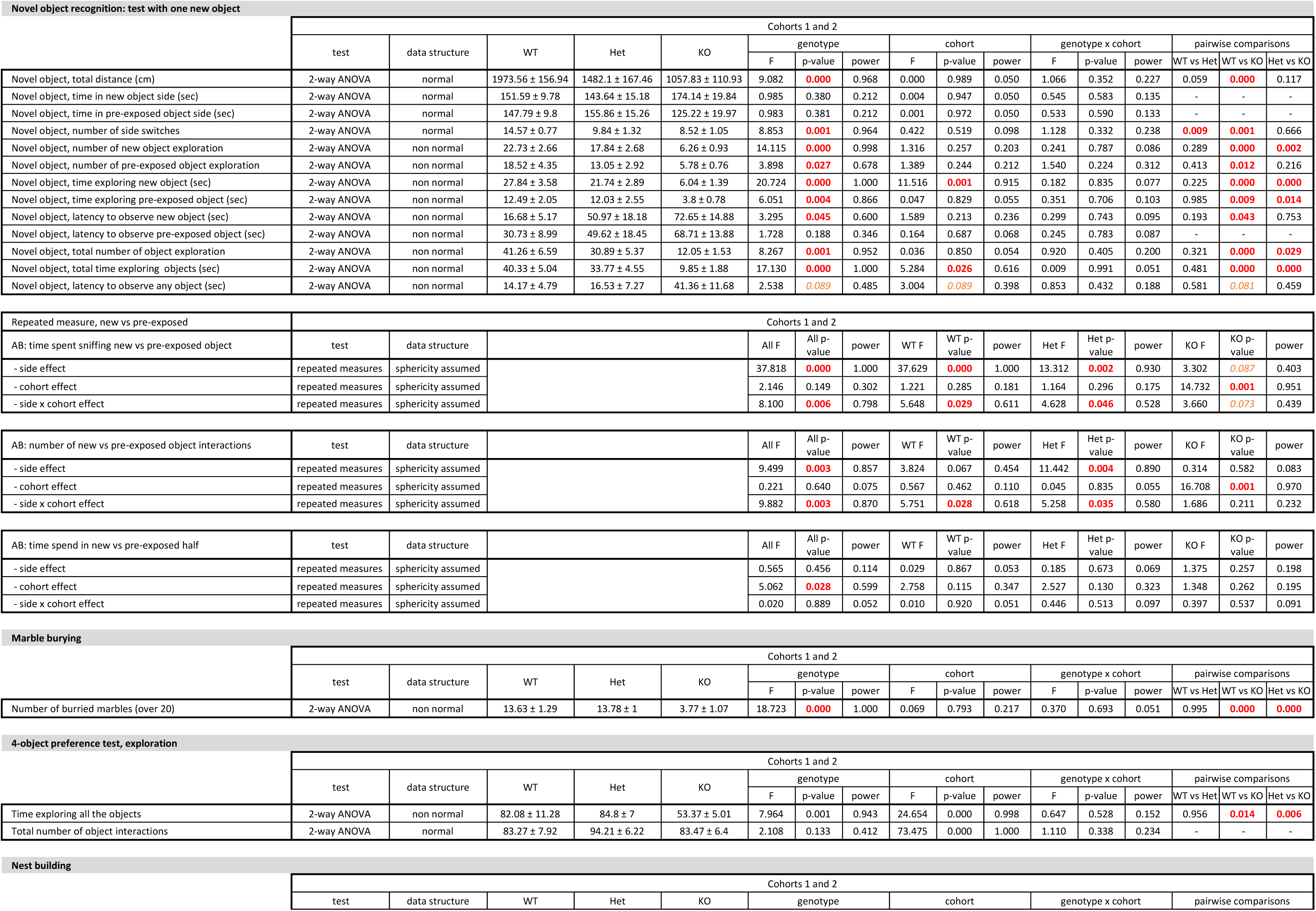
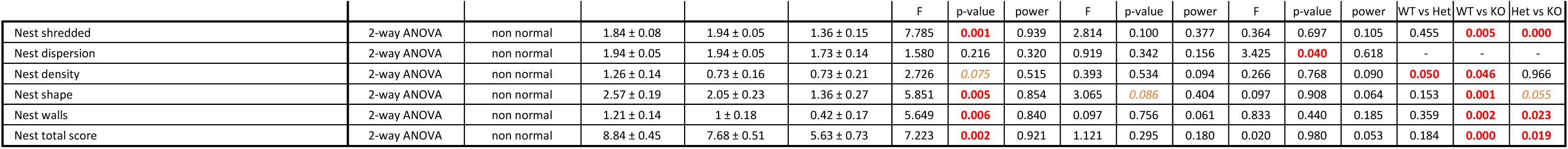
Detailed results and statistical analyses related to the avoidance behavior. WT, wild-type mice; Het, heterozygous mice; KO, homozygous knockout mice. Group values are reported as means ± s.e.m. Red font indicates significant results (p<0.05), orange font indicates trends (0.1<p<0.05). Individual results and statistical analyses for cohorts 1 and 2 are available in Extended Table 9-1

This avoidance behavior was initially observed in the novel object recognition task. This highly validated test for recognition memory is designed to evaluate differences in the exploration time of novel and familiar objects. Mice are expected to spend more time investigating a novel object than a familiar object, and this is what was observed for wild-type and heterozygous mice (Figure 6A left panel). However, in homozygous mice, results were difficult to interpret due to strikingly reduced object interactions (Figure 6A left and middle panels). Homozygous mice spent most of both of the test sessions (the first involving familiarization with identical objects and the second involving interaction with one familiar and one novel object) away from both objects, spending excessive time in the corners of the open-field as shown on heatmaps (Extended Figure 6-1 A) and demonstrating longer latency to explore any of the objects (Figure 6A, right panel).

**Figure 6:**
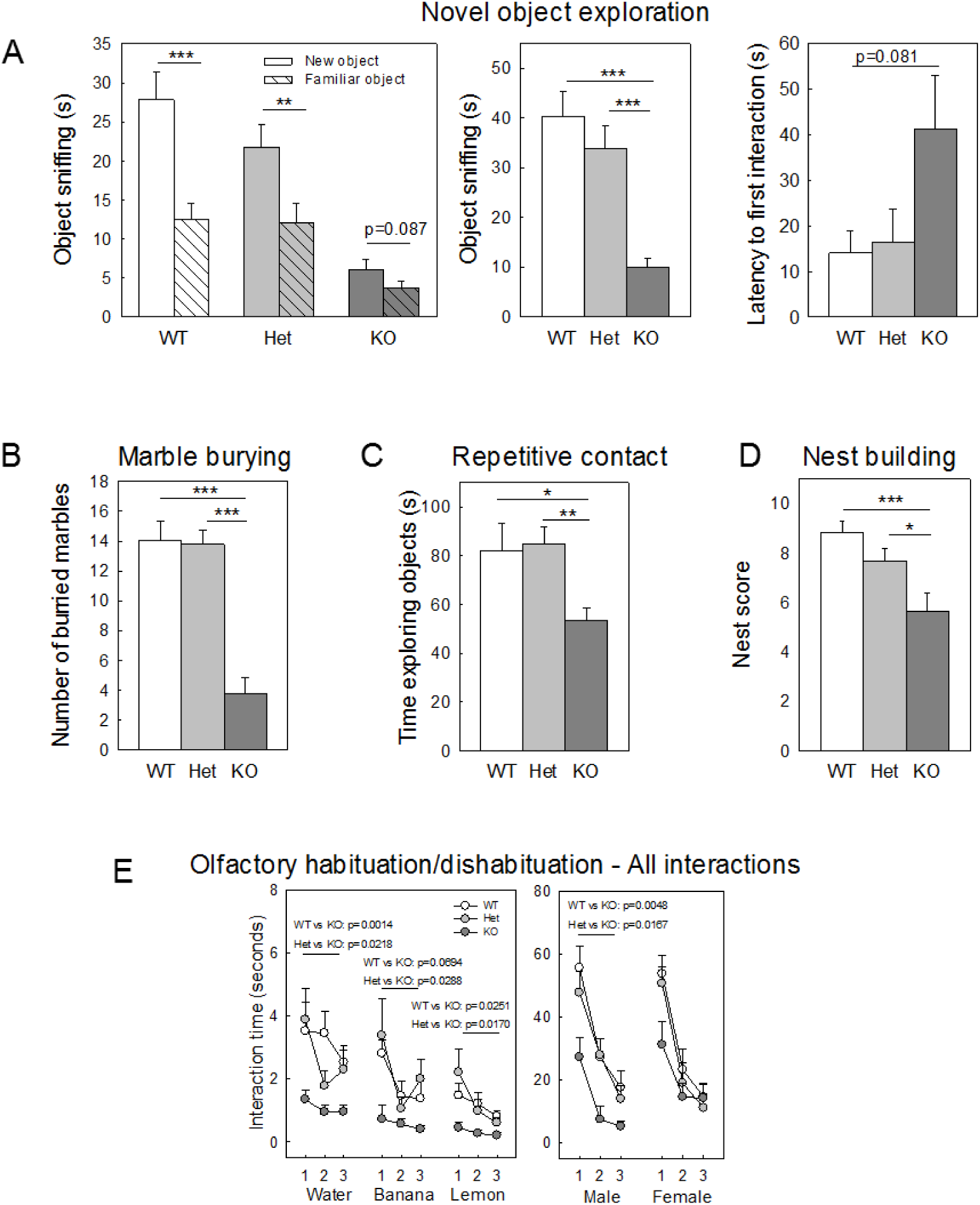
Object avoidance behavior in *Shank3^Δ4-22^-*deficient mice. **(A)** Short term memory measured by the time interaction with familiar and new object in the novel object recognition test. The test consisted of a training with two identical objects followed one hour later by a testing session where one of the object was replaced by a novel object. During the testing session, both wild-type and *Shank3^Δ4-22^* heterozygous mice had a strong preference for the novel object over the familiar object while *Shank3^Δ4-22^* homozygous mice failed to display a preference. However, this failure was due to an avoidance of both objects as shown by the strong decrease of object interaction and the increase of latency to explore any of the object for the first time in *Shank3^Δ4-22^* homozygous animals rather than to a real lack of object preference. Representative heatmaps for the three genotypes are available on Extended Figure 6-1 A. **(B)** Repetitive behavior and object avoidance measured in the marble burying test by the number of marble buried during a 30-minute session. *Shank3^Δ4-22^* homozygous mice displayed a strongly impaired burying behavior leaving most of the marbles undisturbed. Representative pictures and individual data are displayed on Extended Figure 6-1 B. **(C)** Time spend exploring objects in the repetitive novel object contact task. *Shank3^Δ4-22^* homozygous mice spent significantly less time interacting with the objects than their wild-type and heterozygous littermates. **(D)** Nest building scores. *Shank3^Δ4-22^* homozygous mice are building less elaborate nests and use less nesting material than their wild-type and heterozygous littermates. Representative pictures of the nests and individual data are displayed on Extended Figure 6-1 C. **(E)** Time interacting with the scent applicator (touching, biting, climbing) during the olfactory habituation/dishabituation time. *Shank3^Δ4-22^* homozygous mice are avoiding interaction with the scent applicator for all non-social scents and for social male scent but have interaction level similar to wild-type and heterozygous animals when presented with a female scent. WT, wild-type mice; Het, heterozygous mice; KO, homozygous knockout mice. *: p<0.05, **: p<0.1, ***: p<0.001.

Object avoidance was further confirmed in multiple independent tests, including the marble burying, a test used to assess stereotypic behavior and/or anxiety. In this paradigm, 20 marbles were spread across the cage floor in a 4 × 5 pattern, leaving little space for the mice to move around the marbles. While both wild-type and *Shank3^Δ4-22^* heterozygous mice quickly buried most of the marbles as is typical, *Shank3^Δ4-22^* homozygous mice left the marbles almost completely undisturbed for the whole 15-minute duration of the test (Figure 6B and Extended Figure 6-1 B).

Consistent with these result, a significant decrease of the time spent exploring objects in the 4-object exploration test was observed in the *Shank3^Δ4-22^* homozygous mice as compared to their littermate (Figure 6C).

During assessment of nest building, nests build by *Shank3^Δ4-22^* homozygous mice were significantly less elaborate than nests built by wild-type or heterozygous mice, with some homozygotes leaving the nesting material completely untouched (Figure 6D, Extended Figure 6-1 C). Note that, in an attempt to reduce stress and improve breeding rates, dams used to produce the cohorts described here were provided with plastic huts in their home cage. Interestingly, while most of the wild-type dams (seven out of ten) chose to build their nest inside the huts, only a single *Shank3^Δ4-22^* heterozygous dam out of twenty used the hut to establish their nests (wild-type versus heterozygotes: t28=-5.085, p<0.001). Additionally, three of the *Shank3^Δ4-22^* heterozygous dams did not build a nest until after the birth.

Object avoidance might also explain the reduction of the total time of direct interactions (grabbing, touching, biting or climbing) with the applicator used to present the different scents during the olfactory habituation and dishabituation test in *Shank3^Δ4-22^* homozygous mice, compared to their wild-type and heterozygous littermates (Figure 6E).

### Hyper-reactivity and escape behaviors in *Shank3^Δ4-22^*-deficient mice

Unusual hyper-reactivity was observed in *Shank3^Δ4-22^* homozygous mice during handling and confirmed in several behavioral tests (Table 10). This hyper-reactivity was characterized by a higher score in the touch escape test (Figure 7A, left panel), a lower score (reflecting a higher tendency to struggle in response to sequential handling) in the positional passivity (Figure 7A, middle panel) and a shorter latency to move from the beam during the catalepsy test (Figure 7A, right panel). As in newborn mice, a shorter latency to turn was seen for *Shank3^Δ4-22^* homozygous mice in the negative geotaxis test (Figure 7B, left panel). Similarly, in the beam walking test, the latency to start crossing on the smallest beam was shorter in *Shank3^Δ4-22^* homozygous mice (Figure 7B right panel) but often led to a premature fall (Figure 3D).

**Table 10:** Detailed results and statistical analyses related to the hyper-reactivity and escape behavior. WT, wild-type mice; Het, heterozygous mice; KO, homozygous knockout mice. Group values are reported as means ± s.e.m. Red font indicates significant results (p<0.05), orange font indicates trends (0.1<p<0.05). Individual results and statistical analyses for cohorts 1 and 2 are available in Extended Table 10-1

| Reflexes and reactions to simple stimuli |  |  |  |  |  |  |  |  |  |  |  |  |  |  |  |  |  |
| --- | --- | --- | --- | --- | --- | --- | --- | --- | --- | --- | --- | --- | --- | --- | --- | --- | --- |
|  | Cohorts 1 and 2 |  |  |  |  |  |  |  |  |  |  |  |  |  |  |  |  |
|  | test | data structure | WT | Het | KO | genotype |  |  | cohort |  |  | genotype x cohort |  |  | pairwise comparisons |  |  |
|  |  |  |  |  |  | F | p-value | power | F | p-value | power | F | p-value | power | WT vs Het | WT vs KO | Het vs KO |
| Touch escape | 2-way ANOVA | non normal | 1.26 ± 0.1 | 1.15 ± 0.11 | 2 ± 0.15 | 12.962 | **0.000** | 0.996 | 0.046 | 0.831 | 0.055 | 0.862 | 0.428 | 0.190 | 0.648 | **0.000** | **0.000** |
| Positional passivity (sum) | 2-way ANOVA | non normal | 2.15 ± 0.25 | 1.84 ± 0.23 | 2.84 ± 0.2 | 11.737 | **0.000** | 0.992 | 106.722 | **0.000** | 1.000 | 1.993 | 0.147 | 0.393 | **0.034** | **0.011** | **0.000** |
| Positional passivity (score) | 2-way ANOVA | non normal | 1.78 ± 0.24 | 2.21 ± 0.21 | 0.94 ± 0.2 | 14.029 | **0.000** | 0.998 | 44.935 | **0.000** | 1.000 | 3.871 | **0.027** | 0.675 | **0.034** | **0.004** | **0.000** |
| Catalepsy (4 trials) | 2-way ANOVA | non normal | 2.98 ± 0.57 | 2.75 ± 0.54 | 0.56 ± 0.25 | 7.578 | **0.001** | 0.933 | 4.681 | **0.035** | 0.565 | 1.116 | 0.336 | 0.236 | 0.836 | **0.001** | **0.002** |
| Trunk curl | 2-way ANOVA | non normal | 1 ± 0 | 0.89 ± 0.07 | 1 ± 0 | 2.547 | *0.088* | 0.487 | 2.537 | 0.117 | 0.346 | 2.547 | *0.088* | 0.487 | *0.057* | 1.000 | *0.056* |
| Negative geotaxis, latency to turn | 2-way ANOVA | non normal | 6.73 ± 1.06 | 8.21 ± 1.9 | 3.22 ± 0.58 | 4.201 | **0.020** | 0.713 | 2.978 | 0.090 | 0.395 | 0.707 | 0.498 | 0.163 | 0.499 | **0.042** | **0.008** |

| Beam walking |  |  |  |  |  |  |  |  |  |  |  |  |  |  |  |  |  |
| --- | --- | --- | --- | --- | --- | --- | --- | --- | --- | --- | --- | --- | --- | --- | --- | --- | --- |
|  | Cohorts 1 and 2 |  |  |  |  |  |  |  |  |  |  |  |  |  |  |  |  |
|  | test | data structure | WT | Het | KO | genotype |  |  | cohort |  |  | genotype x cohort |  |  | pairwise comparisons |  |  |
|  |  |  |  |  |  | F | p-value | power | F | p-value | power | F | p-value | power | WT vs Het | WT vs KO | Het vs KO |
| Latency to start crossing the large beam | 2-way ANOVA | non normal | 8.34 ± 2.79 | 10.8 ± 4.52 | 3.96 ± 1.75 | 1.163 | 0.321 | 0.244 | 0.251 | 0.618 | 0.078 | 1.300 | 0.281 | 0.269 | - | - | - |
| Latency to start crossing the medium beam | 2-way ANOVA | non normal | 15.9 ± 4.8 | 7.17 ± 4.16 | 7.65 ± 6.13 | 0.811 | 0.450 | 0.181 | 3.349 | *0.073* | 0.435 | 0.121 | 0.887 | 0.068 | - | - | - |
| Latency to start crossing the small beam | 2-way ANOVA | non normal | 54.71 ± 5.89 | 46.15 ± 7.59 | 33.72 ± 7.84 | 2.204 | 0.121 | 0.430 | 0.235 | 0.630 | 0.076 | 0.437 | 0.648 | 0.117 | - | - | - |

| Escape behavior |  |  |  |  |  |  |  |  |  |  |  |  |  |  |  |  |  |
| --- | --- | --- | --- | --- | --- | --- | --- | --- | --- | --- | --- | --- | --- | --- | --- | --- | --- |
|  | Cohorts 1 and 2 |  |  |  |  |  |  |  |  |  |  |  |  |  |  |  |  |
|  | test | data structure | WT | Het | KO | genotype |  |  | cohort |  |  | genotype x cohort |  |  | pairwise comparisons |  |  |
|  |  |  |  |  |  | F | p-value | power | F | p-value | power | F | p-value | power | WT vs Het | WT vs KO | Het vs KO |
| Buried food, number of escape attempts | 2-way ANOVA | non normal | 0.63 ± 0.35 | 0.68 ± 0.32 | 0.42 ± 0.31 | 0.666 | 0.519 | 0.155 | 1.738 | 0.194 | 0.253 | 0.220 | 0.804 | 0.082 |  |  |  |
| Buried food, percentage of mice escaping | 2-way ANOVA | non normal | 21.05 ± 9.6 | 26.31 ± 10.37 | 15.78 ± 8.59 | 0.760 | 0.473 | 0.172 | 3.639 | *0.062* | 0.464 | 0.159 | 0.853 | 0.073 |  |  |  |
| 4-object exploration, number of escape attempts | 2-way ANOVA | non normal | 0.41 ± 0.21 | 2.05 ± 0.73 | 3.88 ± 1.21 | 5.323 | **0.008** | 0.815 | 5.320 | **0.025** | 0.618 | 3.316 | **0.045** | 0.601 | 0.187 | **0.002** | **0.050** |
| 4-object exploration, percentage of mice escaping | 2-way ANOVA | non normal | 23.52 ± 10.6 | 36.84 ± 11.36 | 50 ± 12.12 | 1.502 | 0.233 | 0.305 | 4.351 | **0.042** | 0.534 | 2.575 | *0.087* | 0.490 |  |  |  |
| Marble burying, number of escape attempts | 2-way ANOVA | non normal | 4.32 ± 1.28 | 10.63 ± 1.98 | 16.05 ± 2.38 | 8.063 | **0.001** | 0.946 | 6.649 | **0.013** | 0.715 | 1.239 | 0.299 | 0.257 | **0.034** | **0.000** | *0.055* |
| Marble burying, percentage of mice escaping | 2-way ANOVA | non normal | 47.36 ± 11.76 | 94.73 ± 5.26 | 100 ± 0 | 12.009 | **0.000** | 0.993 | 7.713 | **0.008** | 0.777 | 4.474 | **0.017** | 0.740 | **0.000** | **0.000** | 0.598 |

**Figure 7:**
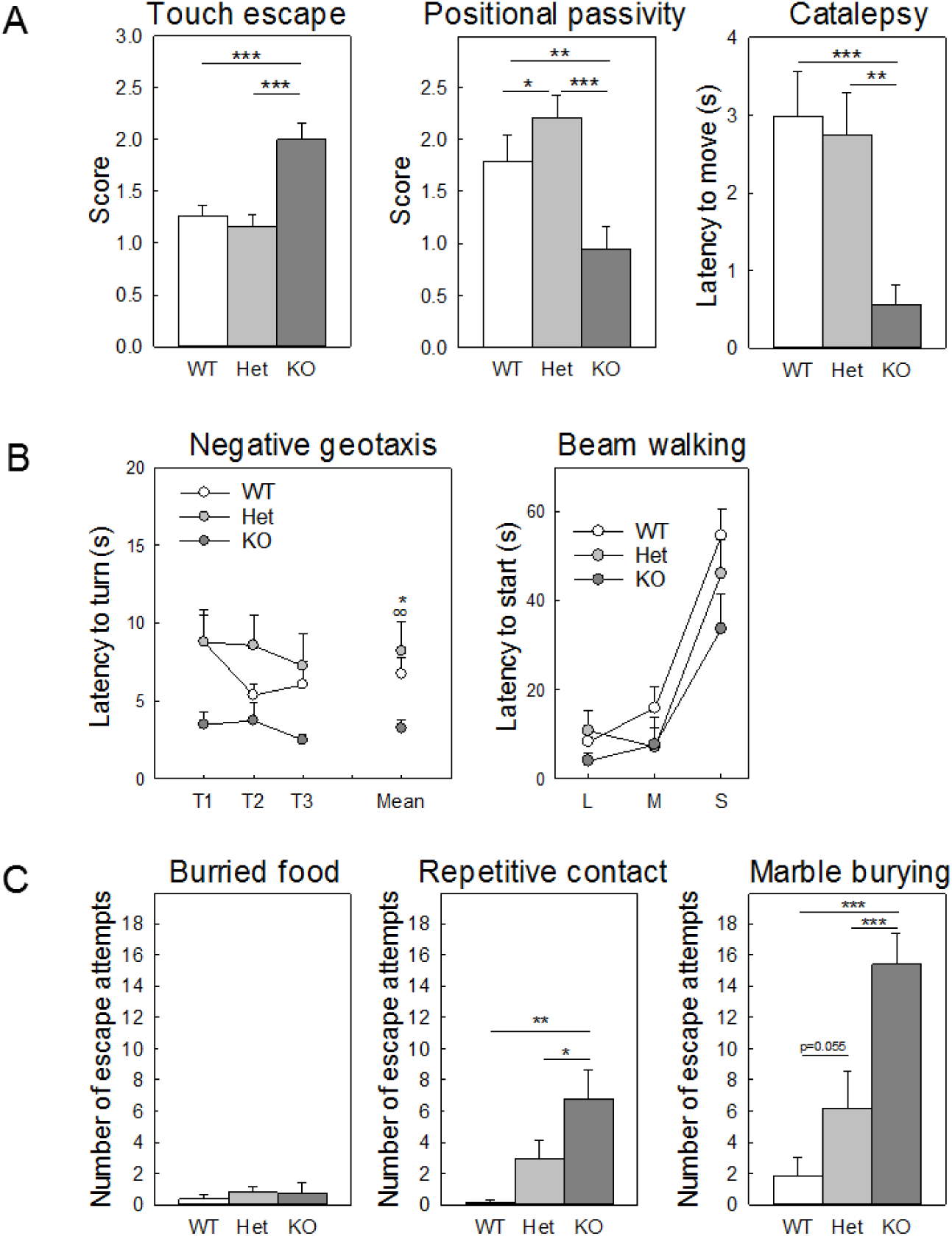
Hyper-reactivity and escape behavior in *Shank3^Δ4-22^-*deficient mice. **(A)** Hyper-reactivity measured by animal response in touch escape, positional passivity and catalepsy. *Shank3^Δ4-^ 22* homozygous mice have hyper-reactive responses as shown by a higher score in the touch escape indicating an escape response to lighter strokes, a lower score in score in positional passivity indicating that they struggle more when restrained and a lower latency to get down a rod in the catalepsy test. **(B)** Impulsivity in the negative geotaxis and beam waling tests. The latency to start turning in the negative geotaxis test and to start crossing in the beam walking test are significantly lower in *Shank3^Δ4-22^* homozygous mice compared to their wild-type and heterozygous littermates and often associated with higher failure rates (not shown) thus demonstrating impulsive behavior. **(C)** Escape behavior measured in different tests with increased inanimate object exposure. No escape attempts were observed for any genotype during the habituation phase of the buried food test (empty home cage with clean bedding). Object exposure induced a significant escape behavior in *Shank3^Δ4-22^* homozygous mice with a number of attempts increasing with the number of objects in the cage (same home cage, four objects in the repetitive novel object contact task, twenty objects in the marble burying test). Very little escape attempts were observed in wild-type mice while an intermediate phenotype was observed in heterozygous mice. WT, wild-type mice; Het, heterozygous mice; KO, homozygous knockout mice. *: p<0.05, **: p<0.1, ***: p<0.001.

Escape attempts were observed in several tests and high-wall enclosures had to be built around testing cages to prevent successful attempts. Escape behaviors were scored in three different home cage tests. During the habituation portion of the buried food test (where no objects were visible at the surface of the cage bedding), no escape behavior nor genotype differences were observed (Figure 7C, left panel). However, when the mice were tested in the same cages during the 4-object interaction test both the number of escape attempts and the percentage of mice engaged in this behavior increased and significant genotype differences were observed (Figure 7C middle panel). This behavior was even more marked in the marble burying test (Figure 7C, right panel), during which 94% of heterozygous mice and 100% of homozygous mice tried to escape. This indicated that the escape behavior is elicited by the presence of unfamiliar objects in the testing cage.

### Repetitive behaviors, stereotypies and inflexibly in *Shank3^Δ4-22^*-deficient mice

Repetitive and restricted behaviors are one of the core features of ASD. Therefore, during all of the behavioral tests, mice were also carefully monitored for stereotypies, as well as perseverative and repetitive behaviors. Detailed results are reported in Table 11.

**Table 11:**
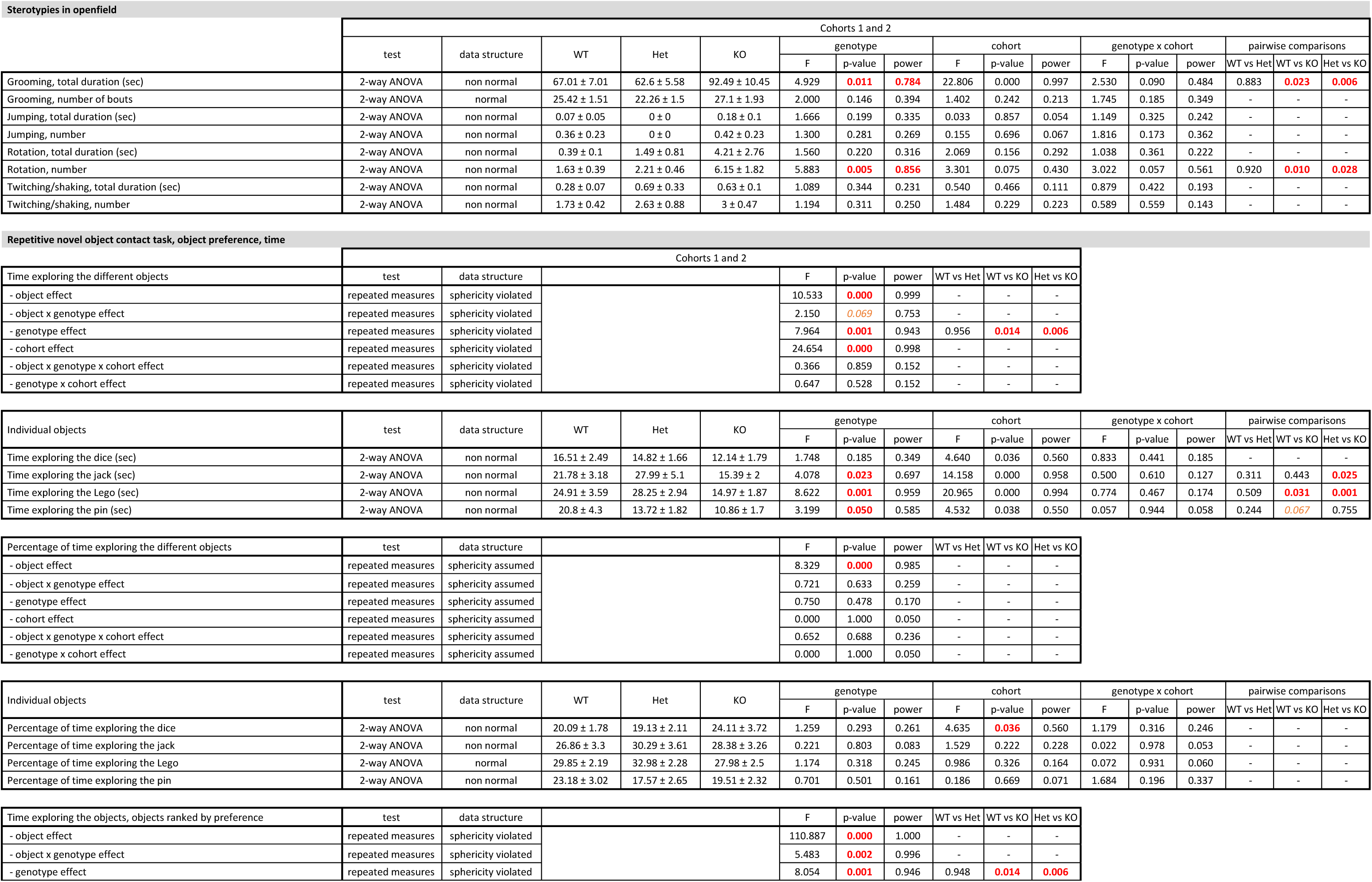
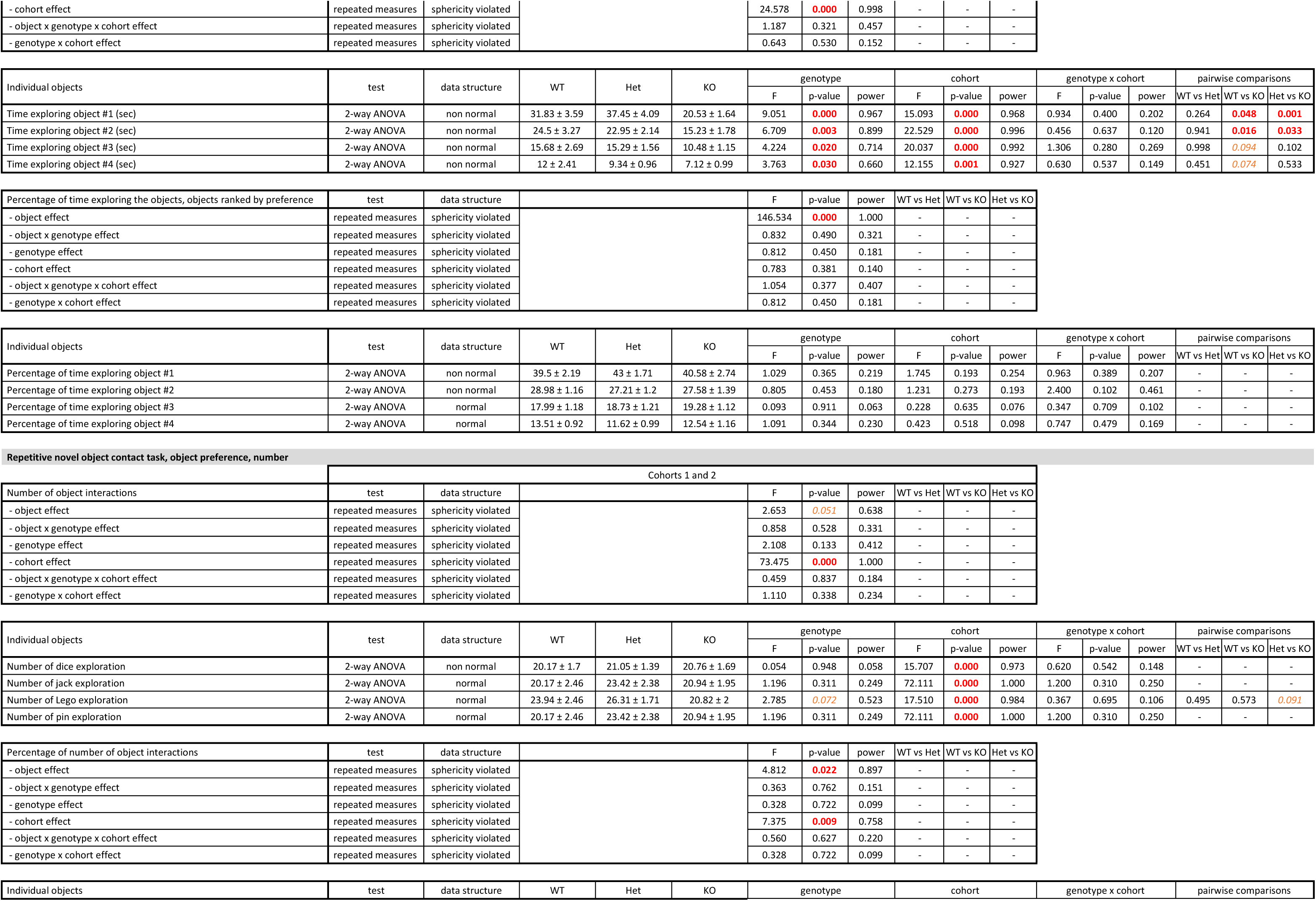
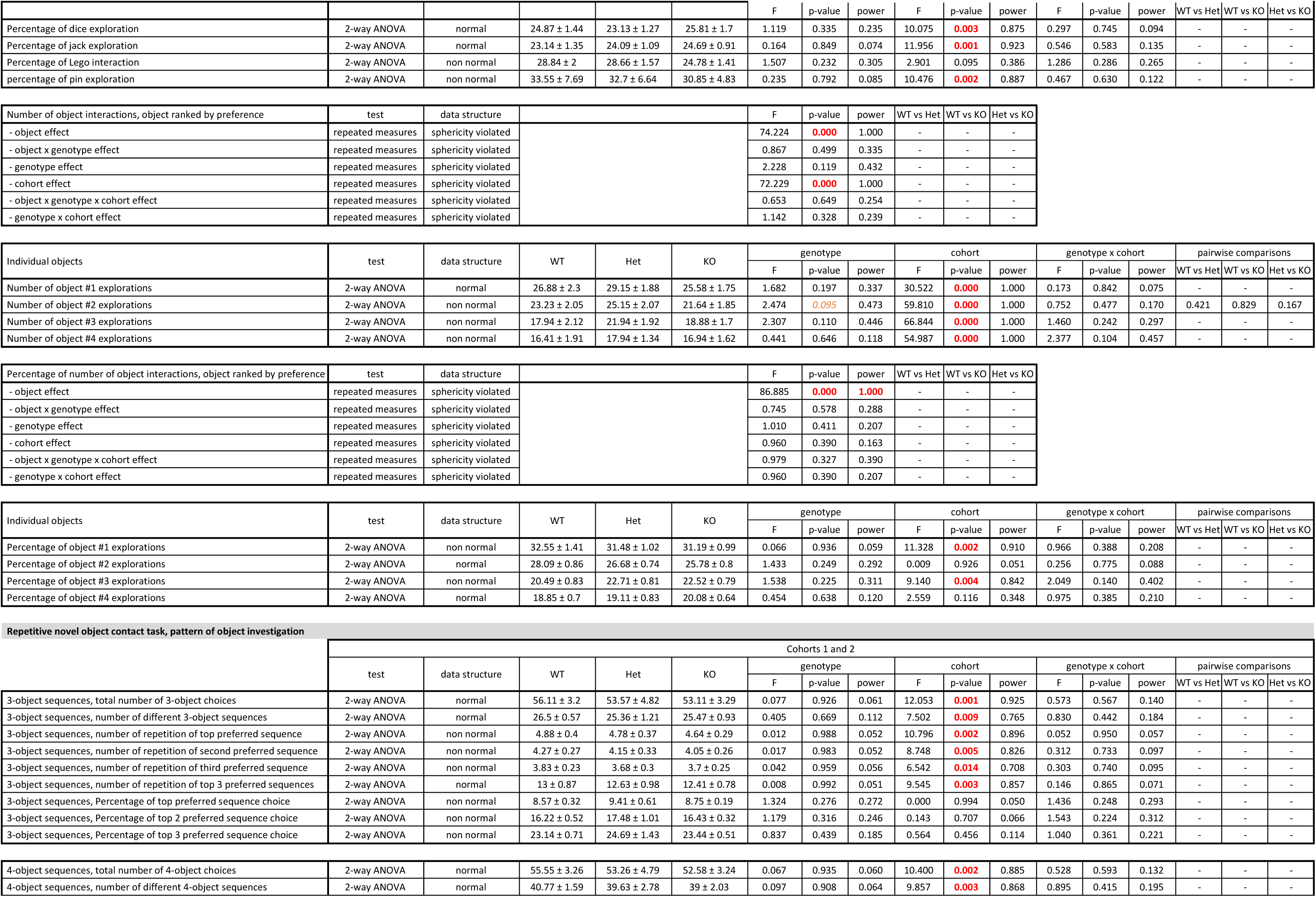
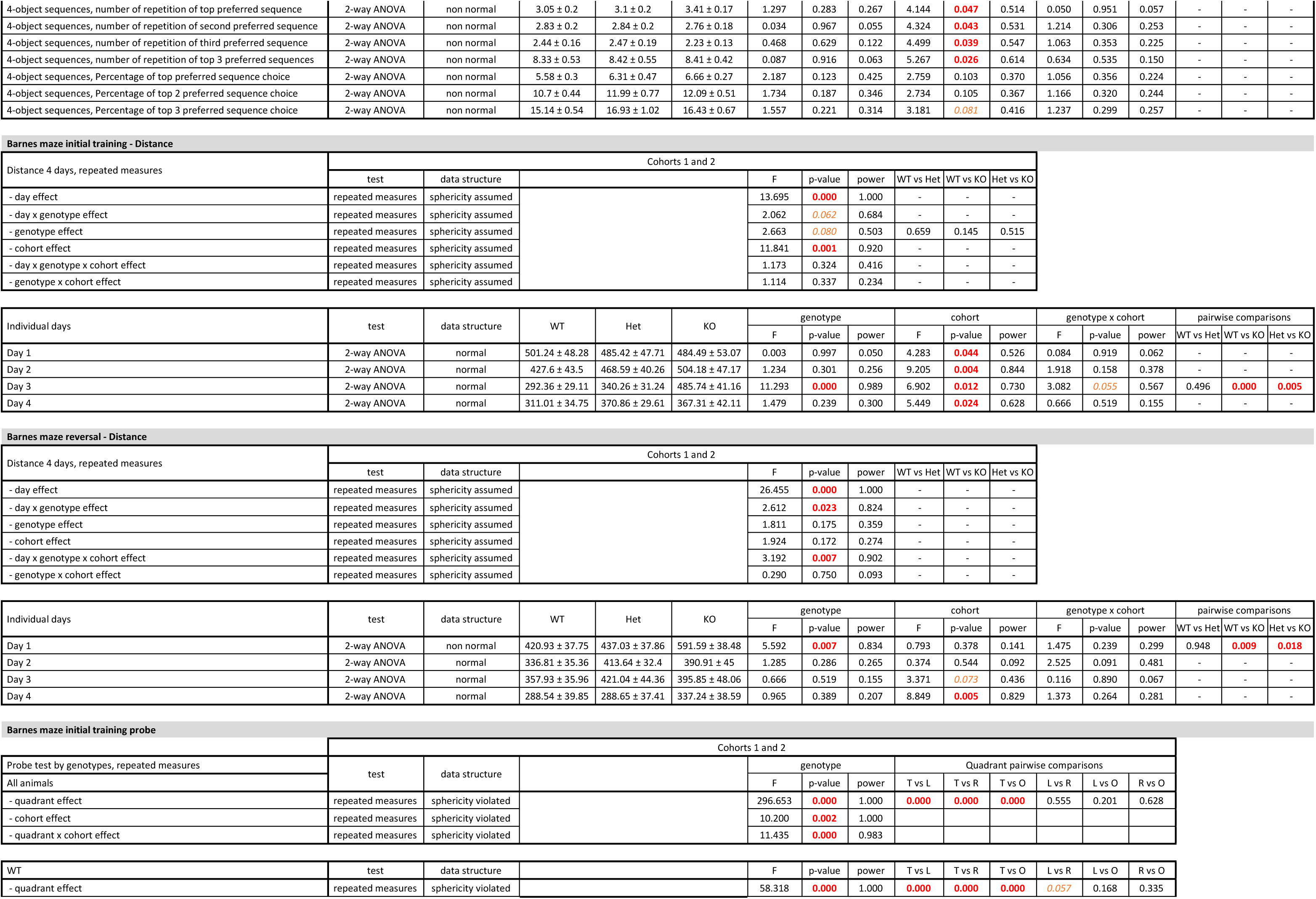
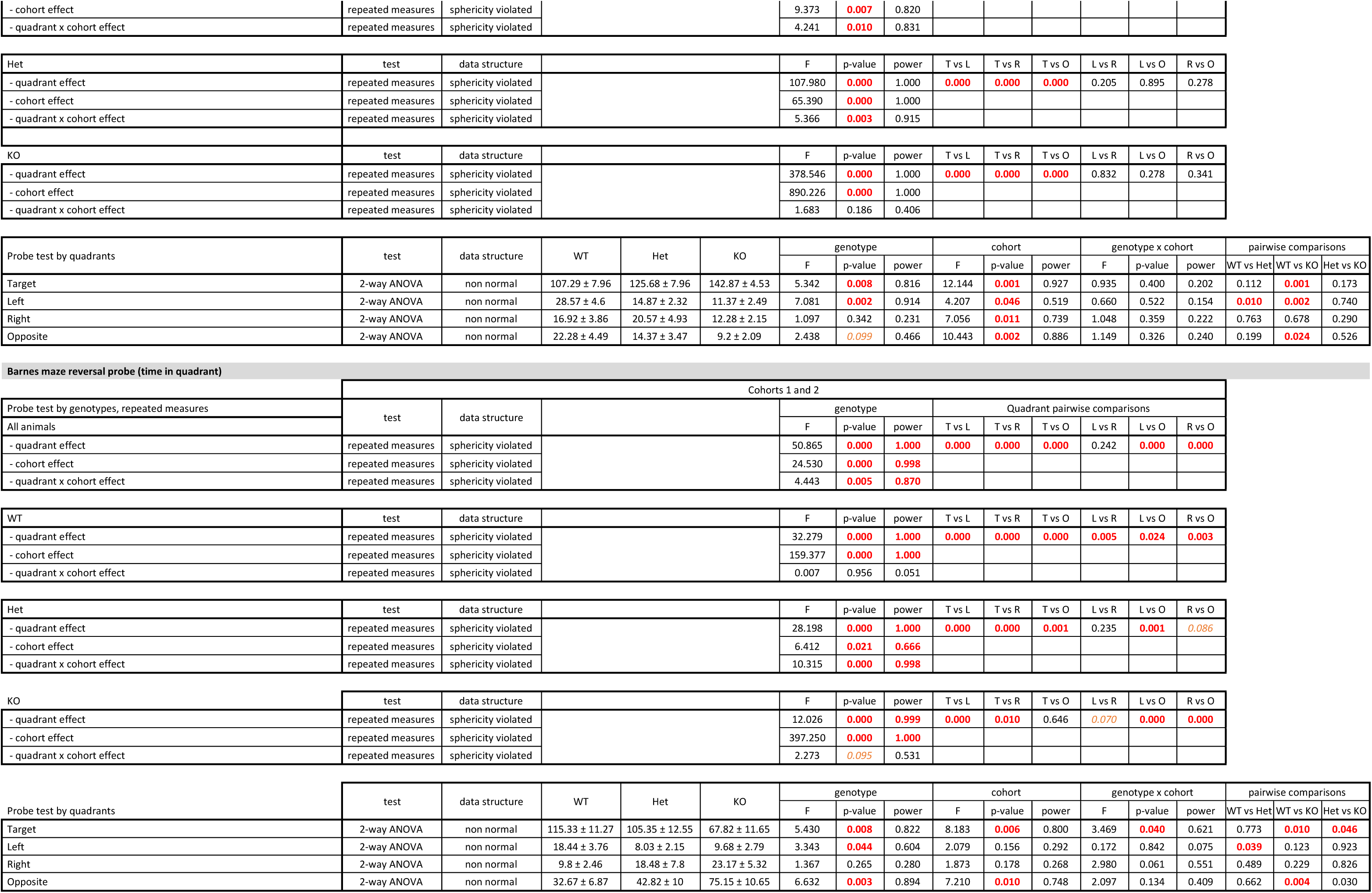
Detailed results and statistical analyses related to stereotypies, repetitive behavior, perseveration and cognitive flexibility. WT, wild-type mice; Het, heterozygous mice; KO, homozygous knockout mice. Group values are reported as means ± s.e.m. Red font indicates significant results (p<0.05), orange font indicates trends (0.1<p<0.05). Individual results and statistical analyses for cohorts 1 and 2 are available in Extended Table 11-1

While no genotype difference was observed in the number of spontaneous grooming bouts observed during the ten first minutes of the open-field test, *Shank3^Δ4-22^* homozygous mice engaged in longer episodes of self-grooming, as shown by a significant increase in the cumulative time spent grooming all body regions when compared to their wild-type and heterozygous littermates. However, skin lesions were frequently observed in older mice (over 8-month-old) of all three genotypes without obvious genotype effect. Significantly more rotations were also observed in *Shank3^Δ4-22^* homozygous animals as well as a trend towards an increase of head twitching/shaking in both *Shank3^Δ4-22^* heterozygous and homozygous mice, as compared to their wild-type littermates (Figure 8A).

**Figure 8:**
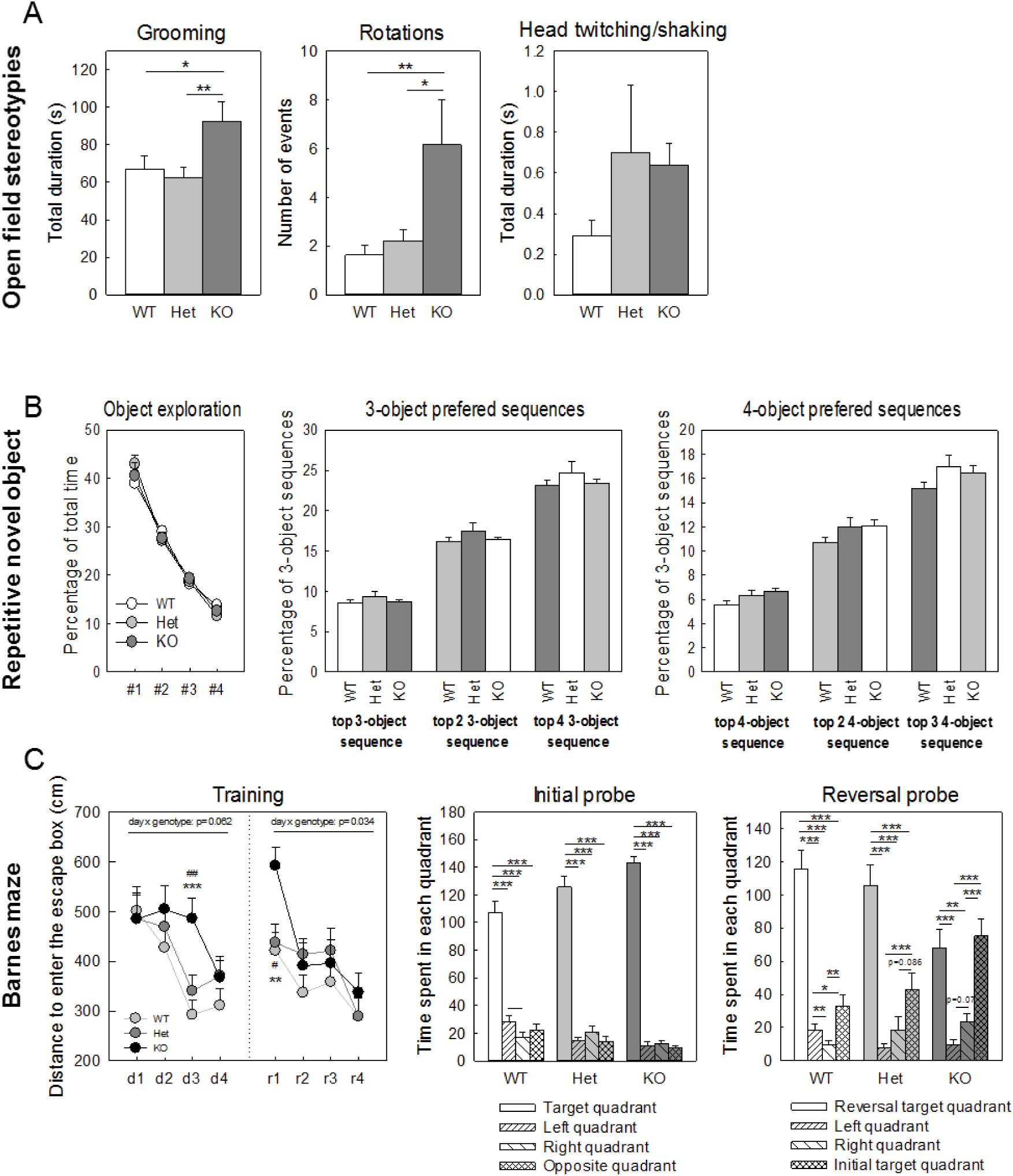
Repetitive behavior, stereotypies and cognitive flexibility in *Shank3^Δ4-22^-*deficient mice. **(A)** Repetitive behaviors in the open field test. *Shank3^Δ4-22^* homozygous mice engaged in significantly more self-grooming and rotations relative to the other genotypes. A trend toward an increase amount of head stereotypies was also observed. **(B)** Object preference and pattern of exploration in the repetitive novel object contact task. For each mouse, the time spent interacting with each object was measured and the objects were then ranked from the most (1) to less (4) preferred (left panel). No genotype differences were observed for the proportions of visits to each object. The pattern of object exploration was analyzed by recording specific sequential pattern of visits to three or four specific toys to identify the total number of 3-object or 4-object sequence investigations, the number of unique sequences and the percentage of choices of the top, top two or top three preferred sequence. All groups had identical percentage of their preferred 3-object or 4-object sequences choices over the total number of sequence choices. **(C)** Cognitive flexibility measured by reversal learning in the Barnes maze. During initial learning (d1 to d4, each day point represents the mean of travelled distance for four independent trials), improvement shown by reduction of the travel distance was faster in *Shank3^Δ4-22^* wild-type and heterozygous mice than in homozygous animals however by day 4 the three groups were not different anymore and all of them had a strong preference for the escape hole quadrant during the initial probe test. During the reversal training (r1 to r4, each day point represents the mean of travel distance for four independent trials) *Shank3^Δ4-22^* homozygous mice initially traveled for longer distances but were still able to learn the new position and performed as well as their littermates on reversal days 2, 3 and 4. However, the reversal probe test at the end of the reversal training showed that while wild-type and heterozygous animals had a significant preference for the new target quadrant, the homozygous mice had a similar preference for the quadrants containing the initial and the reversal escape holes. WT, wild-type mice; Het, heterozygous mice; KO, homozygous knockout mice. *: WT vs KO; #: Het vs KO. *: p<0.05, **: p<0.1, ***: p<0.001.

Object preferences, exploration patterns and frequency of repetitive contacts with novel objects were evaluated in the repetitive novel object contact task. Although the cumulative time spend interacting with the objects was decreased in *Shank3^Δ4-22^* homozygous mice (Figure 6C), this test failed to display genotype difference in either the total number of interactions, the preference for any specific objects or the preference for any specific preferred sequence of 3-object or 4-object explorations (Figure 8B)

Individuals with ASD can maintain rigid habits and frequently show strong insistence on sameness and upset by changes in routine. To examine this domain, *Shank3^Δ4-22^* mice were trained for four days in the Barnes maze, a test of spatial learning and memory, until all the mice were able to quickly locate an escape box hidden under one of the target locations, then the location of the escape box was moved and mice were tested for reversal learning for four additional days. During the initial learning, all the genotypes were able to find the escape hatch equally well, although *Shank3^Δ4-22^* homozygous mice took one day longer to reach criteria (Figure 8C, left panel). All genotypes preferred the correct quadrant in the first probe test ran immediately after the initial training (Figure 8C, middle panel). When the escape hatch was moved to the opposite side of the maze, both *Shank3^Δ4-22^* wild-type and heterozygotes immediately learned the new position, while a one-day delay was, once again, observed for the *Shank3^Δ4-22^* homozygous mice. Genotypes differed markedly in the second probe test, however; while wild-type mice spent most time in the new target quadrant, *Shank3*^Δ4–^ 22 heterozygous mice split their time 75/25% between new and old targets, whereas Shank3^Δ4–22^ homozygous animals spent equal time in both targets (Figure 8C, right panel). This impaired reversal learning implies that *Shank3* deficiency increases susceptibility to proactive interference where learning of a previous rule interferes with the new rule.

### Learning and memory in *Shank3^Δ4-22^*-deficient mice

In addition to the Barnes Maze, animals were tested in two additional learning and memory tests, specifically, the Y-maze spontaneous alternation test and the fear conditioning test. Detailed results are reported in Table 12.

**Table 12:** Detailed results and statistical analyses related to learning and memory. WT, wild-type mice; Het, heterozygous mice; KO, homozygous knockout mice. Group values are reported as means ± s.e.m. Red font indicates significant results (p<0.05), orange font indicates trends (0.1<p<0.05). Individual results and statistical analyses for cohorts 1 and 2 are available in Extended Table 12-1

| Y-maze, spontaneous alternation behavior |  |  | Cohorts 1 and 2 |  |  |  |  |  |  |  |  |  |  |  |  |  |  |
| --- | --- | --- | --- | --- | --- | --- | --- | --- | --- | --- | --- | --- | --- | --- | --- | --- | --- |
|  | test | data structure | WT | Het | KO | genotype |  |  | cohort |  |  | genotype x cohort |  |  | pairwise comparisons |  |  |
|  |  |  |  |  |  | F | p-value | power | F | p-value | power | F | p-value | power | WT vs Het | WT vs KO | Het vs KO |
| Percentage of arm 1 choices | 2-way ANOVA | normal | 32.34 ± 0.87 | 34.24 ± 0.92 | 32.77 ± 1.17 | 0.844 | 0.436 | 0.187 | 0.035 | 0.852 | 0.054 | 0.412 | 0.664 | 0.113 | - | - | - |
| Percentage of arm 2 choices | 2-way ANOVA | normal | 35.17 ± 1.17 | 32.74 ± 1.1 | 35.18 ± 1.36 | 1.548 | 0.223 | 0.314 | 9.976 | 0.003 | 0.873 | 0.119 | 0.888 | 0.067 | - | - | - |
| Percentage of arm 3 choices | 2-way ANOVA | normal | 32.19 ± 1.46 | 32.98 ± 1.09 | 32.04 ± 1.02 | 0.285 | 0.753 | 0.093 | 10.366 | 0.002 | 0.885 | 0.520 | 0.598 | 0.131 | - | - | - |

| Arm preference, t-test comparison to chance level | test | data structure |  | All t | All p-value | power | WT t | WT p-value | power | Het t | Het p-value | power | KO t | KO p-value | power |
| --- | --- | --- | --- | --- | --- | --- | --- | --- | --- | --- | --- | --- | --- | --- | --- |
| Arm1 | One sample T-test | normal |  | -0.359 | 0.721 | NA | -1.123 | 0.276 | NA | 0.988 | 0.336 | NA | -0.469 | 0.644 | NA |
| Arm2 | One sample T-test | normal |  | 1.465 | 0.148 | NA | 1.578 | 0.132 | NA | -0.534 | 0.600 | NA | 1.354 | 0.193 | NA |
| Arm3 | One sample T-test | normal |  | -1.338 | 0.186 | NA | -0.772 | 0.450 | NA | -0.312 | 0.759 | NA | -1.262 | 0.223 | NA |

|  | test | data structure | WT | Het | KO | genotype |  |  | cohort |  |  | genotype x cohort |  |  | pairwise comparisons |  |  |
| --- | --- | --- | --- | --- | --- | --- | --- | --- | --- | --- | --- | --- | --- | --- | --- | --- | --- |
|  |  |  |  |  |  | F | p-value | power | F | p-value | power | F | p-value | power | WT vs Het | WT vs KO | Het vs KO |
| 0-15 min, total number of choices | 2-way ANOVA | normal | 43.42 ± 3.25 | 40.26 ± 2.54 | 38.47 ± 2.75 | 0.612 | 0.546 | 0.147 | 0.164 | 0.687 | 0.068 | 2.244 | 0.116 | 0.437 | - | - | - |
| 0-15 min, number of correct choice | 2-way ANOVA | normal | 57.46 ± 1.59 | 60.68 ± 1.93 | 57.52 ± 1.61 | 1.227 | 0.302 | 0.256 | 2.987 | 0.090 | 0.396 | 0.927 | 0.402 | 0.202 | - | - | - |
| 0-15 min, number of type 1 errors | 2-way ANOVA | normal | 37.8 ± 1.43 | 34.09 ± 1.59 | 38.29 ± 1.79 | 2.296 | 0.111 | 0.445 | 0.300 | 0.586 | 0.084 | 5.165 | 0.009 | 0.804 | - | - | - |
| 0-15 min, number of type 2 errors | 2-way ANOVA | non normal | 4.04 ± 1.11 | 5.47 ± 1.02 | 4.51 ± 1.07 | 0.397 | 0.674 | 0.111 | 4.402 | 0.041 | 0.539 | 3.449 | 0.039 | 0.621 | - | - | - |

| Fear conditioning |  |  |  |  |  |  |  |  |  |
| --- | --- | --- | --- | --- | --- | --- | --- | --- | --- |
| Training, repeated measures | Cohorts 1 and 2 |  |  |  |  |  |  |  |  |
|  | test | data structure |  | F | p-value | power | WT vs Het | WT vs KO | Het vs KO |
| - time effect | repeated measures | sphericity violated |  | 43.998 | 0.000 | 1.000 | - | - | - |
| - time x genotype effect | repeated measures | sphericity violated |  | 3.194 | 0.002 | 0.970 | - | - | - |
| - genotype effect | repeated measures | sphericity violated |  | 14.505 | 0.000 | 0.998 | 0.809 | 0.000 | 0.000 |
| - cohort effect | repeated measures | sphericity violated |  | 12.351 | 0.001 | 0.932 | - | - | - |
| - time x genotype x cohort effect | repeated measures | sphericity violated |  | 0.602 | 0.782 | 0.281 | - | - | - |
| - genotype x cohort effect | repeated measures | sphericity violated |  | 0.494 | 0.613 | 0.127 | - | - | - |

| Training, individual time bins | Cohorts 1 and 2 |  |  |  |  |  |  |  |  |  |  |  |  |  |  |  |  |
| --- | --- | --- | --- | --- | --- | --- | --- | --- | --- | --- | --- | --- | --- | --- | --- | --- | --- |
|  | test | data structure | WT | Het | KO | genotype |  |  | cohort |  |  | genotype x cohort |  |  | pairwise comparisons |  |  |
|  |  |  |  |  |  | F | p-value | power | F | p-value | power | F | p-value | power | WT vs Het | WT vs KO | Het vs KO |
| Training habituation | 2-way ANOVA | non normal | 16.85 ± 2.93 | 10.45 ± 2.37 | 21.6 ± 5.3 | 2.081 | 0.135 | 0.408 | 0.015 | 0.903 | 0.052 | 0.061 | 0.941 | 0.059 | - | - | - |
| Training Pre-tone 0-120 | 2-way ANOVA | non normal | 10.45 ± 1.72 | 8.15 ± 1.43 | 21.01 ± 3.97 | 6.546 | 0.003 | 0.892 | 0.330 | 0.568 | 0.087 | 0.041 | 0.960 | 0.056 | 0.820 | 0.021 | 0.004 |
| Training Tone/shock120-140 | 2-way ANOVA | non normal | 10.58 ± 3.26 | 6.85 ± 2.82 | 19.94 ± 6.06 | 2.361 | 0.105 | 0.456 | 0.020 | 0.887 | 0.052 | 0.199 | 0.820 | 0.079 | - | - | - |
| Training Post-tone140-260 | 2-way ANOVA | non normal | 19.74 ± 3.73 | 18.83 ± 3.86 | 47.68 ± 6.71 | 13.149 | 0.000 | 0.996 | 5.506 | 0.023 | 0.634 | 2.222 | 0.119 | 0.433 | 0.990 | 0.000 | 0.000 |
| Training Tone/shock 260-280 | 2-way ANOVA | non normal | 15.07 ± 4.7 | 24.58 ± 5.22 | 47.23 ± 7.47 | 7.613 | 0.001 | 0.934 | 0.026 | 0.871 | 0.053 | 0.762 | 0.472 | 0.173 | 0.507 | 0.001 | 0.027 |
| Training Tone/shock 260-280 | 2-way ANOVA | non normal | 31.06 ± 5.14 | 37.88 ± 6.6 | 65.23 ± 6.72 | 12.505 | 0.000 | 0.995 | 18.006 | 0.000 | 0.986 | 0.640 | 0.532 | 0.151 | 0.650 | 0.000 | 0.002 |
| Training Tone/shock 400-420 | 2-way ANOVA | non normal | 31.03 ± 5.86 | 40.59 ± 7.61 | 65.21 ± 5.94 | 9.728 | 0.000 | 0.977 | 12.565 | 0.001 | 0.935 | 0.061 | 0.941 | 0.059 | 0.503 | 0.001 | 0.015 |
| Training Post-tone 420-540 | 2-way ANOVA | non normal | 36.71 ± 6.53 | 47.61 ± 7.36 | 61.74 ± 6.78 | 7.880 | 0.001 | 0.942 | 45.207 | 0.000 | 1.000 | 0.135 | 0.874 | 0.070 | 0.303 | 0.003 | 0.139 |

| Context, repeated measures | test | data structure |  | F | p-value | power | WT vs Het | WT vs KO | Het vs KO |
| --- | --- | --- | --- | --- | --- | --- | --- | --- | --- |
| - time effect | repeated measures | sphericity assumed |  | 4.558 | 0.004 | 0.880 | - | - | - |
| - time x genotype effect | repeated measures | sphericity assumed |  | 0.675 | 0.670 | 0.262 | - | - | - |
| - genotype effect | repeated measures | sphericity assumed |  | 1.788 | 0.178 | 0.357 | - | - | - |
| - cohort effect | repeated measures | sphericity assumed |  | 0.542 | 0.465 | 0.112 | - | - | - |
| - time x genotype x cohort effect | repeated measures | sphericity assumed |  | 0.918 | 0.481 | 0.355 | - | - | - |
| - genotype x cohort effect | repeated measures | sphericity assumed |  | 1.026 | 0.366 | 0.219 | - | - | - |

| Context, individual time bins | test | data structure | WT | Het | KO | genotype |  |  | cohort |  |  | genotype x cohort |  |  | pairwise comparisons |  |  |
| --- | --- | --- | --- | --- | --- | --- | --- | --- | --- | --- | --- | --- | --- | --- | --- | --- | --- |
|  |  |  |  |  |  | F | p-value | power | F | p-value | power | F | p-value | power | WT vs Het | WT vs KO | Het vs KO |
| Context 0-60 | 2-way ANOVA | non normal | 63.09 ± 5.13 | 57.19 ± 5.63 | 44.3 ± 6.21 | 2.643 | 0.081 | 0.502 | 0.582 | 0.449 | 0.116 | 0.558 | 0.576 | 0.137 | 0.750 | 0.063 | 0.261 |
| Context 60-120 | 2-way ANOVA | non normal | 66.34 ± 6.83 | 66 ± 6.94 | 59.6 ± 7.41 | 0.230 | 0.795 | 0.084 | 0.676 | 0.415 | 0.127 | 2.233 | 0.118 | 0.435 | - | - | - |
| Context 120-180 | 2-way ANOVA | non normal | 62.65 ± 7.14 | 62.11 ± 6.47 | 42.81 ± 7.83 | 2.263 | 0.114 | 0.440 | 0.392 | 0.534 | 0.094 | 0.958 | 0.390 | 0.207 | - | - | - |
| Context 180-240 | 2-way ANOVA | non normal | 56.12 ± 6.56 | 54.45 ± 7.58 | 43.33 ± 6.29 | 0.944 | 0.396 | 0.205 | 0.073 | 0.788 | 0.058 | 0.109 | 0.897 | 0.066 | - | - | - |
| Context mean | 2-way ANOVA | non normal | 62.05 ± 5.57 | 59.94 ± 5.31 | 47.51 ± 5.84 | 1.788 | 0.178 | 0.357 | 0.542 | 0.465 | 0.112 | 1.026 | 0.366 | 0.219 | - | - | - |

| Cued, repeated measures | test | data structure |  | F | p-value | power | WT vs Het | WT vs KO | Het vs KO |
| --- | --- | --- | --- | --- | --- | --- | --- | --- | --- |
| - time effect | repeated measures | sphericity violated |  | 25.753 | 0.000 | 1.000 | - | - | - |
| - time x genotype effect | repeated measures | sphericity violated |  | 3.101 | 0.002 | 0.968 | - | - | - |
| - genotype effect | repeated measures | sphericity violated |  | 5.657 | 0.006 | 0.841 | 0.645 | 0.007 | 0.065 |
| - cohort effect | repeated measures | sphericity violated |  | 4.255 | 0.044 | 0.525 | - | - | - |
| - time x genotype x cohort effect | repeated measures | sphericity violated |  | 4.116 | 0.000 | 0.995 | - | - | - |
| - genotype x cohort effect | repeated measures | sphericity violated |  | 1.616 | 0.209 | 0.326 | - | - | - |

| Cued, individual time bins | test | data structure | WT | Het | KO | genotype |  |  | cohort |  |  | genotype x cohort |  |  | pairwise comparisons |  |  |
| --- | --- | --- | --- | --- | --- | --- | --- | --- | --- | --- | --- | --- | --- | --- | --- | --- | --- |
|  |  |  |  |  |  | F | p-value | power | F | p-value | power | F | p-value | power | WT vs Het | WT vs KO | Het vs KO |
| Cued Pre-tone 0-60 | 2-way ANOVA | non normal | 1.64 ± 1.14 | 0.4 ± 0.28 | 4.9 ± 2.59 | 1.897 | 0.160 | 0.376 | 5.996 | 0.018 | 0.671 | 1.527 | 0.227 | 0.310 | 0.841 | 0.311 | 0.114 |
| Cued Pre-tone 60-120 | 2-way ANOVA | non normal | 0.84 ± 0.38 | 1.96 ± 0.89 | 2.37 ± 0.88 | 1.056 | 0.355 | 0.225 | 18.576 | 0.000 | 0.988 | 0.747 | 0.479 | 0.170 | 0.461 | 0.238 | 0.896 |
| Cued Tone 120-140 | 2-way ANOVA | non normal | 10.94 ± 4.37 | 12.14 ± 4.53 | 25.23 ± 6.38 | 3.005 | 0.058 | 0.558 | 7.144 | 0.010 | 0.746 | 1.298 | 0.282 | 0.268 | 0.984 | 0.106 | 0.150 |
| Cued Post-tone 140-200 | 2-way ANOVA | non normal | 8.52 ± 2.11 | 7.41 ± 1.76 | 13.09 ± 3.74 | 1.367 | 0.264 | 0.281 | 1.610 | 0.210 | 0.238 | 0.118 | 0.889 | 0.067 | - | - | - |
| Cued Post-tone 200-260 | 2-way ANOVA | non normal | 2.29 ± 1.03 | 6.59 ± 1.53 | 10.03 ± 4.8 | 1.551 | 0.222 | 0.314 | 1.595 | 0.212 | 0.236 | 1.203 | 0.309 | 0.251 | - | - | - |
| Cued Tone 260-280 | 2-way ANOVA | non normal | 13.87 ± 5.15 | 19.36 ± 5.92 | 39.08 ± 7.75 | 7.219 | 0.002 | 0.921 | 15.352 | 0.000 | 0.970 | 7.888 | 0.001 | 0.942 | 0.733 | 0.003 | 0.025 |
| Cued Post-tone 280-340 | 2-way ANOVA | non normal | 8.92 ± 2.71 | 19.61 ± 5.1 | 26.7 ± 6.01 | 3.891 | 0.027 | 0.677 | 5.448 | 0.024 | 0.629 | 1.180 | 0.316 | 0.247 | 0.240 | 0.024 | 0.527 |
| Cued Post-tone 340-400 | 2-way ANOVA | non normal | 5.09 ± 1.34 | 9.96 ± 2.78 | 20.99 ± 4.81 | 6.189 | 0.004 | 0.874 | 1.570 | 0.216 | 0.233 | 1.068 | 0.351 | 0.227 | 0.549 | 0.003 | 0.054 |

When looking at the spontaneous alternation behavior in the Y-maze, no differences were observed between the genotypes in any of the background strains regarding either the total number of choices, the percentage of correct choices or the percentage of errors (Figure 9A). Moreover, no arm preference was seen for any of the groups.

**Figure 9:**
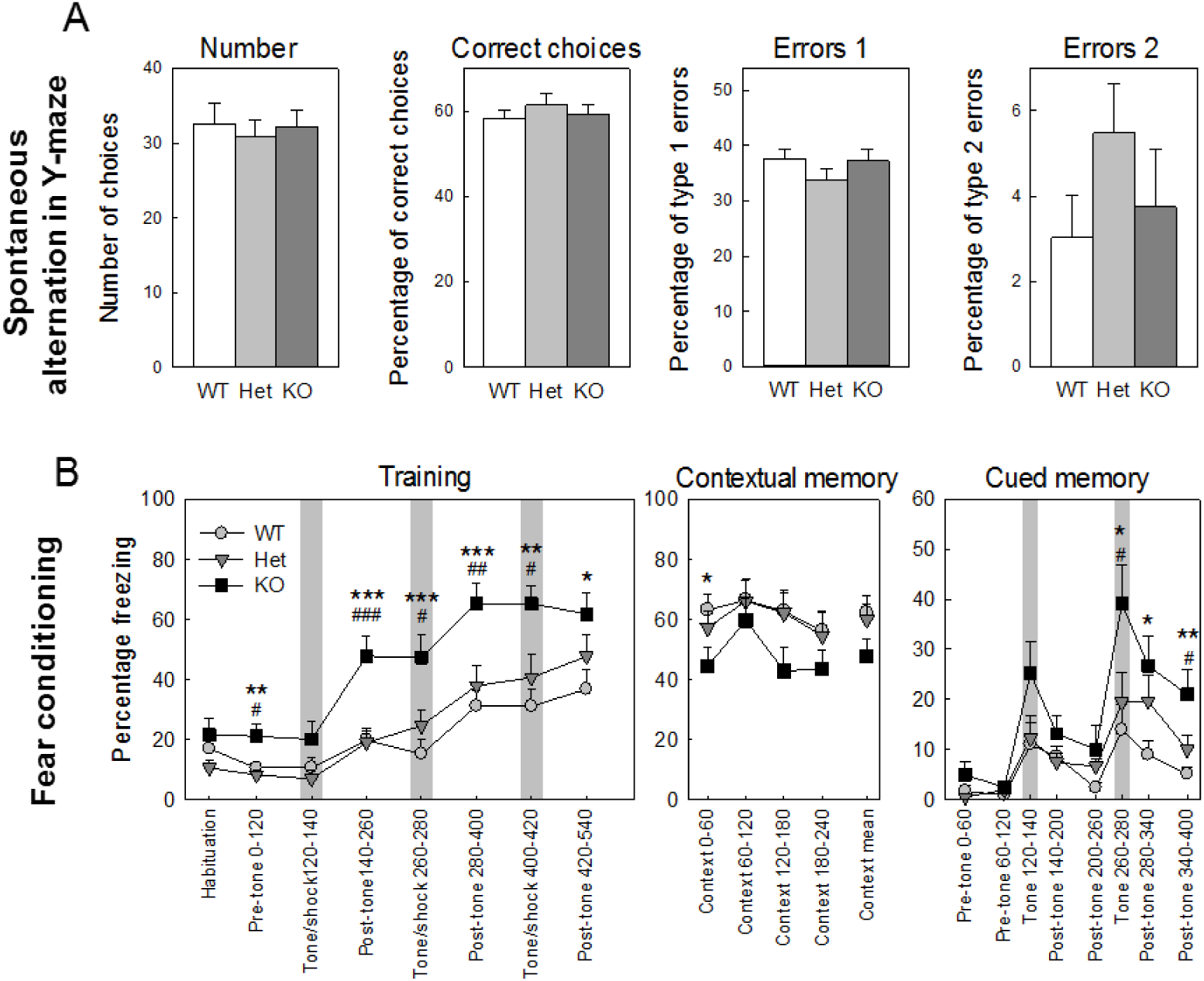
Learning and memory in *Shank3^Δ4-22^-*deficient mice. **(A)** Working memory in Y-maze measured by spontaneous alternation behavior. All genotypes showed comparable number of arm choices, percentage of correct choices (3-way alternation), type 1 error (three consecutive choices where the first and third choices are identical) or type 2 error (three consecutive choices where the second and third choices are identical). **(B)** Contextual and cued fear conditioning in *Shank3* mice. A higher percentage of freezing was observed in *Shank3^Δ4-^ 22* homozygous mice compared to wild-type and heterozygous animals on day one. While the difference was already present before the sound-shocks associations, it was strongly increased post-training. No genotype differences were detected in freezing scores in the post-training session on day 1. Opposite results were observed for contextual conditioning (day 2) and cued conditioning (day 3): *Shank3^Δ4-22^* homozygous mice showed an impairment of contextual learning compared to their wild-type and heterozygous littermates but and enhancement of freezing post-cues during the cued testing. WT, wild-type mice; Het, heterozygous mice; KO, homozygous knockout mice. *: WT vs KO; o: WT vs Het, #: Het vs KO. *: p<0.05, **: p<0.1, ***: p<0.001.

In the training session of the fear conditioning test minimal levels of freezing behavior were seen for all the genotypes during the 5-minute habituation period, however, while this percentage of spontaneous freezing decreased before the presentations of cue–shock pairings for the *Shank3^Δ4-22^* wild-type and heterozygotes it remained at significantly higher level for *Shank3^Δ4-22^* homozygous mice. A significant genotype effect was then found during the training session in post-shock freezing, with *Shank3^Δ4-22^* homozygous mice displaying higher levels of freezing compared with wild-type and heterozygous mice (Figure 9B left panel). The opposite was observed during contextual recall where even if all the mice freeze significantly more than during the habituation of the training sessions a trend toward a decrease (significant during the first minute) of freezing was observed for *Shank3^Δ4-22^* homozygous mice compared to wild-type or heterozygous littermates (Figure 9B, middle panel). An increase of freezing was seen in both during and after the cue presentation (trend for the first cue, significant during and after the second cue) *Shank3^Δ4-22^* homozygous mice (Figure 9B, right panel).

### Anxiety-related behaviors in *Shank3^Δ4-22^*-deficient mice

Anxiety-like behaviors were monitored in the open-field and in the elevated zero-maze, and detailed results are displayed in Table 13.

**Table 13:**
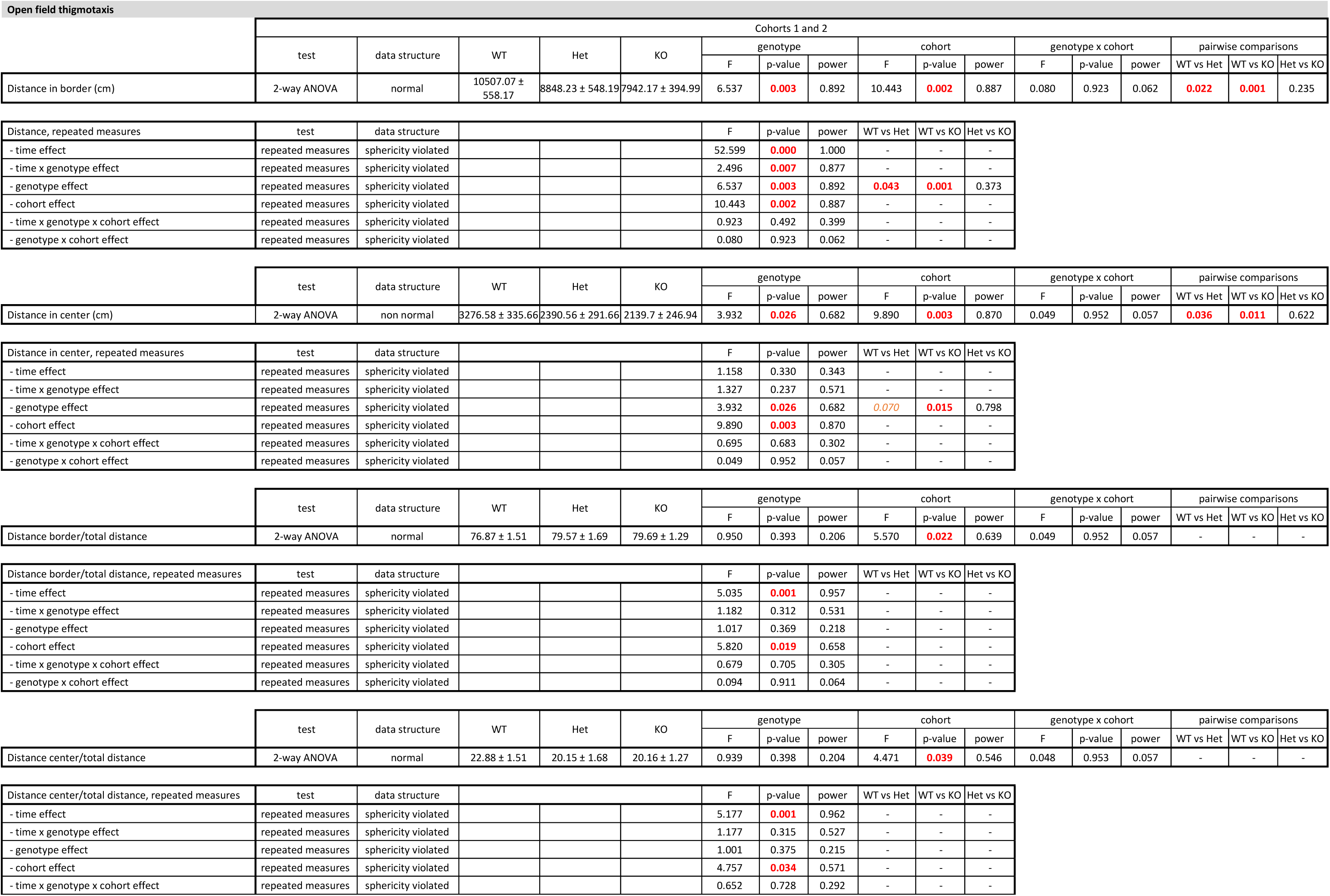
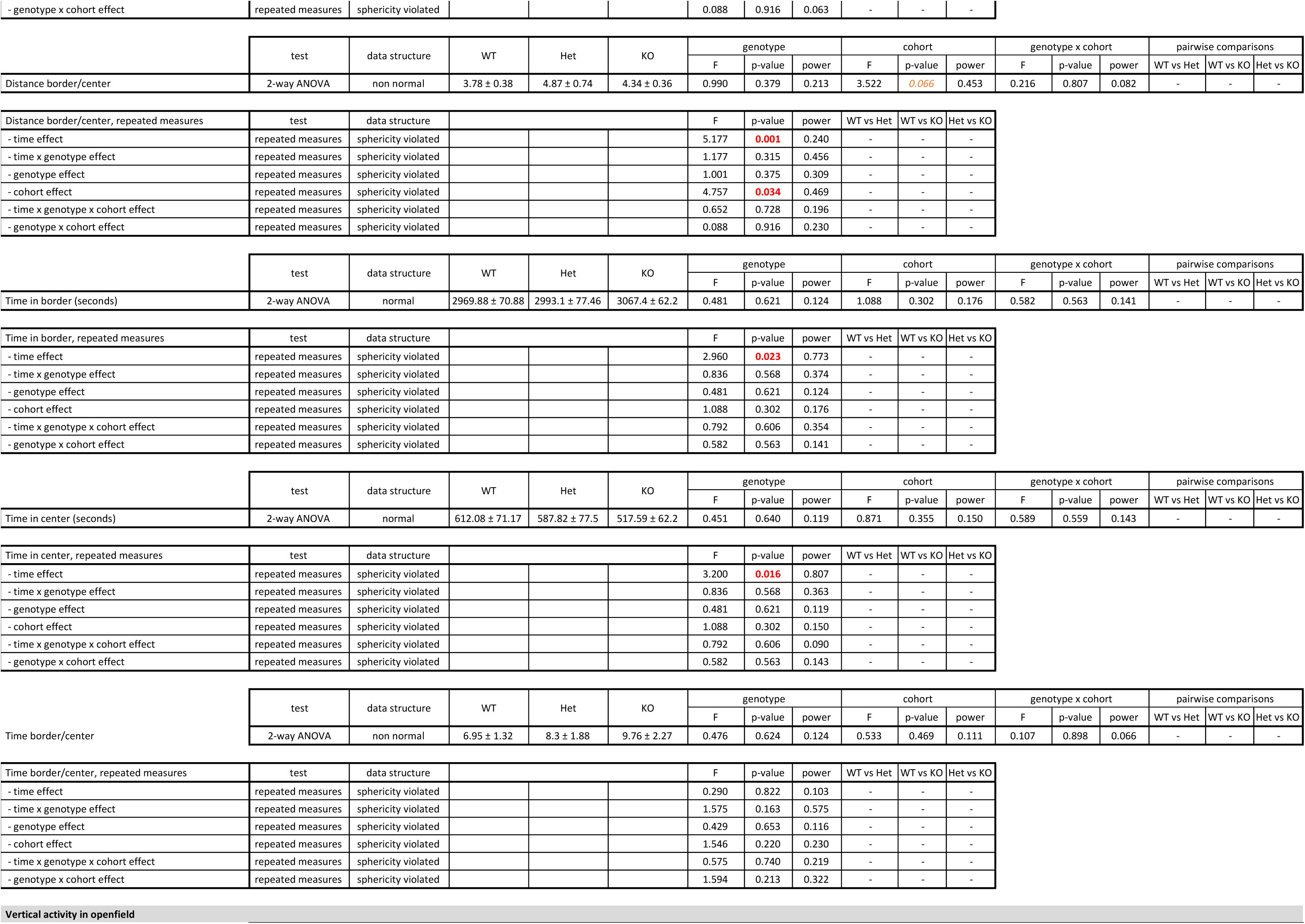
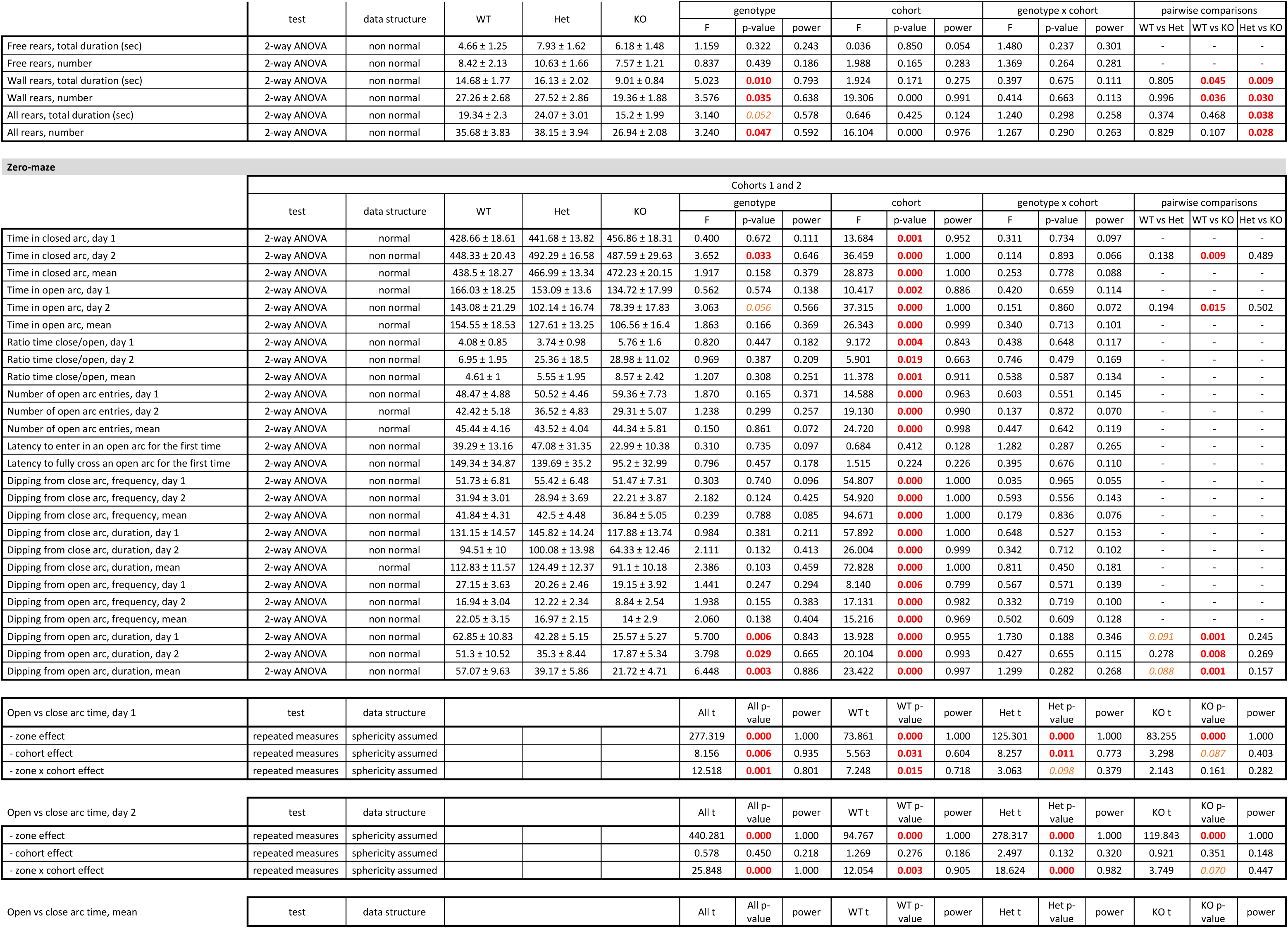
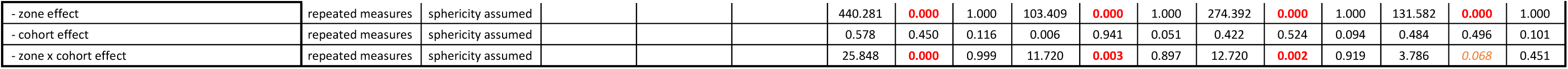
Detailed results and statistical analyses related to anxiety-like behaviors. WT, wild-type mice; Het, heterozygous mice; KO, homozygous knockout mice. Group values are reported as means ± s.e.m. Red font indicates significant results (p<0.05), orange font indicates trends (0.1<p<0.05). Individual results and statistical analyses for cohorts 1 and 2 are available in Extended Table 13-1

| Open field thigmotaxis |  |  |  |  |  |  |  |  |  |  |  |  |  |  |  |  |  |
| --- | --- | --- | --- | --- | --- | --- | --- | --- | --- | --- | --- | --- | --- | --- | --- | --- | --- |
|  | Cohorts 1 and 2 |  |  |  |  |  |  |  |  |  |  |  |  |  |  |  |  |
|  | test | data structure | WT | Het | KO | genotype |  |  | cohort |  |  | genotype x cohort |  |  | pairwise comparisons |  |  |
|  |  |  |  |  |  | F | p-value | power | F | p-value | power | F | p-value | power | WT vs Het | WT vs KO | Het vs KO |
| Distance in border (cm) | 2-way ANOVA | normal | 10507.07 ± 558.17 | 8848.23 ± 548.19 | 7942.17 ± 394.99 | 6.537 | **0.003** | 0.892 | 10.443 | **0.002** | 0.887 | 0.080 | 0.923 | 0.062 | **0.022** | **0.001** | 0.235 |
| Distance, repeated measures | test | data structure |  |  |  | F | p-value | power | WT vs Het | WT vs KO | Het vs KO |  |  |  |  |  |  |
| - time effect | repeated measures | sphericity violated |  |  |  | 52.599 | **0.000** | 1.000 | - | - | - |  |  |  |  |  |  |
| - time x genotype effect | repeated measures | sphericity violated |  |  |  | 2.496 | **0.007** | 0.877 | - | - | - |  |  |  |  |  |  |
| - genotype effect | repeated measures | sphericity violated |  |  |  | 6.537 | **0.003** | 0.892 | **0.043** | **0.001** | 0.373 |  |  |  |  |  |  |
| - cohort effect | repeated measures | sphericity violated |  |  |  | 10.443 | **0.002** | 0.887 | - | - | - |  |  |  |  |  |  |
| - time x genotype x cohort effect | repeated measures | sphericity violated |  |  |  | 0.923 | 0.492 | 0.399 | - | - | - |  |  |  |  |  |  |
| - genotype x cohort effect | repeated measures | sphericity violated |  |  |  | 0.080 | 0.923 | 0.062 | - | - | - |  |  |  |  |  |  |
|  | test | data structure | WT | Het | KO | genotype |  |  | cohort |  |  | genotype x cohort |  |  | pairwise comparisons |  |  |
|  |  |  |  |  |  | F | p-value | power | F | p-value | power | F | p-value | power | WT vs Het | WT vs KO | Het vs KO |
|  |  |  |  |  |  | F | p-value | power | F | p-value | power | F | p-value | power | WT vs Het | WT vs KO | Het vs KO |
| Distance in center (cm) | 2-way ANOVA | non normal | 3276.58 ± 335.66 | 2390.56 ± 291.66 | 2139.7 ± 246.94 | 3.932 | **0.026** | 0.682 | 9.890 | **0.003** | 0.870 | 0.049 | 0.952 | 0.057 | **0.036** | **0.011** | 0.622 |
| Distance in center, repeated measures | test | data structure |  |  |  | F | p-value | power | WT vs Het | WT vs KO | Het vs KO |  |  |  |  |  |  |
| - time effect | repeated measures | sphericity violated |  |  |  | 1.158 | 0.330 | 0.343 | - | - | - |  |  |  |  |  |  |
| - time x genotype effect | repeated measures | sphericity violated |  |  |  | 1.327 | 0.237 | 0.571 | - | - | - |  |  |  |  |  |  |
| - genotype effect | repeated measures | sphericity violated |  |  |  | 3.932 | **0.026** | 0.682 | **0.070** | **0.015** | 0.798 |  |  |  |  |  |  |
| - cohort effect | repeated measures | sphericity violated |  |  |  | 9.890 | **0.003** | 0.870 | - | - | - |  |  |  |  |  |  |
| - time x genotype x cohort effect | repeated measures | sphericity violated |  |  |  | 0.695 | 0.683 | 0.302 | - | - | - |  |  |  |  |  |  |
| - genotype x cohort effect | repeated measures | sphericity violated |  |  |  | 0.049 | 0.952 | 0.057 | - | - | - |  |  |  |  |  |  |
|  | test | data structure | WT | Het | KO | genotype |  |  | cohort |  |  | genotype x cohort |  |  | pairwise comparisons |  |  |
|  |  |  |  |  |  | F | p-value | power | F | p-value | power | F | p-value | power | WT vs Het | WT vs KO | Het vs KO |
|  |  |  |  |  |  | F | p-value | power | F | p-value | power | F | p-value | power | WT vs Het | WT vs KO | Het vs KO |
| Distance border/total distance | 2-way ANOVA | normal | 76.87 ± 1.51 | 79.57 ± 1.69 | 79.69 ± 1.29 | 0.950 | 0.393 | 0.206 | 5.570 | **0.022** | 0.639 | 0.049 | 0.952 | 0.057 | - | - | - |
| Distance border/total distance, repeated measures | test | data structure |  |  |  | F | p-value | power | WT vs Het | WT vs KO | Het vs KO |  |  |  |  |  |  |
| - time effect | repeated measures | sphericity violated |  |  |  | 5.035 | **0.001** | 0.957 | - | - | - |  |  |  |  |  |  |
| - time x genotype effect | repeated measures | sphericity violated |  |  |  | 1.182 | 0.312 | 0.531 | - | - | - |  |  |  |  |  |  |
| - genotype effect | repeated measures | sphericity violated |  |  |  | 1.017 | 0.369 | 0.218 | - | - | - |  |  |  |  |  |  |
| - cohort effect | repeated measures | sphericity violated |  |  |  | 5.820 | **0.019** | 0.658 | - | - | - |  |  |  |  |  |  |
| - time x genotype x cohort effect | repeated measures | sphericity violated |  |  |  | 0.679 | 0.705 | 0.305 | - | - | - |  |  |  |  |  |  |
| - genotype x cohort effect | repeated measures | sphericity violated |  |  |  | 0.094 | 0.911 | 0.064 | - | - | - |  |  |  |  |  |  |
|  | test | data structure | WT | Het | KO | genotype |  |  | cohort |  |  | genotype x cohort |  |  | pairwise comparisons |  |  |
|  |  |  |  |  |  | F | p-value | power | F | p-value | power | F | p-value | power | WT vs Het | WT vs KO | Het vs KO |
|  |  |  |  |  |  | F | p-value | power | F | p-value | power | F | p-value | power | WT vs Het | WT vs KO | Het vs KO |
| Distance center/total distance | 2-way ANOVA | normal | 22.88 ± 1.51 | 20.15 ± 1.68 | 20.16 ± 1.27 | 0.939 | 0.398 | 0.204 | 4.471 | **0.039** | 0.546 | 0.048 | 0.953 | 0.057 | - | - | - |
| Distance center/total distance, repeated measures | test | data structure |  |  |  | F | p-value | power | WT vs Het | WT vs KO | Het vs KO |  |  |  |  |  |  |
| - time effect | repeated measures | sphericity violated |  |  |  | 5.177 | **0.001** | 0.962 | - | - | - |  |  |  |  |  |  |
| - time x genotype effect | repeated measures | sphericity violated |  |  |  | 1.177 | 0.315 | 0.527 | - | - | - |  |  |  |  |  |  |
| - genotype effect | repeated measures | sphericity violated |  |  |  | 1.001 | 0.375 | 0.215 | - | - | - |  |  |  |  |  |  |
| - cohort effect | repeated measures | sphericity violated |  |  |  | 4.757 | **0.034** | 0.571 | - | - | - |  |  |  |  |  |  |
| - time x genotype x cohort effect | repeated measures | sphericity violated |  |  |  | 0.652 | 0.728 | 0.292 | - | - | - |  |  |  |  |  |  |

|  |  |  |  |  |  |  |  |  |  |  |  |
| --- | --- | --- | --- | --- | --- | --- | --- | --- | --- | --- | --- |
| - genotype x cohort effect | repeated measures | sphericity violated |  |  |  | 0.088 | 0.916 | 0.063 | - | - | - |

|  | test | data structure | WT | Het | KO | genotype |  |  | cohort |  |  | genotype x cohort |  |  | pairwise comparisons |  |  |
| --- | --- | --- | --- | --- | --- | --- | --- | --- | --- | --- | --- | --- | --- | --- | --- | --- | --- |
|  |  |  |  |  |  | F | p-value | power | F | p-value | power | F | p-value | power | WT vs Het | WT vs KO | Het vs KO |
| Distance border/center | 2-way ANOVA | non normal | 3.78 ± 0.38 | 4.87 ± 0.74 | 4.34 ± 0.36 | 0.990 | 0.379 | 0.213 | 3.522 | 0.066 | 0.453 | 0.216 | 0.807 | 0.082 | - | - | - |

| Distance border/center, repeated measures | test | data structure |  |  |  | F | p-value | power | WT vs Het | WT vs KO | Het vs KO |
| --- | --- | --- | --- | --- | --- | --- | --- | --- | --- | --- | --- |
| - time effect | repeated measures | sphericity violated |  |  |  | 5.177 | 0.001 | 0.240 | - | - | - |
| - time x genotype effect | repeated measures | sphericity violated |  |  |  | 1.177 | 0.315 | 0.456 | - | - | - |
| - genotype effect | repeated measures | sphericity violated |  |  |  | 1.001 | 0.375 | 0.309 | - | - | - |
| - cohort effect | repeated measures | sphericity violated |  |  |  | 4.757 | 0.034 | 0.469 | - | - | - |
| - time x genotype x cohort effect | repeated measures | sphericity violated |  |  |  | 0.652 | 0.728 | 0.196 | - | - | - |
| - genotype x cohort effect | repeated measures | sphericity violated |  |  |  | 0.088 | 0.916 | 0.230 | - | - | - |

|  | test | data structure | WT | Het | KO | genotype |  |  | cohort |  |  | genotype x cohort |  |  | pairwise comparisons |  |  |
| --- | --- | --- | --- | --- | --- | --- | --- | --- | --- | --- | --- | --- | --- | --- | --- | --- | --- |
|  |  |  |  |  |  | F | p-value | power | F | p-value | power | F | p-value | power | WT vs Het | WT vs KO | Het vs KO |
| Time in border (seconds) | 2-way ANOVA | normal | 2969.88 ± 70.88 | 2993.1 ± 77.46 | 3067.4 ± 62.2 | 0.481 | 0.621 | 0.124 | 1.088 | 0.302 | 0.176 | 0.582 | 0.563 | 0.141 | - | - | - |

| Time in border, repeated measures | test | data structure |  |  |  | F | p-value | power | WT vs Het | WT vs KO | Het vs KO |
| --- | --- | --- | --- | --- | --- | --- | --- | --- | --- | --- | --- |
| - time effect | repeated measures | sphericity violated |  |  |  | 2.960 | 0.023 | 0.773 | - | - | - |
| - time x genotype effect | repeated measures | sphericity violated |  |  |  | 0.836 | 0.568 | 0.374 | - | - | - |
| - genotype effect | repeated measures | sphericity violated |  |  |  | 0.481 | 0.621 | 0.124 | - | - | - |
| - cohort effect | repeated measures | sphericity violated |  |  |  | 1.088 | 0.302 | 0.176 | - | - | - |
| - time x genotype x cohort effect | repeated measures | sphericity violated |  |  |  | 0.792 | 0.606 | 0.354 | - | - | - |
| - genotype x cohort effect | repeated measures | sphericity violated |  |  |  | 0.582 | 0.563 | 0.141 | - | - | - |

|  | test | data structure | WT | Het | KO | genotype |  |  | cohort |  |  | genotype x cohort |  |  | pairwise comparisons |  |  |
| --- | --- | --- | --- | --- | --- | --- | --- | --- | --- | --- | --- | --- | --- | --- | --- | --- | --- |
|  |  |  |  |  |  | F | p-value | power | F | p-value | power | F | p-value | power | WT vs Het | WT vs KO | Het vs KO |
| Time in center (seconds) | 2-way ANOVA | normal | 612.08 ± 71.17 | 587.82 ± 77.5 | 517.59 ± 62.2 | 0.451 | 0.640 | 0.119 | 0.871 | 0.355 | 0.150 | 0.589 | 0.559 | 0.143 | - | - | - |

| Time in center, repeated measures | test | data structure |  |  |  | F | p-value | power | WT vs Het | WT vs KO | Het vs KO |
| --- | --- | --- | --- | --- | --- | --- | --- | --- | --- | --- | --- |
| - time effect | repeated measures | sphericity violated |  |  |  | 3.200 | 0.016 | 0.807 | - | - | - |
| - time x genotype effect | repeated measures | sphericity violated |  |  |  | 0.836 | 0.568 | 0.363 | - | - | - |
| - genotype effect | repeated measures | sphericity violated |  |  |  | 0.481 | 0.621 | 0.119 | - | - | - |
| - cohort effect | repeated measures | sphericity violated |  |  |  | 1.088 | 0.302 | 0.150 | - | - | - |
| - time x genotype x cohort effect | repeated measures | sphericity violated |  |  |  | 0.792 | 0.606 | 0.090 | - | - | - |
| - genotype x cohort effect | repeated measures | sphericity violated |  |  |  | 0.582 | 0.563 | 0.143 | - | - | - |

|  | test | data structure | WT | Het | KO | genotype |  |  | cohort |  |  | genotype x cohort |  |  | pairwise comparisons |  |  |
| --- | --- | --- | --- | --- | --- | --- | --- | --- | --- | --- | --- | --- | --- | --- | --- | --- | --- |
|  |  |  |  |  |  | F | p-value | power | F | p-value | power | F | p-value | power | WT vs Het | WT vs KO | Het vs KO |
| Time border/center | 2-way ANOVA | non normal | 6.95 ± 1.32 | 8.3 ± 1.88 | 9.76 ± 2.27 | 0.476 | 0.624 | 0.124 | 0.533 | 0.469 | 0.111 | 0.107 | 0.898 | 0.066 | - | - | - |

| Time border/center, repeated measures | test | data structure |  |  |  | F | p-value | power | WT vs Het | WT vs KO | Het vs KO |
| --- | --- | --- | --- | --- | --- | --- | --- | --- | --- | --- | --- |
| - time effect | repeated measures | sphericity violated |  |  |  | 0.290 | 0.822 | 0.103 | - | - | - |
| - time x genotype effect | repeated measures | sphericity violated |  |  |  | 1.575 | 0.163 | 0.575 | - | - | - |
| - genotype effect | repeated measures | sphericity violated |  |  |  | 0.429 | 0.653 | 0.116 | - | - | - |
| - cohort effect | repeated measures | sphericity violated |  |  |  | 1.546 | 0.220 | 0.230 | - | - | - |
| - time x genotype x cohort effect | repeated measures | sphericity violated |  |  |  | 0.575 | 0.740 | 0.219 | - | - | - |
| - genotype x cohort effect | repeated measures | sphericity violated |  |  |  | 1.594 | 0.213 | 0.322 | - | - | - |

| Vertical activity in openfield |
| --- |
| Cohorts 1 and 2 |

|  | test | data structure | WT | Het | KO | genotype |  |  | cohort |  |  | genotype x cohort |  |  | pairwise comparisons |  |  |
| --- | --- | --- | --- | --- | --- | --- | --- | --- | --- | --- | --- | --- | --- | --- | --- | --- | --- |
|  |  |  |  |  |  | F | p-value | power | F | p-value | power | F | p-value | power | WT vs Het | WT vs KO | Het vs KO |
| Free rears, total duration (sec) | 2-way ANOVA | non normal | 4.66 ± 1.25 | 7.93 ± 1.62 | 6.18 ± 1.48 | 1.159 | 0.322 | 0.243 | 0.036 | 0.850 | 0.054 | 1.480 | 0.237 | 0.301 | - | - | - |
| Free rears, number | 2-way ANOVA | non normal | 8.42 ± 2.13 | 10.63 ± 1.66 | 7.57 ± 1.21 | 0.837 | 0.439 | 0.186 | 1.988 | 0.165 | 0.283 | 1.369 | 0.264 | 0.281 | - | - | - |
| Wall rears, total duration (sec) | 2-way ANOVA | non normal | 14.68 ± 1.77 | 16.13 ± 2.02 | 9.01 ± 0.84 | 5.023 | **0.010** | 0.793 | 1.924 | 0.171 | 0.275 | 0.397 | 0.675 | 0.111 | 0.805 | **0.045** | **0.009** |
| Wall rears, number | 2-way ANOVA | non normal | 27.26 ± 2.68 | 27.52 ± 2.86 | 19.36 ± 1.88 | 3.576 | **0.035** | 0.638 | 19.306 | 0.000 | 0.991 | 0.414 | 0.663 | 0.113 | 0.996 | **0.036** | **0.030** |
| All rears, total duration (sec) | 2-way ANOVA | non normal | 19.34 ± 2.3 | 24.07 ± 3.01 | 15.2 ± 1.99 | 3.140 | **0.052** | 0.578 | 0.646 | 0.425 | 0.124 | 1.240 | 0.298 | 0.258 | 0.374 | 0.468 | **0.038** |
| All rears, number | 2-way ANOVA | non normal | 35.68 ± 3.83 | 38.15 ± 3.94 | 26.94 ± 2.08 | 3.240 | **0.047** | 0.592 | 16.104 | 0.000 | 0.976 | 1.267 | 0.290 | 0.263 | 0.829 | 0.107 | **0.028** |

Zero-maze
|  | test | data structure | WT | Het | KO | Cohorts 1 and 2 |  |  |  |  |  |  |  |  |  |  |  |
| --- | --- | --- | --- | --- | --- | --- | --- | --- | --- | --- | --- | --- | --- | --- | --- | --- | --- |
|  |  |  |  |  |  | genotype |  |  | cohort |  |  | genotype x cohort |  |  | pairwise comparisons |  |  |
|  |  |  |  |  |  | F | p-value | power | F | p-value | power | F | p-value | power | WT vs Het | WT vs KO | Het vs KO |
| Time in closed arc, day 1 | 2-way ANOVA | normal | 428.66 ± 18.61 | 441.68 ± 13.82 | 456.86 ± 18.31 | 0.400 | 0.672 | 0.111 | 13.684 | **0.001** | 0.952 | 0.311 | 0.734 | 0.097 | - | - | - |
| Time in closed arc, day 2 | 2-way ANOVA | non normal | 448.33 ± 20.43 | 492.29 ± 16.58 | 487.59 ± 29.63 | 3.652 | **0.033** | 0.646 | 36.459 | **0.000** | 1.000 | 0.114 | 0.893 | 0.066 | 0.138 | **0.009** | 0.489 |
| Time in closed arc, mean | 2-way ANOVA | normal | 438.5 ± 18.27 | 466.99 ± 13.34 | 472.23 ± 20.15 | 1.917 | 0.158 | 0.379 | 28.873 | **0.000** | 1.000 | 0.253 | 0.778 | 0.088 | - | - | - |
| Time in open arc, day 1 | 2-way ANOVA | normal | 166.03 ± 18.25 | 153.09 ± 13.6 | 134.72 ± 17.99 | 0.562 | 0.574 | 0.138 | 10.417 | **0.002** | 0.886 | 0.420 | 0.659 | 0.114 | - | - | - |
| Time in open arc, day 2 | 2-way ANOVA | non normal | 143.08 ± 21.29 | 102.14 ± 16.74 | 78.39 ± 17.83 | 3.063 | **0.056** | 0.566 | 37.315 | **0.000** | 1.000 | 0.151 | 0.860 | 0.072 | 0.194 | **0.015** | 0.502 |
| Time in open arc, mean | 2-way ANOVA | normal | 154.55 ± 18.53 | 127.61 ± 13.25 | 106.56 ± 16.4 | 1.863 | 0.166 | 0.369 | 26.343 | **0.000** | 0.999 | 0.340 | 0.713 | 0.101 | - | - | - |
| Ratio time close/open, day 1 | 2-way ANOVA | non normal | 4.08 ± 0.85 | 3.74 ± 0.98 | 5.76 ± 1.6 | 0.820 | 0.447 | 0.182 | 9.172 | **0.004** | 0.843 | 0.438 | 0.648 | 0.117 | - | - | - |
| Ratio time close/open, day 2 | 2-way ANOVA | non normal | 6.95 ± 1.95 | 25.36 ± 18.5 | 28.98 ± 11.02 | 0.969 | 0.387 | 0.209 | 5.901 | **0.019** | 0.663 | 0.746 | 0.479 | 0.169 | - | - | - |
| Ratio time close/open, mean | 2-way ANOVA | non normal | 4.61 ± 1 | 5.55 ± 1.95 | 8.57 ± 2.42 | 1.207 | 0.308 | 0.251 | 11.378 | **0.001** | 0.911 | 0.538 | 0.587 | 0.134 | - | - | - |
| Number of open arc entries, day 1 | 2-way ANOVA | non normal | 48.47 ± 4.88 | 50.52 ± 4.46 | 59.36 ± 7.73 | 1.870 | 0.165 | 0.371 | 14.588 | **0.000** | 0.963 | 0.603 | 0.551 | 0.145 | - | - | - |
| Number of open arc entries, day 2 | 2-way ANOVA | normal | 42.42 ± 5.18 | 36.52 ± 4.83 | 29.31 ± 5.07 | 1.238 | 0.299 | 0.257 | 19.130 | **0.000** | 0.990 | 0.137 | 0.872 | 0.070 | - | - | - |
| Number of open arc entries, mean | 2-way ANOVA | normal | 45.44 ± 4.16 | 43.52 ± 4.04 | 44.34 ± 5.81 | 0.150 | 0.861 | 0.072 | 24.720 | **0.000** | 0.998 | 0.447 | 0.642 | 0.119 | - | - | - |
| Latency to enter in an open arc for the first time | 2-way ANOVA | non normal | 39.29 ± 13.16 | 47.08 ± 31.35 | 22.99 ± 10.38 | 0.310 | 0.735 | 0.097 | 0.684 | 0.412 | 0.128 | 1.282 | 0.287 | 0.265 | - | - | - |
| Latency to fully cross an open arc for the first time | 2-way ANOVA | non normal | 149.34 ± 34.87 | 139.69 ± 35.2 | 95.2 ± 32.99 | 0.796 | 0.457 | 0.178 | 1.515 | 0.224 | 0.226 | 0.395 | 0.676 | 0.110 | - | - | - |
| Dipping from close arc, frequency, day 1 | 2-way ANOVA | non normal | 51.73 ± 6.81 | 55.42 ± 6.48 | 51.47 ± 7.31 | 0.303 | 0.740 | 0.096 | 54.807 | **0.000** | 1.000 | 0.035 | 0.965 | 0.055 | - | - | - |
| Dipping from close arc, frequency, day 2 | 2-way ANOVA | non normal | 31.94 ± 3.01 | 28.94 ± 3.69 | 22.21 ± 3.87 | 2.182 | 0.124 | 0.425 | 54.920 | **0.000** | 1.000 | 0.593 | 0.556 | 0.143 | - | - | - |
| Dipping from close arc, frequency, mean | 2-way ANOVA | non normal | 41.84 ± 4.31 | 42.5 ± 4.48 | 36.84 ± 5.05 | 0.239 | 0.788 | 0.085 | 94.671 | **0.000** | 1.000 | 0.179 | 0.836 | 0.076 | - | - | - |
| Dipping from close arc, duration, day 1 | 2-way ANOVA | non normal | 131.15 ± 14.57 | 145.82 ± 14.24 | 117.88 ± 13.74 | 0.984 | 0.381 | 0.211 | 57.892 | **0.000** | 1.000 | 0.648 | 0.527 | 0.153 | - | - | - |
| Dipping from close arc, duration, day 2 | 2-way ANOVA | non normal | 94.51 ± 10 | 100.08 ± 13.98 | 64.33 ± 12.46 | 2.111 | 0.132 | 0.413 | 26.004 | **0.000** | 0.999 | 0.342 | 0.712 | 0.102 | - | - | - |
| Dipping from close arc, duration, mean | 2-way ANOVA | normal | 112.83 ± 11.57 | 124.49 ± 12.37 | 91.1 ± 10.18 | 2.386 | 0.103 | 0.459 | 72.828 | **0.000** | 1.000 | 0.811 | 0.450 | 0.181 | - | - | - |
| Dipping from open arc, frequency, day 1 | 2-way ANOVA | non normal | 27.15 ± 3.63 | 20.26 ± 2.46 | 19.15 ± 3.92 | 1.441 | 0.247 | 0.294 | 8.140 | **0.006** | 0.799 | 0.567 | 0.571 | 0.139 | - | - | - |
| Dipping from open arc, frequency, day 2 | 2-way ANOVA | non normal | 16.94 ± 3.04 | 12.22 ± 2.34 | 8.84 ± 2.54 | 1.938 | 0.155 | 0.383 | 17.131 | **0.000** | 0.982 | 0.332 | 0.719 | 0.100 | - | - | - |
| Dipping from open arc, frequency, mean | 2-way ANOVA | non normal | 22.05 ± 3.15 | 16.97 ± 2.15 | 14 ± 2.9 | 2.060 | 0.138 | 0.404 | 15.216 | **0.000** | 0.969 | 0.502 | 0.609 | 0.128 | - | - | - |
| Dipping from open arc, duration, day 1 | 2-way ANOVA | non normal | 62.85 ± 10.83 | 42.28 ± 5.15 | 25.57 ± 5.27 | 5.700 | **0.006** | 0.843 | 13.928 | **0.000** | 0.955 | 1.730 | 0.188 | 0.346 | **0.091** | **0.001** | 0.245 |
| Dipping from open arc, duration, day 2 | 2-way ANOVA | non normal | 51.3 ± 10.52 | 35.3 ± 8.44 | 17.87 ± 5.34 | 3.798 | **0.029** | 0.665 | 20.104 | **0.000** | 0.993 | 0.427 | 0.655 | 0.115 | 0.278 | **0.008** | 0.269 |
| Dipping from open arc, duration, mean | 2-way ANOVA | non normal | 57.07 ± 9.63 | 39.17 ± 5.86 | 21.72 ± 4.71 | 6.448 | **0.003** | 0.886 | 23.422 | **0.000** | 0.997 | 1.299 | 0.282 | 0.268 | **0.088** | **0.001** | 0.157 |

|  | test | data structure |  | All t | All p-value | power | WT t | WT p-value | power | Het t | Het p-value | power | KO t | KO p-value | power |
| --- | --- | --- | --- | --- | --- | --- | --- | --- | --- | --- | --- | --- | --- | --- | --- |
| Open vs close arc time, day 1 |  |  |  |  |  |  |  |  |  |  |  |  |  |  |  |
| - zone effect | repeated measures | sphericity assumed |  | 277.319 | **0.000** | 1.000 | 73.861 | **0.000** | 1.000 | 125.301 | **0.000** | 1.000 | 83.255 | **0.000** | 1.000 |
| - cohort effect | repeated measures | sphericity assumed |  | 8.156 | **0.006** | 0.935 | 5.563 | **0.031** | 0.604 | 8.257 | **0.011** | 0.773 | 3.298 | **0.087** | 0.403 |
| - zone x cohort effect | repeated measures | sphericity assumed |  | 12.518 | **0.001** | 0.801 | 7.248 | **0.015** | 0.718 | 3.063 | **0.098** | 0.379 | 2.143 | 0.161 | 0.282 |
| Open vs close arc time, day 2 |  |  |  |  |  |  |  |  |  |  |  |  |  |  |  |
| - zone effect | repeated measures | sphericity assumed |  | 440.281 | **0.000** | 1.000 | 94.767 | **0.000** | 1.000 | 278.317 | **0.000** | 1.000 | 119.843 | **0.000** | 1.000 |
| - cohort effect | repeated measures | sphericity assumed |  | 0.578 | 0.450 | 0.218 | 1.269 | 0.276 | 0.186 | 2.497 | 0.132 | 0.320 | 0.921 | 0.351 | 0.148 |
| - zone x cohort effect | repeated measures | sphericity assumed |  | 25.848 | **0.000** | 1.000 | 12.054 | **0.003** | 0.905 | 18.624 | **0.000** | 0.982 | 3.749 | **0.070** | 0.447 |
| Open vs close arc time, mean |  |  |  |  |  |  |  |  |  |  |  |  |  |  |  |
| - zone effect | repeated measures | sphericity assumed |  |  |  | 440.281 | 0.000 | 1.000 | 103.409 | 0.000 | 1.000 | 274.392 | 0.000 | 1.000 | 131.582 | 0.000 | 1.000 |
| - cohort effect | repeated measures | sphericity assumed |  |  |  | 0.578 | 0.450 | 0.116 | 0.006 | 0.941 | 0.051 | 0.422 | 0.524 | 0.094 | 0.484 | 0.496 | 0.101 |
| - zone x cohort effect | repeated measures | sphericity assumed |  |  |  | 25.848 | 0.000 | 0.999 | 11.720 | 0.003 | 0.897 | 12.720 | 0.002 | 0.919 | 3.786 | 0.068 | 0.451 |

No significant difference between the genotypes was observed in the open field thigmotaxis level (Figure 10A), but a decrease in the total number of times the mice reared (mainly driven by against wall rears) was observed in the *Shank3^Δ4-22^* homozygous animals (Figure 10B). No significant effects of an interaction between the time and genotype were observed for any of the parameters.

**Figure 10:**
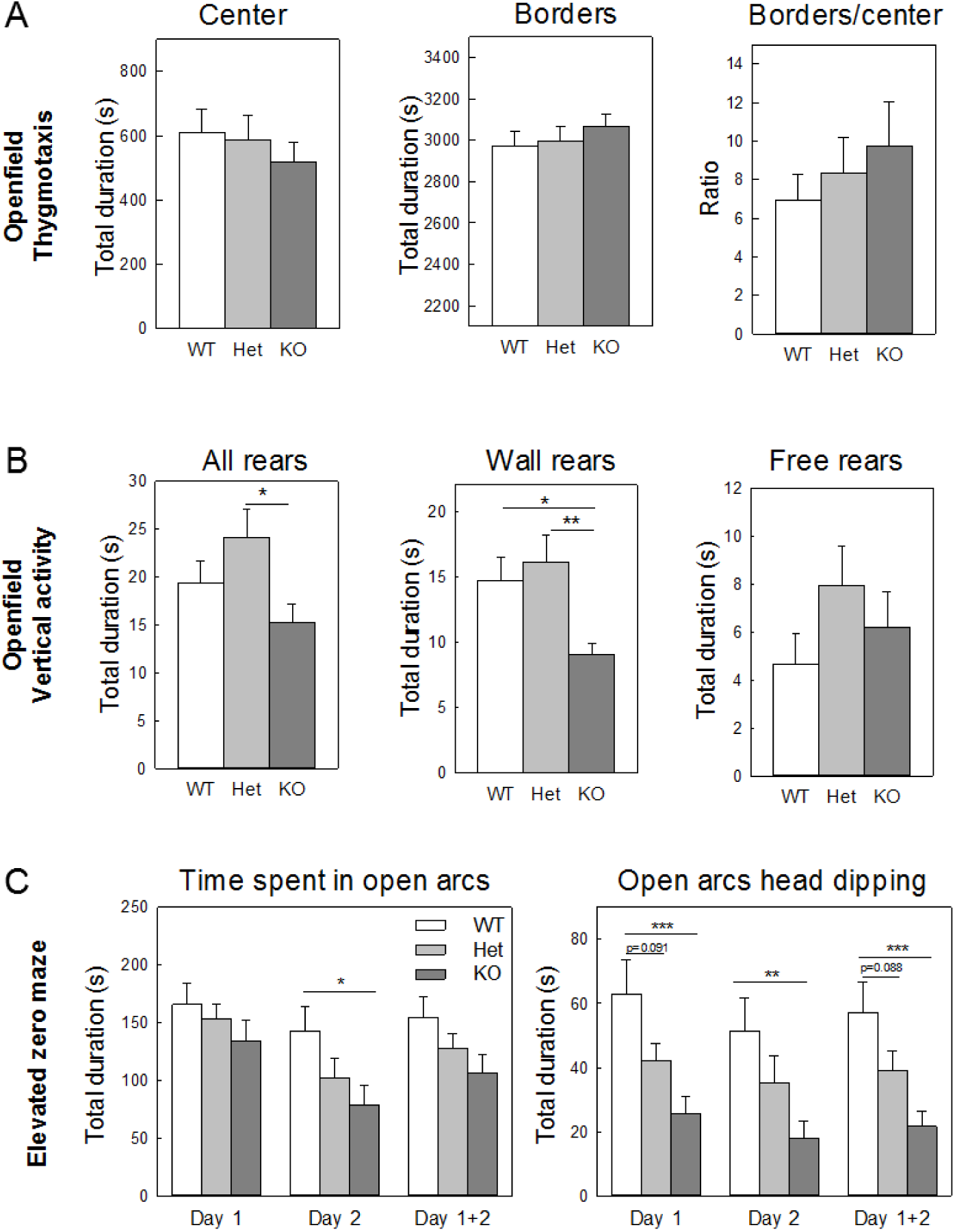
Anxiety-like behavior in *Shank3^Δ4-22^-*deficient mice. **(A)** Thigmotaxic behavior in open field. No genotype differences were found for the time spent in the center of the open field, the time spent close to the chamber walls (borders) or their ratio. **(B)** Vertical activity in open field. The cumulated time spend in free standing rears and rears against the walls of the open field were both counted. When compared to wild-type and heterozygotes littermates *Shank3^Δ4-22^* homozygous mice displayed decreased rearing activity due to a decrease of wall rears rather than free standing rears. **(C)** *Shank3^Δ4-22^* homozygous mice spent a lower amount of time in the open area when compared to wild-type and heterozygous mice. Similarly, the number of head dipping from the open arcs to the outside of the maze was reduced in *Shank3^Δ4-22^* homozygous mice. WT, wild-type mice; Het, heterozygous mice; KO, homozygous knockout mice. *: p<0.05, **: p<0.1, ***: p<0.001.

In the elevated zero-maze, all animals showed a preference for the closed arcs versus the open arcs, however *Shank3^Δ4-22^* homozygotes spent less time in the open arcs than their wild-type and heterozygous littermates. Similarly, a significant decrease of the duration of head dipping exploratory behavior in the open arcs was seen in those animals (Figure 10B). No genotype differences were seen for other parameters.

This indicates increases in anxiety in the *Shank3^Δ4-22^* homozygotes. In support of this, the long-lasting spontaneous freezing observed in *Shank3^Δ4-22^* homozygous animals during the habituation and before the sound-shock association in the fear conditioning training (Figure 9B) could also be explained by a higher anxiety level those animals.

## Discussion

Given the prevalence of complete *SHANK3* deletions in PMS, we generated *Shank3^Δ4–22^* mice by targeting exons 4-22, thereby disrupting all isoforms and providing improved construct validity compared to previously reported models. We conducted an extensive behavioral phenotyping of neonatal (P0-P21) and adult (3-8 months) mice to address both core symptoms and comorbidities observed in PMS. We confirmed our prediction that *Shank3*^Δ4–22^ mice homozygous and in some instances heterozygous mice have a more severe phenotype than previously published models with partial deletions of *Shank3* (summarized in Table 14). Our findings are consistent with recent results from an independent model also generated by disrupting all *Shank3* isoforms (Wang et al., 2016b).

**Table 14:**
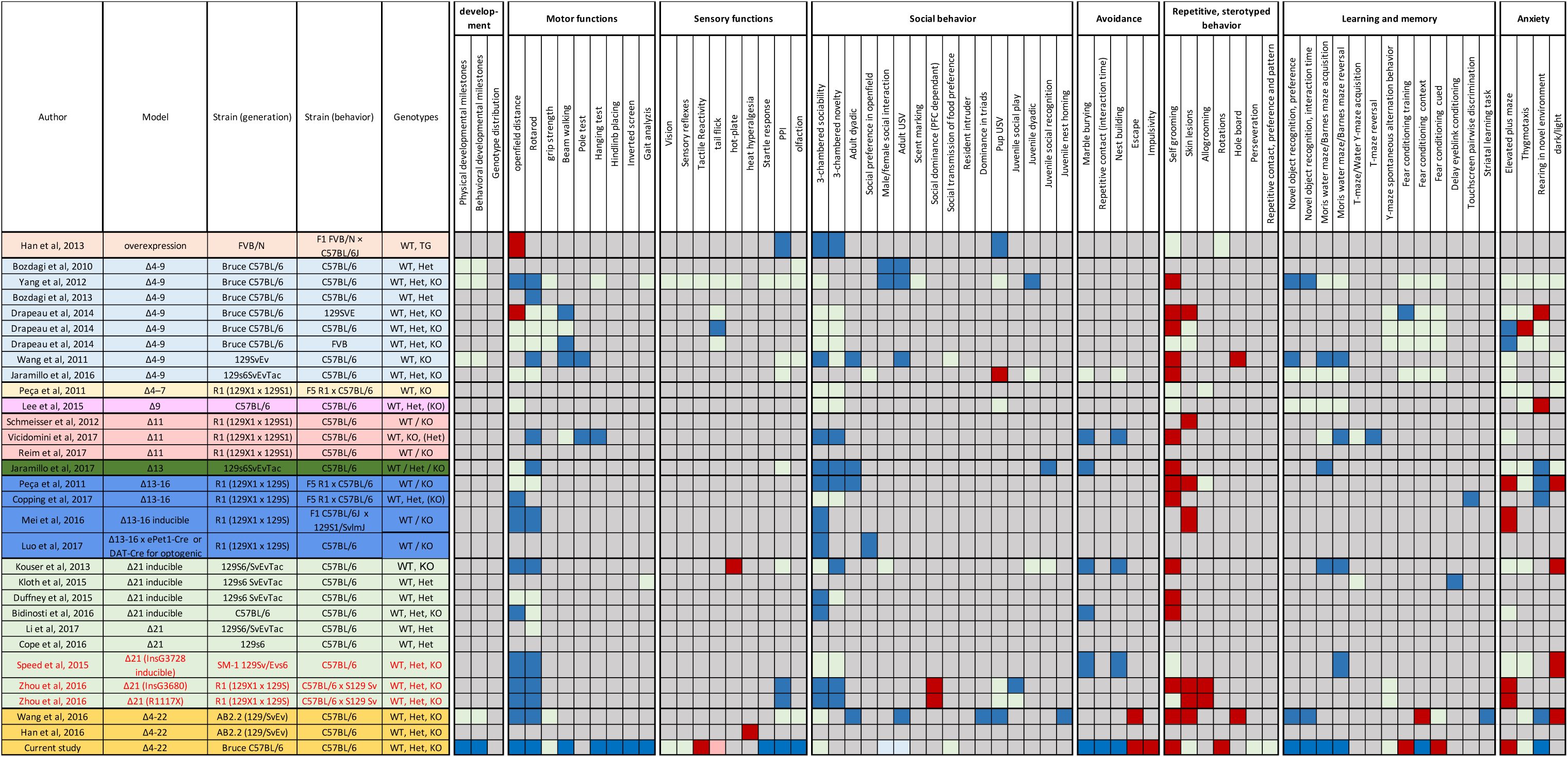
Main features and comorbidities associated with Phelan-McDermid displayed by different mouse models with *Shank3* deficits. Green indicates an absence of genotype difference. Blue indicates a decrease of the associated behavior in Shank3-deficient animals. Red indicates an increase of the associated behavior in Shank3-deficient animals. Grey indicates the behavior has not been studied in the corresponding article.

PMS is a neurodevelopmental disorder that manifests as early as in infancy by neonatal hypotonia and a generalized developmental delay. Previous studies have shown normal neonatal development in Δ4-9 mice (Bozdagi et al., 2010; Wang et al., 2011; Yang et al., 2012) or only minor delays limited to ear opening and paw positioning in Δ4-22 mice (Wang et al., 2016b). In the current study, both physical and behavioral developmental milestones were investigated. While physical delays were limited to a slower growth rate in *Shank3^Δ4–22^*-deficient animals, extensive sensory-motor deficits were observed. Some of them, such as the response to an auditory startle or the air righting ability, were only delayed while others, such as performances in the wire suspension tests and the grasping reflex, were still present at the time of weaning. Upon home-cage observation and physical examination of adult mice we did not observe severe deficits that would preclude advanced testing. Weight examined at three months of age was no longer different and survival curves between two and twenty-two months were not statistically different.

Hypotonia, motor-coordination impairments and gait abnormalities are a hallmark of PMS that persists beyond childhood (Phelan and McDermid, 2012; Soorya et al., 2013). In previous studies, motor performances have been frequently found to be impaired in adult *Shank3-*deficient mice (Table 14). Hence, decreased locomotion in the open field has been reported in many existing models including models with Δ4-9, Δ13-16, Δ21 deletions or point-mutations (Bidinosti et al., 2016; Copping et al., 2017; Kouser et al., 2013; Mei et al., 2016; Speed et al., 2015; Yang et al., 2012; Zhou et al., 2016) even if not always replicated in other models with similar or different deletions (Δ4-9, Δ9, Δ13, Δ13-16, Δ21 (Drapeau et al., 2014; Duffney et al., 2015; Jaramillo et al., 2017; Jaramillo et al., 2016; Lee et al., 2015; Peca et al., 2011)). Similarly, motor learning in accelerating rotarod was found to be impaired in Δ4-9, Δ11, Δ13, Δ13-16 and Δ21 models (Bozdagi et al., 2010; Jaramillo et al., 2017; Kouser et al., 2013; Mei et al., 2016; Speed et al., 2015; Vicidomini et al., 2016; Wang et al., 2011; Yang et al., 2012; Zhu et al., 2014) although not replicated in other studies (Δ4-9, Δ13-16 or Δ21 (Bidinosti et al., 2016; Drapeau et al., 2014; Duffney et al., 2015; Jaramillo et al., 2016; Li et al., 2017; Peca et al., 2011)). In agreement with Wang et al, both spontaneous locomotion and rotarod learning were strongly impaired in our *Shank3^Δ4–^ 22* mouse model. Interestingly, while most models only reported deficits in homozygous animals, heterozygous mice were also affected, albeit less severely. Difficulties in fine motor coordination have been described in Δ4-9 and Δ11 *Shank3-* deficient mice (Drapeau et al., 2014; Vicidomini et al., 2016; Wang et al., 2011) and were confirmed in the current study. In addition, our homozygous mice were strongly impaired in the hanging test, the hindlimb placing test and the inverted screen and had small gait abnormalities.

Hypersensitivity or hyposensitivity to sensory stimuli is frequently observed in PMS and ASD patients (Klintwall et al., 2011; Phelan and Betancur, 2011). However, little was known regarding the sensory abilities of *Shank3*-deficient mice. No deficits were reported in Δ4-9 or Δ4-22 animals for either olfaction, audition, vision, neuromuscular reflexes or pain sensitivity (Bozdagi et al., 2010; Wang et al., 2016b; Wang et al., 2011; Yang et al., 2012). Normal pre-pulse inhibition was observed in many models including Δ4-9, Δ13, Δ21 and Δ4-22 *Shank3*-deficient mice (Jaramillo et al., 2017; Kouser et al., 2013; Wang et al., 2016b; Yang et al., 2012) even if decreased pre-pulse inhibition was reported in in lines with point mutations in exon 21 (Zhou et al., 2016). Here, we observed that *Shank3^Δ4–22^* homozygous mice have no strong visual deficits, normal neuromuscular reflexes but are hyper-reactive in response to handling and tactile stimuli. In addition, we observed a delay in the acquisition of the startle response in newborns and a decrease of the startle response in both heterozygous and homozygous adults. Since social behavior strongly relies upon olfaction in rodents, we used different behavioral paradigms to evaluate our model. Interestingly, *Shank3^Δ4–22^* homozygous mice had a low interest for non-social olfactory stimuli as shown by deficits in the buried food test and by low amount of sniffing during the olfactory habituation/dishabituation paradigm. However, *Shank3^Δ4-22^-*deficient mice were able to discriminate odors in the test for social transmission of food preference or to show interest for social stimuli during olfactory habituation/dishabituation suggesting that they do not have anosmia but rather show reduced interest in non-social scents, which can be overcome when adding a social component.

One of the defining features of autism is the impairment of social interactions that can manifest by deficits in social approach, reciprocal social interactions and/or verbal and non-verbal communication. Mild social deficits have been reported, however with variability, in some of the previous studies of PMS mouse models (Table 14). In one of the most commonly used test, the three-chambered social approach test, no differences between the genotypes were reported in Δ4-9, Δ4-7, Δ9, Δ13-16, and Δ21 *Shank3*-deficient mice (Copping et al., 2017; Drapeau et al., 2014; Kouser et al., 2013; Lee et al., 2015; Peca et al., 2011; Speed et al., 2015; Yang et al., 2012) whereas a lack of preference for a social stimulus was observed other models targeting the same or different exons (Δ4-9, Δ11, Δ13, Δ13-16 and Δ21 (Bidinosti et al., 2016; Duffney et al., 2015; Jaramillo et al., 2017; Luo et al., 2017; Mei et al., 2016; Peca et al., 2011; Vicidomini et al., 2016; Wang et al., 2011; Zhou et al., 2016)). In freely interacting dyads, normal behavior was observed in most of the lines investigated (Δ4-9, Δ13, Δ13-16, Δ4-22 (Jaramillo et al., 2017; Peca et al., 2011; Wang et al., 2016b; Wang et al., 2011)) however reduction of sniffing time and ultrasonic vocalizations during adult male female social interaction were reported in some studies (Δ4-9, Δ21 and Δ4-22 (Bozdagi et al., 2010; Wang et al., 2016b; Wang et al., 2011; Yang et al., 2012)). Rodent social behavior is highly influenced by experimental conditions such as the animals’ age, housing conditions, or animals handling and that can explain differences observed between cohorts of animals with identical or similar alterations of the *Shank3* gene. Consistently Wang and colleagues’ study, we observed only minimal social deficits. All genotypes had a similar preference for social stimulus in the 3-chambered social approach test or the social transmission of food preference and only trends toward a decrease of interaction time and vocalization were found during male-female social interactions. While not representative of typical autism, this subtle behavior can reflect the phenotype of many patients with PMS. Indeed, unlike patients with idiopathic ASD, individuals with PMS show preserved responses to social communication cues (Soorya et al., 2013; Wang et al., 2016a) and roughly equal orienting to social versus non-social stimuli despite meeting criteria for ASD. Moreover, the fact that not all individuals with PMS are diagnosed with ASD indicates that animal models for PMS should not necessarily present with strong social behavioral deficits.

One of the strongest phenotype observed in the current study was an active avoidance of inanimate objects. In the novel object recognition test, lack of preference for a novel object had previously been observed in two lines of Δ4-9 mice (Wang et al., 2011; Yang et al., 2012) but not in a third line (Jaramillo et al., 2016) nor in Δ9 *Shank3*-deficient mice (Lee et al., 2015). However, in the present study homozygous animals had very little interactions with both familiar and novel object making it impossible to properly compare novelty preference. Instead they mostly spent their time in the corners of the open field away from the objects. Surprisingly, similar avoidance behavior was observed in the marble burying test and in the repetitive novel object contact task. We also observed a strong decrease of direct interactions with the applicator in the olfactory habituation/dishabituation test and a reduction of the quality of the nests build by *Shank3^Δ4–22^*-deficient animals with some mice even leaving the building material fully untouched. Some studies have reported that children with autism respond to novelty with avoidance behaviors and patients with PMS have enhanced reactivity to novel environments and very little interest for objects. Decrease of marble burying has been consistently been described in other models of *Shank3*-deficiency as were nest building impairments (Δ11, Δ13, Δ21 and exon 21 point mutations (Bidinosti et al., 2016; Jaramillo et al., 2017; Kouser et al., 2013; Speed et al., 2015; Vicidomini et al., 2016)). While we have shown that those animals are hypoactive and have significant motor deficits that could impact behavioral assays relying on exploratory locomotion, it is unlikely that this avoidance behavior is attributable to impaired motor activity or poor motivation as homozygous mice have normal pattern of investigation in an empty open field and actively avoid objects or even escape from the cages by jumping out while they will not escape from an empty cage or a cage containing an unfamiliar mouse. Furthermore, the number of escape attempts increased in relation with the number of objects present in the cage. In addition to this escape behavior, a high level of impulsivity was observed for adult homozygous mice in the beam walking test and for both newborn and adult homozygous mice in the negative geotaxis test.

Stereotypies, repetitive behaviors with restricted interests and resistance to change form the second set of core symptoms of ASD. Excessive grooming with or without development of skin lesions is the most commonly observed repetitive behavior in rodents. Repetitive/compulsive grooming has been reported in most of the previously published *Shank3* mouse models (Table 14) while skin lesions where noticed only in some of them (Δ4-9, Δ11, Δ13-16, Δ21 and point mutations in exon 21 (Drapeau et al., 2014; Mei et al., 2016; Peca et al., 2011; Schmeisser et al., 2012; Zhou et al., 2016)) suggesting different levels of severity. The homozygous mice from Wang et al. (2016) displayed both increased grooming and development of skin lesions. However, in the present study, even if we did occasionally observe some bald patches with or without skin lesions in our oldest animals all genotypes were concerned and group differences where only found for the grooming behavior. Our *Shank3^Δ4–22^*-deficient mice also engaged more frequently in other stereotyped and repetitive behaviors. By contrast, we did not observe any perseveration in the Y-maze nor object or pattern preference in the repetitive novel object contact task. To investigate both cognitive flexibility and insistence on sameness our animals were tested in the Barnes maze. The initial training showed a delay in the acquisition of the task in homozygous mice but after four days of training all genotypes had comparable performances and spent similar amount of time in the target quadrant during a probe test. Mice were then retrained after moving the escape box. Our homozygous mice exhibited impaired cognitive flexibility characterized by a delay in the time needed to learn the new rule and more especially by a similar preference for either the reversal target quadrant or the initial target quadrant during the probe test while heterozygous mice had an intermediate phenotype. This suggests that *Shank3* deficiency increases susceptibility to proactive interference, a deficit associated with prefrontal cortex dysfunction. Similar reversal impairments have been published in either the Morris water maze or T-maze in Δ4-9, Δ11, Δ21, point mutations or Δ4-22 mice (Kouser et al., 2013; Speed et al., 2015; Vicidomini et al., 2016; Wang et al., 2016b; Wang et al., 2011) while other models had comparable results for all genotypes (Δ4-9, Δ9 (Jaramillo et al., 2016; Lee et al., 2015; Yang et al., 2012)).

Because a majority of patients with *SHANK3* mutation/deletion exhibit some degree of intellectual disability, our animals were also tested for short-term memory by examining spontaneous alternation behavior in the Y-maze and for hippocampo-or amygdala-dependent memories using contextual and cued fear conditioning. As in other models investigated (Δ4-9 and point insertions (Drapeau et al., 2014; Zhou et al., 2016)) we found no differences in performance in the Y-maze spontaneous alternation test suggesting normal basic working memory. Neither contextual nor cued memories had been found to be affected by genotype in any of the previously published exon specific models (Δ4-9 (Drapeau et al., 2014; Jaramillo et al., 2016; Yang et al., 2012)) while a small increase of freezing was noticed in Δ4-22 homozygous mice during contextual recall (Wang et al., 2016b). Interestingly, in our new mouse model, we observed distinct responses to each phase of the testing. While not different at first during the pre-training habituation phase, the level of freezing quickly decreased in wild-type and heterozygous mice but not in the homozygous animals likely reflecting a higher anxiety level. Upon presentation of the sound/shock associations, the increase of freezing was significantly more important in homozygous mice. Remarkably, the opposite was observed during the contextual recall thus demonstrating an impairment of hippocampo-dependent in homozygous animals while the same mice displayed increased freezing upon the presentation of sounds during the amygdala-dependent cued recall. Specific subcategories of learning and memory behaviors have only been studied in limited number of previous models. Δ13-16 *Shank3*-deficient mice are impaired in pairwise visual discrimination learning in the automated touchscreen task (Copping et al., 2017), heterozygous Δ21 mice exhibit impaired eye-blink conditioning, a cerebellar-dependent learning task (Kloth et al., 2015) and Δ4-22 homozygous mice have deficits in a striatal-dependent instrumental learning task (Wang et al., 2016b).

Altogether, the hyper-reactivity to handling and tactile stimuli, the impulsivity, the object neophobia, the escape behavior, the increased freezing response in the pre-training phase of the fear conditioning and in cued retrieval suggest high levels of anxiety in our mouse model. Hyperactivity and anxiety are other common features of PMS (Dhar et al., 2010; Sarasua et al., 2014a; Soorya et al., 2013). In previously published models, analysis of anxiety-like behavior measured either elevated mazes, in the open fields or in dark/light emergence boxes have demonstrated a relationship between the targeted isoforms and the manifestations of anxiety like-behavior. While little differences were observed in mouse models with Δ4-9, Δ4-7, Δ9 and Δ11 deletions (Drapeau et al., 2014; Jaramillo et al., 2016; Lee et al., 2015; Peca et al., 2011; Reim et al., 2017; Schmeisser et al., 2012; Vicidomini et al., 2016; Wang et al., 2011; Yang et al., 2012) increased levels of anxiety were reported in mice with Δ13, Δ13-16 and Δ21 deletions or point mutations (Copping et al., 2017; Jaramillo et al., 2017; Kouser et al., 2013; Mei et al., 2016; Peca et al., 2011; Speed et al., 2015; Zhou et al., 2016). Increased level of anxiety was confirmed in the light-dark emergence test and decrease and in the open field in the in Δ4-22 mouse model from Wang and colleagues or in the elevated maze and in the open field in our model.

In conclusion, our complete *Shank3^Δ4–22^* mouse line provides a new and improved genetic model for studying mechanisms underlying ASD and PMS and is characterized both by better construct and face validities than previously reported lines of *Shank3* mutants. Our in-depth behavioral characterization revealed behavioral features that reflect those observed in PMS and therefore suggest a greater potential as a translational model. Mice with a complete deletion of *Shank3* are more severely affected than previously published mouse models with a partial deletion. Both sensory and motor disabilities were detected in neonate and adult mice. *Shank3^Δ4–22^*-deficient mice showed modest deficits in social behavior, reflected in reduced male to female anogenital sniffing and ultrasonic vocalization, but no major deficits in social preference in the 3-chambered social interaction task. These findings are consistent with an independently generated mouse model (Wang et al., 2016b). Also in agreement with Wang’s study, our *Shank3*^Δ4–22^ mice showed increased anxiety and hyper-reactivity to novel stimuli, increased escape behaviors, and increased repetitive behaviors. Together with the increased freezing behavior in the cued fear conditioning, this suggest a dysregulation of amygdala circuitry that will require further investigation. In addition, our mice displayed impairments in several hippocampo-dependant learning and memory tests as well as cognitive inflexibility thus recapitulating intellectual disability and insistence on sameness observed in the majority of patients with PMS. Although PMS patients are heterozygous for *Shank3* mutations/deletions, most of the previous models have failed to demonstrate any relevant phenotype in heterozygous animals. Here, we were able to observe an intermediate phenotype for heterozygous mice in several of the parameters tested, notably in the open field, rotarod, startle response, escape behavior, reversal probe test and elevated zero-maze. Heterozygous animals being less affected than their homozygous, we hypothesis that more challenging paradigms, for example by introducing a variable reward probability in tests such as the Barnes maze would allow us too furthermore highlight differences in heterozygous animals. Past studies have often failed to replicate behavioral phenotype even in models with very similar *shank3* disruption or in different cohorts from the same model. The concordant findings from two independently derived and analyzed *Shank3* mouse models, including the comparison of two independent cohorts in our laboratory, demonstrate, for the first time, strong reproducibility and validity for a genetically modified mouse model of PMS providing a valuable model for further investigations of the neurophysiological basis of PMS and ASD.

**Author contributions**
J.D.B. and E.D. designed the experiments. E.D. and M.R. performed the behavioral experiments. Y.K. performed the western blot and PCR analysis. E.D. analyzed data. The manuscript was written by E.D. and all authors reviewed the manuscript before submission.

## Acknowledgements

We are indebted to Jacqueline Crawley for her help all along this study and her precious comments on the manuscript and to Jill Silverman for reviewing our results. We thank Dr. Nikolaos Daskalakis for all our helpful discussions and his help with data analysis.

